# Photosynthetic demands on translational machinery drive retention of redundant tRNA metabolism in plant organelles

**DOI:** 10.1101/2023.08.01.551541

**Authors:** Rachael Ann DeTar, Joanna Chustecki, Anna Martinez-Hottovy, Luis Federico Ceriotti, Amanda K. Broz, M. Virginia Sanchez-Puerta, Christian Elowsky, Alan C. Christensen, Daniel B. Sloan

## Abstract

Eukaryotic nuclear genomes often encode distinct sets of translation machinery for function in the cytosol vs. organelles (mitochondria and plastids). This raises questions about why multiple translation systems are maintained even though they are capable of comparable functions and whether they evolve differently depending on the compartment where they operate. These questions are particularly interesting in plants because translation machinery, including aminoacyl-tRNA synthetases (aaRS), is often dual-targeted to the plastids and mitochondria. These organelles have different functions, with much higher rates of translation in plastids to supply the abundant, rapid-turnover proteins required for photosynthesis. Previous studies have indicated that plant organellar aaRS evolve more slowly compared to mitochondrial aaRS in eukaryotes that lack plastids. Thus, we investigated the evolution of nuclear-encoded organellar and cytosolic aaRS and tRNA maturation enzymes across a broad sampling of angiosperms, including non-photosynthetic (heterotrophic) plant species with reduced plastid gene expression, to test the hypothesis that translational demands associated with photosynthesis constrain the evolution of enzymes involved in organellar tRNA metabolism. Remarkably, heterotrophic plants exhibited wholesale loss of many organelle-targeted aaRS and other enzymes, even though translation still occurs in their mitochondria and plastids. These losses were often accompanied by apparent retargeting of cytosolic enzymes and tRNAs to the organelles, sometimes preserving aaRS-tRNA charging relationships but other times creating surprising mismatches between cytosolic aaRS and mitochondrial tRNA substrates. Our findings indicate that the presence of a photosynthetic plastid drives the retention of specialized systems for organellar tRNA metabolism.

**Significance:** The process by which endosymbionts are integrated into a host and become organelles results in a combination of gene loss, transfer to the nucleus, and retention in the organellar genome. It is not clear why some endosymbiont-derived genes may be retained when a functional host counterpart exists whose gene product could simply be retargeted to the organelles. This study revealed that the photosynthetic activity in plant plastids may be responsible for retention of functionally redundant tRNA processing machinery, while mitochondria are more flexible regarding substitution with cytosolic-type enzymes. Therefore, functional constraint in the plastid is likely more important than in the mitochondria for shaping the evolution and retention of functionally redundant proteins that are dual targeted to both organelles.

## Introduction

The evolutionary transition from a free-living bacterium to an endosymbiont and eventually to an organelle requires the physical, biochemical, and genomic integration of two organisms. One common consequence of this process is a reduction in the genome of the endosymbiont, driven by relocation of essential genes to the nucleus via horizontal transfer or outright loss of some genes that are no longer useful after transition to an intracellular lifestyle. This trajectory has been observed repeatedly for various endosymbionts and organelles, including mitochondria, plastids, and more recently acquired intracellular bacteria such as the cyanobacterial-derived chromatophores of *Paulinella* amoeba or the many heritable endosymbionts in insects (1–6). However, there are some endosymbiotically derived genes that are recalcitrant to loss despite the presence of host counterparts with comparable functions. Redundant genes may be maintained in the organelle genomes or in the nuclear genome alongside genes encoding proteins for the same role in other cellular compartments. These observations elicit some interesting questions; 1) Why are some functionally redundant genes retained? 2) Do these redundant genes evolve differently depending on the cellular compartment where they operate?

The genes that encode the machinery required for organelle translation are widely retained, as eukaryotes maintain distinct translation systems for nuclear and mitochondrial transcripts. Translation becomes even more complicated in eukaryotes bearing three genomes. For example, plant cells contain nuclear, mitochondrial, and plastid genomes. The translation machinery associated with each of these cellular compartments includes ribosomal RNAs (rRNAs), ribosomal proteins, transfer RNAs (tRNAs), and the accompanying enzymes involved in tRNA processing and aminoacylation. The aminoacyl-tRNA synthetases (aaRS) that catalyze the “charging” of a tRNA with its cognate amino acid tend to be highly conserved, with distinct versions for function in the organelles and cytosol (7). The degree of redundancy of tRNAs and their interactors between cellular compartments also varies significantly within eukaryotes. These characteristics make tRNAs and their interactors, particularly aaRS, an ideal case study for understanding cytonuclear gene redundancy.

The concerted activity of the aaRS along with other enzymes involved in post-transcriptional modification such as tRNases P and Z (end processing in tRNA maturation) and CCAses (addition of CCA tail) ensures the tRNA has the proper structure to fulfill its role in translation. Generally, organisms would need a complement of at least 20 aaRS, one for each amino acid. However, eukaryotes often have more than this minimum requirement, with some overlapping sets of enzymes targeted to different compartments (8, 9). It is not fully understood why some eukaryotes retain distinct, redundant sets of tRNA processing machinery, while others do not. However, retention of organelle-targeted aaRS correlates with the retention of tRNA genes within organelle genomes. For example, *Homo sapiens* retains 38 aaRS with nearly distinct sets for the cytosol and mitochondria (9–11). Correspondingly, all necessary tRNA genes are present in the mitochondrial genome (mitogenome) (8). In contrast, trypanosomatids do not encode any tRNAs in their mitogenomes but rather import nuclear-encoded tRNAs into the mitochondria and have a reduced set of aaRS that colocalize to the cytosol and the mitochondria (9, 12, 13). In plants, closely related species may have very different numbers of tRNAs encoded in their mitogenomes, with import of cytosolic tRNAs to compensate for losses (8, 14, 15). Notably, plants also dual-target and import an organellar set of enzymes into both plastids and mitochondria (9, 16). The co-occurrence of redundant tRNAs and aaRS could be explained by the observation that organelle-encoded tRNAs are often structurally divergent from cytosolic tRNAs and require specific enzymatic partners for recognition (8, 17).

For most eukaryotes, organellar and cytosolic aaRS are exclusively nuclear-encoded. The shared genomic location of these genes facilitates direct comparison of sequence conservation because any differences in evolutionary rate more likely reflect differences in selection, rather than other factors like compartment-specific mutation rates or effective population size (18–21). Previous studies have quantified rates of evolution for these enzymes in various animal lineages and a small number of plants. Mitochondria-specific aaRS in animals exhibit lower sequence conservation than their cytosolic counterparts (22–24). In contrast, plant organellar aaRS exhibit equivalent or greater sequence conservation compared to their cytosolic counterparts (25). Furthermore, plastid genomes (plastomes) tend to retain a set of 30 tRNA genes that is fully sufficient for translation (26, 27). Plastomes also generally have high rates of transcription (28) and translation (29) due to the abundance and rapid turnover of proteins required for photosynthesis. Thus, the disparity in evolutionary rates between animal mitochondrial aaRS and plant dual-organelle targeted aaRS may be due to the specific translational needs of the plastid. We hypothesize that aaRS and other translation-associated enzymes operating in the plastid face great functional constraints, thus promoting unusually high sequence conservation of organellar translation machinery compared to other eukaryotes. A prediction stemming from this hypothesis is that plant lineages with reduced or nonexistent plastid gene expression would exhibit more rapid evolution of organellar aaRS due to relaxed selection.

In this work, we test whether photosynthetic activity contributes to sequence conservation of organellar tRNA processing and charging enzymes by analyzing evolution of this machinery across diverse lineages of autotrophic and heterotrophic plants. It should be noted there is a gradient of heterotrophy amongst plants – for the purposes of this study, we use “heterotrophic” to describe plants that do not photosynthesize at all and exclusively rely on parasitism for acquisition of carbon. They do this by tapping directly into host plants or by exploiting mycorrhizal fungi (mycoheterotrophy) (30, 31). We refer to facultative autotrophs that supplement photosynthetic carbon with parasitism as “hemiparasitic.” The hallmark of many heterotrophic/hemiparasitic plants is a reduced or even non-existent plastome (32). Highly expressed genes related to photosynthesis such as the Rubisco large subunit and the Photosystem II D1 reaction center tend to be absent from plastomes in heterotrophs, thereby drastically decreasing demands on plastid gene expression. In addition, many tRNA genes have been lost from the plastomes of these plant species (32), providing a good system to examine the relationship between evolutionary redundancy of aaRS, tRNA content of the organelle genomes, and trophic lifestyle.

## Results

### Heterotrophic plants exhibit extensive loss of organellar aaRS and tRNA processing enzymes but not ribosomal protein subunits

We searched for organellar and cytosolic aaRS orthologs in a broad sampling of angiosperm species with a range of trophic lifestyles and plastome reduction (**Supp. Dataset 1**). For each hemiparasitic/heterotrophic species in the analysis, we included the closest fully autotrophic relative with a nuclear genome or transcriptome available, excepting the Balanophoraceae where we used hemiparasite *Santalum album* as the reference autotroph. In total, we sampled 7 heterotrophs, 8 hemiparasites, and 10 autotrophs, including reference species *Arabidopsis thaliana.* Among heterotrophs, we included the endoparasites *Rafflesia cantleyi* and *Sapria himalayana* from the family Rafflesiaceae, likely the most extreme examples of plant heterotrophy, as several studies have suggested wholesale loss of the plastome in these species (33–36).

We found orthologs across these diverse taxa by running Orthofinder (37, 38) on protein models derived from nuclear genome or transcriptome data to find groups of likely orthologous proteins based on similarity and phylogeny, i.e., “orthogroups”. Then we extracted sequences that fall into an orthogroup with known *A. thaliana* aaRS (**Supp. Dataset 2**). It is important to note that the number of homologous sequences found for each species in this analysis may not directly reflect the absolute number of genes or proteins due to the fragmented nature of gene models made from transcriptome data and the inclusion of two independent sets of protein models for *Balanophora fungosa* and *S. himalayana.* Unexpectedly, many lineages of heterotrophs did not have any orthologs for many of the organellar aaRS. This loss of organelle-specific orthologs in heterotrophs was surprising as even species entirely lacking the plastome must still conduct tRNA aminoacylation within mitochondria. Further investigation using reciprocal-best-hit (RBH) searches with BLAST between *A. thaliana* and each species supported that these were true losses. These losses primarily occurred in purely heterotrophic plants, while hemiparasites retained most of their organellar aaRS. (**Fig. 1**). Yet, within the heterotrophs, there was a gradient of loss. The two endophytic species from Rafflesiaceae (*R. cantleyi* and *S. himalayana*) exhibited the most striking loss of organellar aaRS, retaining only PheRS. In contrast, obligate mycoheterotroph *Monotropa hypopitys* retained more aaRS than any other heterotroph, missing only AlaRS, CysRS, ThrRS, and TrpRS (**Fig. 1**).

**Figure 1:**
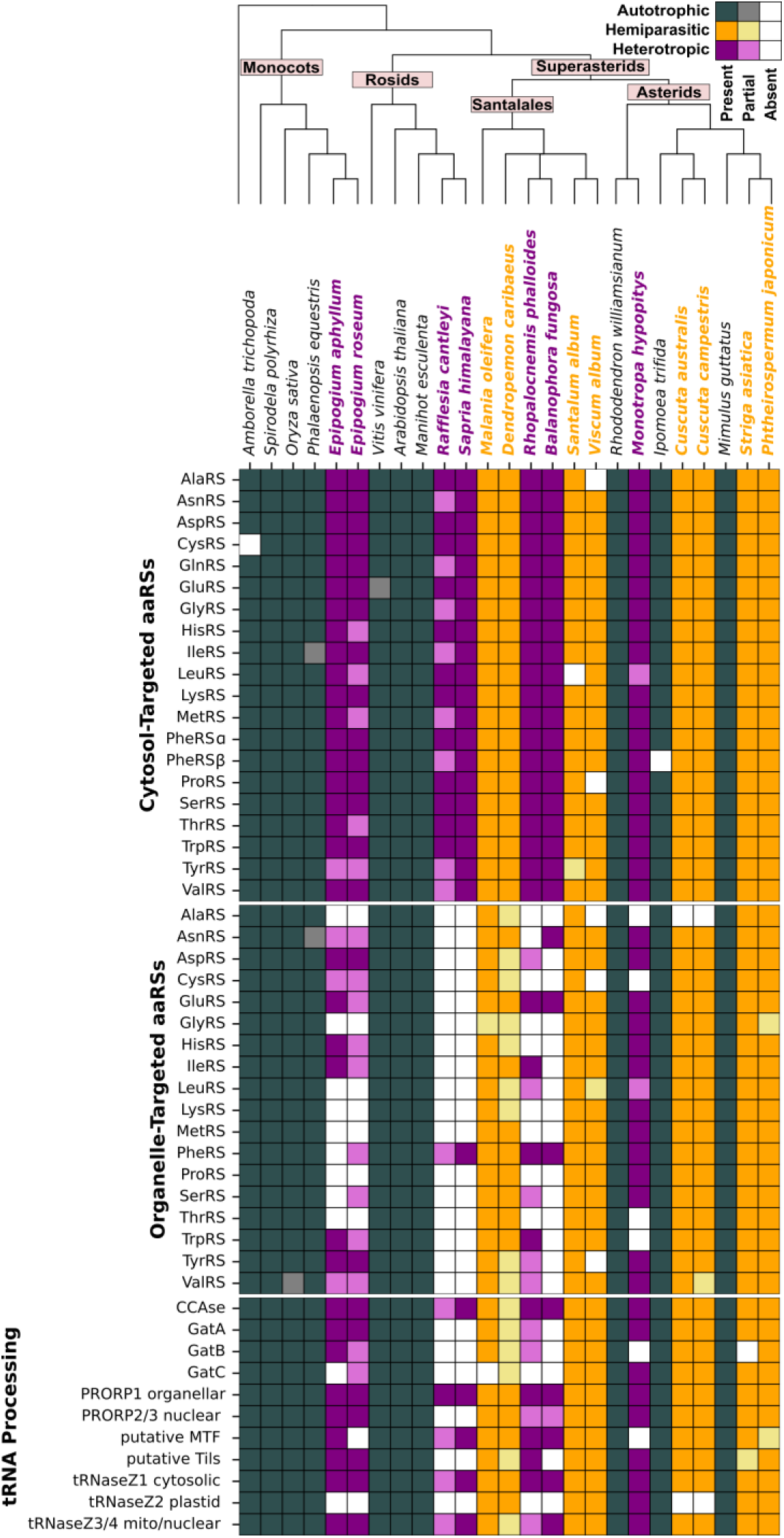
Presence/absence matrix of selected organellar and cytosolic aaRS and tRNA-processing enzymes shows major losses of organellar enzymes in heterotrophic lineages. Hue of boxes indicates metabolism (gray = autotroph, orange = hemiparasite, purple = heterotroph), shade indicates degree of presence/absence (white = no sequence found, light colors = partial sequence found, dark colors = complete or near complete sequence found). Note that there is no organellar GlnRS as plants typically use the bacterial-type GatCAB amidotransferase system to charge organellar Gln-tRNA. ArgRS is not presented as it has no distinct organellar enzyme; gene products are either cytosol-targeted, or triple-targeted to the cytosol, mitochondria, and plastids.

This striking pattern led us to investigate the retention of other enzymes involved in tRNA maturation and processing; see (15) and **Supplemental Dataset 2** for a comprehensive list. Unsurprisingly, the CCAse enzyme responsible for adding CCA tails to tRNAs in all compartments was retained across all species. Two enzymes that maintain distinct cytosolic and organellar versions, tRNase Z (involved in 3′ end processing) and Protein Only RNase P (PRORP; required for 5′ end processing), exhibited two different trends. We found that plastid tRNase Z (tRNase Z2) (39) was missing from all heterotrophs except *M. hypopitys* and hemiparasites of the *Cuscuta* genus yet was retained in the autotrophic relatives. Conversely, the organellar PRORP1 (40, 41) appeared to be universally retained while cytosolic PRORP2/3 orthologs were missing from the Rafflesiaceae. We also investigated the retention of bacterial-like organelle-specific machinery with no cytosolic counterpart, including the methionyl-tRNA formyltransferase (MTF), tRNA-Ile lysidine synthetase (TilS), and the bacterial-type glutamine amidotransferase (GatCAB) complex. We found orthologs for the putative *A. thaliana* MTF in all species except the heterotrophs *Epipogium roseum* and *M. hypopitys,* and putative TilS orthologs were absent only from the heterotrophs Rafflesiaceae and *B. fungosa*. GatCAB is the key complex used by organelles in land plants to accurately charge tRNA^Gln^ in the absence of a dedicated organellar GlnRS (42–44). This involves a two-step process, whereby tRNA^Gln^ is initially mis-acylated with Glu by GluRS. GatCAB then catalyzes the transamidation of Glu to Gln. Most of our test species, even heterotrophs, retained genes for at least some of the GatCAB subunits. However, Rafflesiaceae species appear to have lost all subunits in this complex.

To assess whether other functionally redundant translation machinery was lost in heterotrophic lineages, we analyzed presence/absence of nuclear-encoded riboproteins with exclusive targeting to a given compartment (Table S1, Supp. Dataset 2, “Riboproteins”). While there was minor loss of cytosolic, mitochondrial, and plastid subunits in all heterotrophic lines, the Rafflesiaceae lineages exhibited a dramatic reduction in plastid subunits, which is expected because these lineages appear to have dispensed with the plastid genome entirely and consequently no longer need plastid ribosomes. This finding reveals a key contrast between the fate of organellar aaRS and riboproteins in response to loss of photosynthesis and may reflect the fact that aaRS are shared between the two organelles, whereas the riboproteins are compartment-specific.

Generally, we observed reductive evolution in all heterotrophs via loss of organellar aaRS and bacterial-type enzymes for tRNA modification. This raises the question as to how gene expression is accomplished in the organelles of these species. One possibility is that cytosolic enzymes are retargeted to the organelles to functionally replace organellar enzymes.

### Heterotrophic plants may compensate for loss of organellar aaRS by retargeting cytosolic counterparts

To test for cytosolic aaRS retargeting, we applied two *in silico* targeting prediction programs to protein models for all cytosolic aaRS in our heterotrophs and their close autotrophic relatives (**Fig. 2**, **Fig. S1**). Our analysis revealed that canonically cytosolic aaRS are disproportionately retargeted to mitochondria in obligate heterotrophs compared to autotrophs (*p* = 0.018). There was also a non-significant trend towards retargeting to the plastid (*p* = 0.11). Targeting of mitochondrial OXPHOS components and plastid housekeeping proteins including subunits of the caseinolytic protease (Clp) complex and the acetyl-CoA carboxylase (ACC) complex involved in producing malonyl-CoA for fatty acid biosynthesis were included as known organellar positive controls (Supp. Dataset 2, “targeting controls”). Protein degradation and fatty acid biosynthesis are essential plastid functions that are still necessary in the absence of photosynthesis. Cell wall synthesis enzymes were included as negative controls. Of the controls, the only group that showed a different degree of targeting than expected was the mitochondrial OXPHOS components, where recovery of proteins with known mitochondrial targeting was low (∼60%) for both autotrophs and heterotrophs (Supp. Table 2). Thus, these *in silico* approaches appear to successfully (albeit imperfectly) predict subcellular localization in both heterotrophic and autotrophic plants.

**Figure 2:**
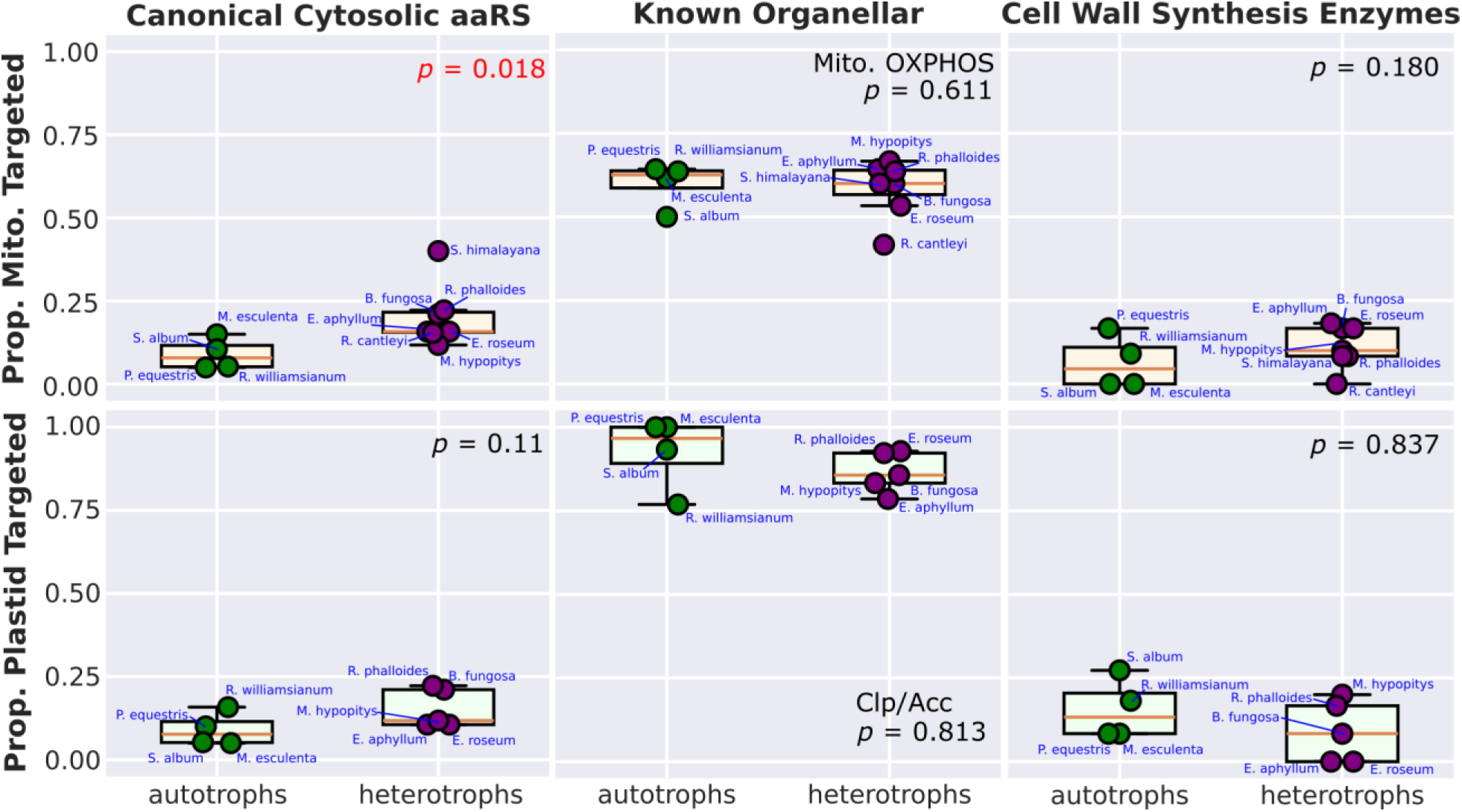
Proportion of retargeted cytosolic aaRS is higher in heterotrophs than autotrophs based on *in-silico* predictions. Proportion of enzymes with predicted retargeting (probability ≥ 0.5) to either mitochondria (top row) or plastids (bottom row) based on the outputs of LOCALIZER and/or TargetP. Autotrophs and heterotrophs are plotted in green and purple, respectively. Targeting predictions for proteins involved in mitochondrial oxidative phosphorylation (OXPHOS) are included as a positive control for mitochondrial targeting, and proteins involved in non-photosynthesis related housekeeping roles (ACC and Clp subunits) are included as a positive control for plastid targeting. Predictions for cell-wall synthesis associated enzymes are included as a negative control for organelle targeting. Reported *p*-values from *t*-tests or Mann-Whitney U tests depending on distribution of the data. Note that *R. cantleyi* and *S. himalayana* are not included in plastid retargeting analysis due to suspected loss of the plastome, but they were predicted to have retargeted 7.7% and 30.0% of their cytosolic aaRS to the plastid, respectively. These species have lost too many plastid housekeeping genes to assess proportion targeted. For both species, no cell wall synthesis enzymes exhibited plastid retargeting.

We would specifically expect that a given cytosolic aaRS is more likely to be retargeted if its organellar counterpart is lost. We examined this possibility and found a clear and statistically significant correlation between loss of an organellar aaRS and retargeting of the corresponding cytosolic enzyme to one or both compartments (*p* = 0.005, **Fig. 4**, **Fig. S2**).

The variable quality of gene models for each species and high false negative rate for retargeting prediction confound interpretation of retargeting for individual enzymes in each species (**Fig. 4**, **Fig. S3**). However, when looking only at *S. himalayana*, a species with relatively high-quality genome-based models, we observed some interesting phenomena. Cytosolic AlaRS, GlnRS, MetRS, ProRS, and TrpRS were predicted to be retargeted to mitochondria. Retargeting of cytosolic GlnRS could explain how this species copes with loss of the GatCAB complex (**Fig. 1**). The cytosolic AspRS, GluRS, and GlyRS were predicted to be dual targeted to both organelles, while cytosolic TyrRS and the PheRSα subunit were predicted to be targeted to the plastid despite the likely absence of a plastid genome. All other cytosolic enzymes were either not predicted to be retargeted or were missing a large portion of the N-terminus. The fact that some cytosolic aaRS corresponding to lost organellar aaRS were not predicted to be retargeted to the mitochondria and that some cytosolic aaRS were targeted to a plastid presumed to lack gene expression may reflect an underestimation of mitochondrial retargeting due to the constraints of *in silico* predictions and the potential divergence of TP sequences in Rafflesiaceae. Indeed, an investigation of subunits for the protein import apparatus of plastids and mitochondria revealed that the Rafflesiaceae are missing some key components of the plastid import apparatus (Fig. S6), some of which are responsible for TP binding and recognition.

To assess whether our *in silico* predictions hold true *in vivo,* we conducted targeting assays on a subset of aaRS (see **Supplemental Dataset 3** for sequence information) by inducing transient expression of N-terminal fusions of *S. himalayana* cytosolic aaRS transit peptides (TPs) to green fluorescent protein (GFP) in *Nicotiana benthamiana* (**Fig. 3**, **Fig. S4-S5**). We used eqFP611 tagged with the TP from mitochondrial isovaleryl-CoA dehydrogenase (IVD) as a positive control for mitochondrial targeting. Infiltration with single expression vector with just IVD_FP611 was used as a negative control (**Fig. S4**). We found that proteins predicted to be retargeted *in silico* were almost universally retargeted to one or both organelles *in vivo*. In some cases, putative TPs retargeted GFP to a different organelle than predicted *in silico.* AsnRS displayed the plastid targeting that was predicted *in silico* and additional mitochondrial targeting *in vivo* (**Fig. 3**). GluRS and GlyRS were predicted to be dual-targeted but exhibited targeting to only one of the two organelles. The only two TPs for which organellar GFP localization completely mismatched the *in silico* prediction (but still exhibited organelle targeting) were ProRS and TrpRS, which both showed strong plastid GFP fluorescence *in vivo* despite mitochondrial predictions. These results were consistently observed with two different cloning and expression strategies (Fig. 3, Fig. S5). The cytosolic AspRS TP was the only one we cloned that did not exhibit targeting to either organelle *in vivo* despite a clear biological expectation that it would have evolved retargeting (**Fig. S4**). GFP tagged with this TP formed punctate structures in the cytosol, which may be artefactual aggregates resulting from overexpression. Generally, *in vivo* assays provided evidence for retargeting of cytosolic aaRS to at least one organelle and suggested that *in silico* prediction tools may not be adept at distinguishing plastid from mitochondrial targeting.

**Figure 3:**
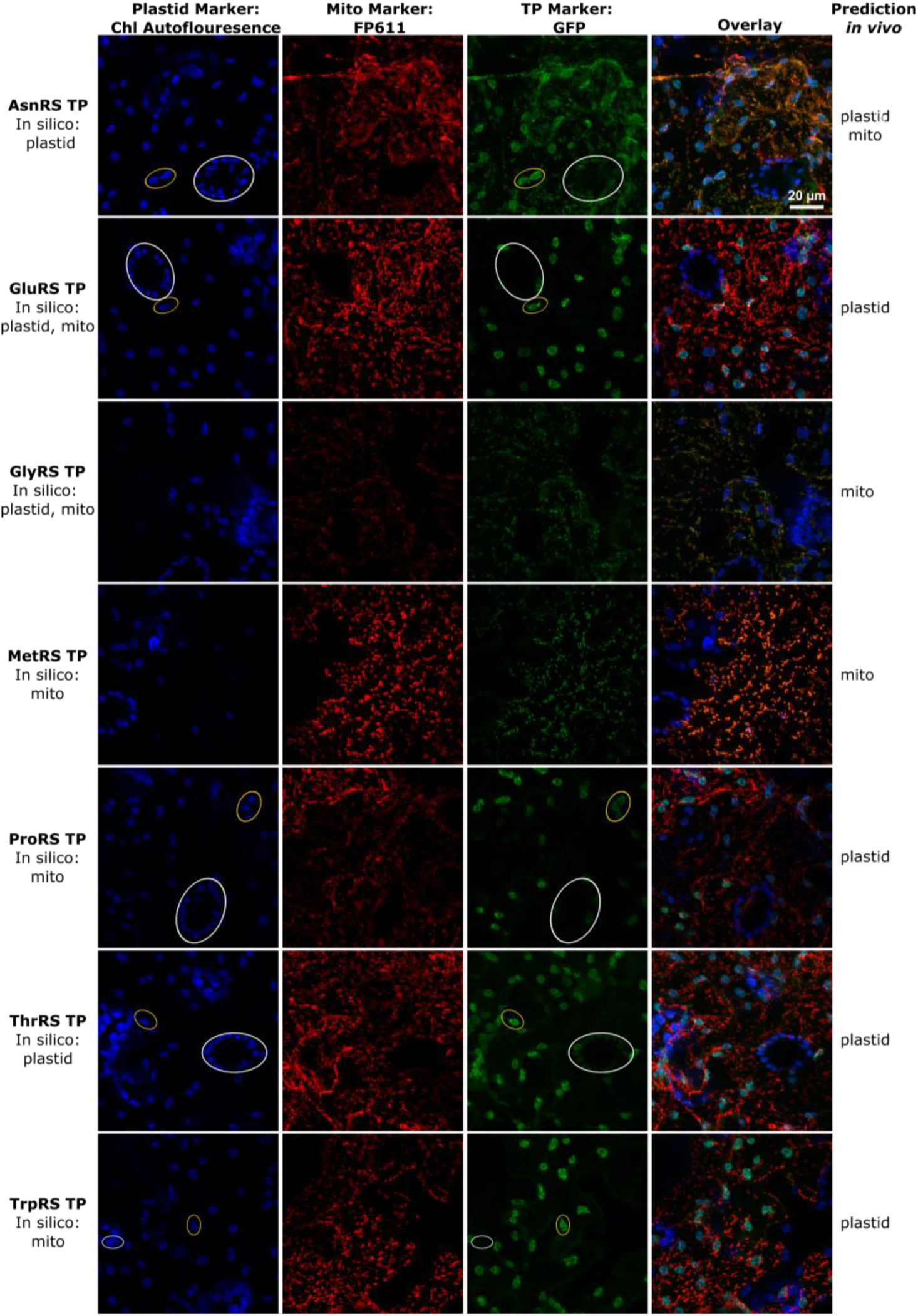
Cytosolic-type aaRS transit peptides from heterotroph *S. himalayana* can retarget enzymes to organelles in *Nicotiana benthamiana*. Transit peptides (TPs) from *S. himalayana* fused to GFP (middle right column) were transiently expressed in *N. benthamiana.* Chlorophyll autofluorescence (far left column) was used for colocalization with plastids. As a positive control for mitochondrial targeting, eqFP611 was tagged with the TP from mitochondrial isovaleryl-CoA dehydrogenase (middle left). The overlay of all three channels is shown on far right. Scale indicated in top right panel is the same for all images. As a means of differentiating true GFP florescence from chlorophyll autofluorescence, yellow ovals are shown in panels to indicate plastids in transformed cells, whereas white ovals indicate plastids in non-transformed cells. Note the presence of GFP-fluorescing stromules in transformed plastids, an indication of true GFP signal. For more information, see methods and supplemental dataset 3.

Generally, organellar tRNAs differ from their cytosolic counterparts in evolutionary history, sequence, and posttranscriptional modification (8). Identity elements that govern the specificity of interaction between a tRNA and its conjugate aaRS may also differ between cytosolic and organellar systems (45). However, in many systems (including plants), cytosolic tRNAs can be imported into the organelles (8, 9, 14). Thus, our observation of retargeting opens some interesting questions – are the retargeted aaRS charging organellar tRNAs? Or is there replacement of the entire organellar system of tRNA aminoacylation, including both tRNAs and aaRS? We investigated the tRNA gene content of the organelles of these species to gain insight into which tRNA substrates may be in use.

### Organelle-encoded tRNA retention often, but not always, mirrors retention of organelle-type aaRS

To investigate the possibility of enzyme-substrate mismatch, we cataloged the presence/absence of organellar tRNA-encoding genes (*trn*) for all sampled heterotrophic species (**Fig. 4**, **Supplemental Dataset 4**). Generally, there was strong correspondence between retention of organellar aaRS and plastome/mitogenome tRNA genes. As the Rafflesiaceae species are thought to lack plastomes entirely, they likely have no plastome-encoded tRNAs (**Fig. 4**). The Balanophoraceae species used in this study also have been reported to exhibit loss of all plastid tRNA genes, except for an extremely divergent *trnE* in *B. fungosa* (46–49). The Rafflesiaceae and Balanophoraceae have additionally lost most of their mitogenome-encoded tRNAs. Correspondingly, these two families also have the most reduced sets of organellar aaRS. At the opposite extreme, *M. hypopitys* retains most of its mitochondrial and plastid tRNA genes and corresponding organellar aaRS.

**Figure 4:**
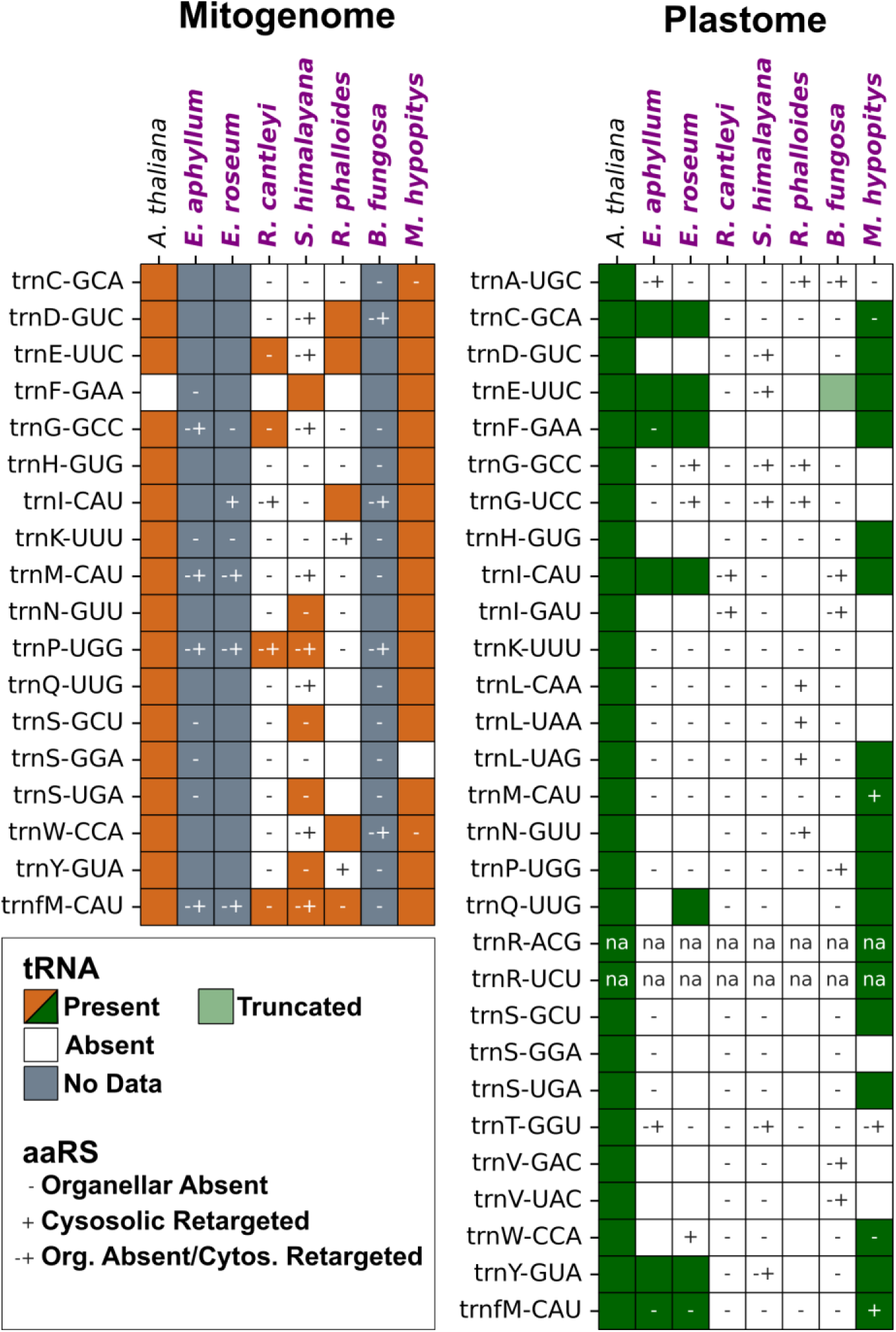
Instances of both concordance and mismatch between organellar tRNA gene loss and aaRS loss/retargeting. Heterotrophic species are written in purple. *Arabidopsis thaliana* is included for reference. Left: mitogenome tRNAs. Right: plastome tRNAs. Color of boxes indicates presence/absence of tRNA. The – and + annotation denotes loss of corresponding organellar aaRS and retargeting of cytosolic tRNA respectively as described for Fig. 2 and Fig. S2. Retargeting predictions are specific for each organelle. Note that loss and retargeting of the corresponding aaRS was not assessed for *trnR* as there are no organelle-specific aaRS orthologs responsible for charging this tRNA. Loss of GatCAB and associated cytosolic GlnRS retargeting are presented for *trnQ*.

However, there are some incidences where a cytosolic enzyme looks to be charging an organellar tRNA due to loss of the organellar aaRS (**Fig. 4**). For example, the few remaining mitogenome-encoded tRNAs in the Rafflesiaceae are missing their organellar aaRS partner (except *S. himalayana* tRNA^Phe^) and are likely charged by a cytosolic enzyme. Alternatively, they may be pseudogenes that have been functionally replaced by an imported tRNA. For both compartments, there are numerous instances where there is no encoded tRNA despite retention of the organellar aaRS (**Fig. 4**). For example, *R. phalloides* retains its organellar SerRS, PheRS, and GlnRS but has lost the corresponding mitochondrial tRNAs. No retargeting of cytosolic enzymes is predicted in these cases, suggesting the organellar enzyme may be acting on an imported cytosolic tRNA.

### Testing for accelerations in organellar aaRS sequence evolution in heterotrophs

Although wholesale loss of many organellar aaRS in heterotrophs precludes analysis of sequence conservation, *M. hypopitys* retained the majority of organellar aaRS despite exhibiting an obligate heterotrophic lifestyle. Thus, we quantified sequence conservation of organellar and cytosolic aaRS in *M. hypopitys* and its autotrophic relative *Rhododendron williamsianum.* We built gene trees and ran relative rate analysis to test for accelerated evolution on the *M. hypopitys* branch (**Supplemental Dataset 5**). We also directly compared branch length ratios of *M. hypopitys* to *R. williamsianum* in cytosolic and organellar aaRS (**Fig. S7, Supplemental Dataset 5).** Neither analysis showed a statistically significant decrease in sequence conservation in organellar enzymes of *M. hypopitys* (**Fig. S7**). However, the deep rooting of our tree on *A. thaliana* may result in poor resolution of the sequence evolution between *M. hypopitys* and *R. williamsianum.* Additionally, the transition to heterotrophy on the *M. hypopitys* branch may have occurred relatively recently compared to the split from *R. williamsianum*, thus diluting any signals of accelerated evolution.

## Discussion

### Loss of photosynthesis relaxes constraints on both plastid and mitochondrial translation machinery

Our primary objective in this study was to test whether the high translational demands of the photosynthetic chloroplast create functional constraint on the evolution of shared translation systems in plastids and mitochondria. To do so, we investigated relative rates of protein evolution in cytosolic and organellar aaRS in heterotrophic plants compared to their photosynthetic relatives. To our surprise, heterotrophic plants underwent widespread loss of organellar aaRS and many bacterial-like enzymes involved in tRNA processing. These losses suggest an extreme relaxation of selection pressures in the absence of a photosynthesizing plastid and support our hypothesis that the translational demands in chloroplasts are the linchpin responsible for the unusually strong functional constraint on the redundant organelle-specific tRNA metabolism in plants. These correlated effects on mitochondrial translation appear to be stronger when the same proteins are shared by mitochondria and plastids (e.g., aaRS) because we found that the most reduced heterotrophs from Rafflesiaceae lost many plastid-specific riboproteins but not their mitochondrial-specific counterparts.

There was variation in the degree of loss across the heterotrophs, often correlated with the number of genes retained in the plastome, particularly tRNAs (see **Supp. Dataset 1** for plastome/mitogenome content). For example, the most aaRS losses occurred in the Rafflesiaceae, all of which are obligate endoparasites and have likely lost their plastomes entirely (33–36, 50). The fewest aaRS losses occurred in *M. hypopitys*, which has the highest number of plastome-encoded genes of all the heterotrophic species we sampled – and notably the greatest number of retained plastid tRNAs (51). Thus, *M. hypopitys* may not be as far along the path of reductive evolution as the other species, which might provide an additional explanation as to why our comparative rate analyses on the remaining organellar aaRS in *M. hypopitys* did not support accelerated evolution compared to cytosolic enzymes (**Fig. S7**). The Balanophoraceae and *Epipogium* species exhibited intermediate degrees of aaRS and plastome gene loss, although the Balanophoraceae species used in this study have lost all plastid tRNAs save for a truncated *trnE* in *B. fungosa* that is likely retained for use in the first step of heme biosynthesis (*46*). An increased plastome mutation rate caused by loss of DNA repair machinery (52, 53) in the Balanophoraceae may explain accelerated plastome (tRNA) gene losses relative to nuclear (aaRS) gene loss. Generally, loss of organellar aaRS correlates well with reduction of the plastome across heterotrophs.

### Loss of organellar aaRS is associated with retargeting of cytosolic aaRS

The wholesale loss of organellar aaRS in heterotrophs was unexpected, as most non-photosynthetic plants still contain a minimal plastome to support heme and fatty acid biosynthesis (32, 54) and fully intact mitogenomes (see **Supp. Dataset 1**). This conundrum prompted us to investigate if cytosolic aaRS are retargeted to organelles in heterotrophs to compensate for gene loss, as has been shown for aaRS/tRNAs in mitochondria of other plant species (55). Retargeting a protein to the organelles generally involves obtaining an N-terminal TP sequence that can be recognized by organellar protein import complexes (56–58). We discovered that many cytosolic aaRS in the heterotrophs had N-terminal sequences predicted to target to one or both organelles (**Fig. 2-3**, **S2-S5**). Therefore, the import of cytosolic aaRS likely compensates for the loss of organellar aaRS and even the loss of the GatCAB complex in Rafflesiaceae via retargeting of cytosolic GlnRS.

There was not perfect correspondence between loss of organellar aaRS and predicted retargeting of cytosolic aaRS. However, the extent of retargeting may have been underestimated for two main reasons. Firstly, TP sequence composition and protein import machinery vary considerably by plant lineage (59), making *in silico* predictions more challenging in uncharacterized species. Thus, some organelle-imported cytosolic aaRS may eschew the use of typical TPs that are key for targeting prediction *in silico* (60). Indeed, some organellar proteins are known to not rely at all on N-terminal TPs for import (61, 62). TP sequence divergence could also be accelerated by alteration of the sequence or subunit composition of protein import machinery. Notably, our presence/absence analysis of plastid and mitochondrial protein import machinery revealed some critical subunits for plastid protein import are lost in the heterotrophs, especially the Rafflesiaceae, which could influence which sequences are recognized as transit peptides (**Fig. S6**). For example, Toc159, one of the key outer envelope import receptors involved in TP binding and recognition, is missing from the Rafflesiaceae (63). This subunit specializes in binding and import of proteins involved in photosynthesis, whereas Toc132, which is retained in these species, specializes in “non-green” protein import (64, 65). The Rafflesiaceae are also missing Tic236, the protein that spans the space between the envelope membranes to physically connect the inner and outer envelope (66). This raises interesting questions about the nature of plastid protein import and the ultrastructure of envelope membranes in Rafflesiaceae. Other heterotrophs in addition to the Rafflesiaceae are also missing some subunits present in *A. thaliana*; *Epipogium* species, *M. hypopitys* and the Rafflesiaceae lack one or more Tic20 and Tic22 paralogs. The Tic22 protein family is highly conserved and expressed, with a hypothesized role in outer membrane maturation in addition to its role in protein import (67, 68). Study of Tic-22 loss-of-function mutants revealed that these proteins are not essential, at least in *A. thaliana* (67, 69). Tic20 forms the structural basis of the inner pore of the translocon, with Tic20-I/Tic20-IV being the essential isoforms for plant survival (63, 70–72). Similar trends have also recently been found by Guo et al. 2024 (73). These alterations to the composition of the TIC-TOC complex may reduce the specificity of plastid protein targeting predictions. In contrast, the heterotrophic lines retained almost all TIM-TOM subunits forming the mitochondrial translocon. The divergence of the TIC-TOC complex in some heterotrophs may result in accelerated evolution of TPs, thus, targeting predictions made based on characteristics of photosynthetic plants may not be fully accurate.

Secondly, uncertain protein models for some of our species may have led to true N-terminal sequences being absent or incomplete. Retargeting of cytosolic aaRS may occur through two different mechanisms; 1) alternative splicing or alternative transcription/translation start sites within one gene resulting in cytosolic and organellar isoforms and 2) duplicated genes within the genome encoding cytosolic and organellar paralogs. Our ability to investigate these two mechanisms was limited as we generally only had transcriptome or genome information but not both for each species. However, observations of transcriptomes assembled with consideration for alternative splice isoforms indicate that both mechanisms are potentially at play (**Supplemental Dataset 6**). For example, *Rhopalocnemis phalloides* has two separate paralogs for cytosolic ThrRS, one of which is predicted to be mitochondrial-targeted (Rphallo_DN1669_c0_g1) while the other is not (Rphallo_DN1669_c0_g2). Conversely, *R. phalloides* TyrRS is encoded by a single locus (Rphallo_DN204_c0_g1) with a multitude of splice isoforms, some of which are putatively mitochondrial-targeted. Genome sequencing and long-read mRNA sequencing would allow for more thorough analysis of which mechanism is most common in each species.

While almost all the TPs that we investigated with *in vivo* microscopy assays exhibited targeting to mitochondria and/or plastids, the specific organelle sometimes differed from the *in silico* prediction. The ProRS and TrpRS results in particular were unexpected as the *in silico* predictions were strongly mitochondrial (**Supp. Dataset 3**), but we observed clear plastid localization *in vivo* in two separate rounds of imaging with different vectors (**Fig 3**, **Fig. S4, S5**). There were other TPs where targeting predictions were only partly supported by GFP localization experiments, including GluRS and GlyRS which exhibited targeting to only one of the two predicted organelles, and AsnRS which was only predicted to have plastid localization but additionally exhibited mitochondrial localization *in vivo*.

These results may reflect the fact that target peptide prediction models like those used in this study do not have complete precision or sensitivity for a given subcellular compartment even when applied to benchmark datasets. This is not surprising, as changing just a few amino acids in a transit peptide can alter protein targeting from singular to dual targeting *in vivo* (74). Furthermore, predictions made by a classifier trained on a broad sampling of TPs from diverse plant species cannot accurately predict every TP for an individual species, since species-specific features may not be represented enough to be detected by the model. Even if we assume our microscopy data are a perfect reflection of “true” targeting in the native system, we might expect some inaccuracies in our targeting predictions just based on the sensitivity and precision of the prediction algorithms (TargetP-2.0 and LOCALIZER). Of the two models, TargetP-2.0 specifically identifies mitochondrial and plastid transit peptides with 85-86% sensitivity and 87-90% precision and differentiates proteins lacking TPs with over 98% sensitivity and precision (75, 76). The publication describing LOCALIZER does not directly quantify the tool’s ability to distinguish proteins with no transit peptide but has 60-73% sensitivity and 60-80% precision for mitochondrial and plastid predictions respectively, counting proteins with predicted dual-targeting as positive hits for both plastid and mitochondria.

The artificial nature of heterologous expression in *N. benthamian* can also create artefacts. We can see that in some cases predictions for a given TP can change when it is fused to GFP as opposed to the native aaRS sequence (Supp. Dataset 3). This implies that more than just the N-terminal extension of a protein determines targeting and may create some artefacts in assessing targeting based on TP-GFP fusions. Furthermore, alternative splicing or translation initiation sites can be a mechanism of alternative targeting (77–79). Alternative translation initiation may require RNA cis regulatory factors i.e., 5¢ UTR sequences not captured in our heterologous expression system (80). Additionally, the import of a heterologously expressed TP-tagged GFP can change based on the age of leaves during infiltration, the efficiency of protein folding, protein expression levels, and truncation of an internal targeting sequence (81–83). As comparison of the *in silico* and heterologous expression assays is effectively combining the errors created by both methods, we would not expect perfect concordance between the methods regarding the specific identity of the retargeted organelle. Rather, these methods are complementary tools to assess general retargeting to the endosymbiotic organelles rather than differentiating between TPs specifically targeting mitochondria or plastids from those that are dual targeted. An additional method for future investigation of targeting to plastids or mitochondria would be to subfractionate tissue from one or more of these heterotrophic plants species and conduct comparative high throughput proteomics on each fraction.

### Potential rewiring of aaRS-tRNA interactions

Retargeting of cytosolic aaRS led us to investigate organelle genome tRNA content and whether there was substrate-enzyme mismatch in the organelles, i.e. instances where the cytosolic enzyme was interacting with an organellar tRNA or vice-versa. Our interpretations presume that loss of organellar tRNA genes is compensated by import of the corresponding cytosolic tRNAs (9, 14, 84). We found that the retention of organellar aaRS in heterotrophs generally matched the retention of plastid tRNAs but not necessarily mitochondrial tRNAs (**Fig. 4**). This is further supported by observations in hemiparasite *Viscum album*, which has lost most of its mitochondrial tRNAs, yet retains most organellar aaRS in conjunction with a nearly full set of plastid tRNAs (32, 85, 86). Substrate-enzyme mismatch was rare within the plastid and usually involved loss of a plastid tRNA yet continued presence of the corresponding organellar aaRS (e.g., see *R. phalloides* where all plastid tRNAs are lost but a handful of organellar aaRS remain). There were also instances of organellar aaRS apparently charging imported cytosolic tRNAs in the mitochondria (e.g., *trnF* in *R. cantleyi and R. phalloides*). More surprisingly, however, there were cases where a mitochondrial tRNA was retained despite loss of the corresponding organellar aaRS and apparent retargeting of the cytosolic enzyme, particularly in the Rafflesiaceae. This type of mismatch is unexpected because cytosolic aaRS are thought to generally be more discriminating enzymes than their mitochondrial counterparts (8, 55, 87–89), making it more difficult to transition to a new tRNA substrate. However, our results suggest some cytosolic enzymes may be interacting with bacterial-type tRNAs in these heterotrophic species, although we cannot rule out the possibility that cytosolic tRNAs are being redundantly imported. Indeed, our findings raise fascinating questions about the process of functionally replacing tRNAs and interacting enzymes in these dynamic systems (55, 90), which could be explored with comparative and biochemical approaches in closely related species still undergoing these transitions. Given that parasitic plant lineages tend to be fast evolving across all three genomes (91), the cytosolic aaRS in these species may have evolved lower specificity and the capability to charge organellar tRNAs. Overall, our findings suggest that the plastid tRNA content, and not the mitochondrial tRNA content, drives retention of organellar aaRS, again placing the photosynthetic plastid as a central factor in the retention of functionally redundant translation machinery.

The identity of retained organellar aaRS versus those substituted by cytosolic counterparts was seemingly non-random. GluRS and PheRS were the two mostly highly retained organellar aaRS across these diverse lineages of plants (**Fig. 1**). Organellar PheRS is likely retained because the cytosolic PheRS is unusual in that it is a heteromeric complex with independently encoded α and β subunits (92). Therefore, functional replacement of the organellar PheRS would require two independent acquisitions of TPs, as well as stoichiometric import and assembly within the organelles. Such a coordinated set of changes may be unlikely to evolve (24, 55). The one exception was *E. aphyllum,* for which we could not find an organellar PheRS. However, this may be an artefact of low expression, as protein models for this species were curated from RNA-seq data. As tRNA^Glu^ is the most frequently retained tRNA in heterotrophic plant plastomes due to its additional role in the first step of heme biosynthesis (54, 93, 94), it is not surprising that GluRS was among the most retained organellar aaRS in this study.

Intriguingly, the *Rafflesiaceae* are the only species that lost the organellar GluRS, as well as plastid *trnE*. Yet, genes encoding heme synthesis enzymes are retained in these species (92), opening interesting questions into how they carry out the first step in the pathway. In *S. himalayana*, we found retargeting of the cytosolic GluRS to the plastid, which along with presumed import of the cytosolic tRNA^Glu^, could functionally replace the first step in the heme biosynthesis pathway. This has the interesting implication that the glutamyl-tRNA reductase catalyzing the next step in heme biosynthesis is non-discriminating towards cytosolic tRNAs. The organellar GluRS is also necessary for mis-aminoacylation of tRNA^Gln^ prior to transamidation of Glu to Gln by the GatCAB complex(96). Cytosolic GlnRS is retargeted to the mitochondria in *S. himalayana*, thus allowing for direct aminoacylation of tRNA^Gln^ assuming there is also import of cytosolic tRNA^Gln^. This likely precipitated the loss of the GatCAB complex and negated the need for a non-discriminating organellar GluRS.

At the other extreme, there was repeated loss of organellar AlaRS, which was absent from all heterotrophs and the hemiparasitic *Cuscuta* species, possibly because plastomes of heterotrophs are often AT-rich and thus have underrepresentation of amino acids with GC-rich codons like Ala, Gly and Pro (47, 48). Thus, the corresponding organellar aaRS and tRNAs may be subject to decreased functional constraint and be less likely to be retained. This hypothesis is supported by the fact that we also see frequent loss of organellar GlyRS and ProRS in heterotrophs (**Fig. 1**).

Our results indicate that imported cytosolic aaRS and tRNAs are sufficient to sustain mitochondrial translation in plants, supporting the hypothesis that the selective pressure to keep organellar aaRS/tRNAs is maintained by translational demands associated with photosynthesis. Yet other eukaryote lineages that never bore plastids often still retain organelle-specific tRNAs and aaRS for their mitochondria (8, 9, 22, 24). This is even more puzzling when considering that various eukaryotic lineages have jettisoned nearly all their mitochondrial tRNAs/aaRS, including cnidarians, ctenophores, some fungi, and trypanosomes (12, 24, 97). One possible explanation is the logical parallel to our finding that photosynthesis governs plastid translation rates and maintains gene redundancy; eukaryotic lineages with high respiratory rates may require high mitochondrial gene expression and, therefore, retain mitochondria-specific translation machinery. Indeed, mitochondrial gene expression tracks well with respiratory rate and cellular activity across different tissues and metabolic states (98–100). This connection to cell energetics would also explain why mitochondrial tRNAs/aaRS are dispensable in parasitic eukaryotes like trypanosomes and heterotrophic plants, which presumably have lower energy demands. By studying non-model plant species with unusual physiology and life histories, we were able to link plastid bioenergetic processes to the evolution of both organellar and nuclear genomes. The sequencing of mitogenomes and characterization of metabolism in non-model eukaryotes will provide exciting opportunities to conduct similar studies to better understand how mitochondrial metabolism effects gene redundancy and elucidate the principles governing endosymbiotic integration.

## Methods

### Data acquisition and ortholog search

Gene models for species used in the study were collected from publicly available data sources (**Supplemental Dataset 1**). When protein sequences were not available, we generated sequences from transcriptome data using Transdecoder v5.5.0 (101), applying a minimum protein length of 50 amino acids and retaining only protein sequences with blast hits to any sequence in the Uniprot Taxonomic Divisions Plant database (https://ftp.uniprot.org/pub/databases/uniprot/knowledgebase/taxonomic_divisions/). Then, the Transdecoder primary_transcript.fa script was used to extract only the longest translated protein sequence from the primary transcript for each gene to reduce the search space. SeqKit 2.3.0 (102) was used to create uniform gene handles for all files and remove stop codons from sequences. Orthofinder v2.5.5 (37) was run on all sets of protein models with a user-defined species tree (see **Fig. 1** for tree topology) to find groups of likely orthologous proteins, i.e. “orthogroups” in two steps as follows:

orthofinder.py -f <folder_with_protein_models> -t 24 -a 6

orthofinder.py -ft <folder_with_results_> -s userdefined_speciestree.txt -t 24 -a 6

Orthogroups containing proteins of interest including cytosolic/organellar aaRS and tRNA-processing enzymes characterized in *A. thaliana* (**Supplemental Dataset 2**) were extracted and checked for completeness via RBH against all protein models for a given species using BLASTP 2.15.0+ (103) and custom Python/Bash scripts. After ortholog discovery via Orthofinder, full sets of protein models (not just the longest) were searched for *A. thaliana* protein sets using BLASTP to ensure that narrowing of the search space for Orthofinder had not resulted in loss of matching gene models. In cases where available curated protein models excluded alternative transcripts, transcriptome data were obtained and Transdecoder was run to predict protein-coding sequences (see **Supplemental Dataset 1** for details). Note that ArgRS was not included in the ortholog search as there are no distinct organellar versions of this enzyme in *A. thaliana.* One ArgRS gene product is cytosol-targeted, and the other is triple-targeted to cytosol, mitochondria, and plastids (9, 16). In the case of GlnRS, there is no organellar ortholog as plants typically use the bacterial-type GatCAB amidotransferase system to charge Gln-tRNA indirectly in mitochondria and plastids (42). For CysRS, tRNase Z and PRORP, sequences for both cytosolic and organellar proteins were clustered together in one orthogroup likely due to recent shared ancestry. For these genes, cytosolic vs. organellar versions were differentiated by aligning sequences in each orthogroup using MAFFT v7.525 (104) with the --auto setting, followed by trimming of the alignment using trimAl v1.5.rev0 on the -automated1 setting (105). Alignments were then viewed using MEGA 11 (Tamura et al., 2021). Maximum-likelihood tree building was conducted using RAxML v 8.2.12 (107) as follows:

raxmlHPC-PTHREADS-SSE3 -f a -# 100 -p 12345 -x 12345 -s <fasta file with orthogroup sequences> -n

<orthogroup> -m PROTGAMMALG -T 12

Inclusion in cytosolic versus organellar groups was then determined by relationship to known *A. thaliana* cytosolic/nuclear/organellar proteins in maximum-likelihood trees (**Fig. S8-10**).

Once we had a curated set of protein sequences for all species, they were checked for completeness of length by alignment with *A. thaliana* orthologs by MAFFT and subsequent visualization with a custom Python script. A “full-length” ortholog was defined as having at least 50% alignment with any *A. thaliana* protein in the orthogroup, excluding pseudogenes. Custom Python and Bash scripts are available via Dryad (https://doi.org/10.5061/dryad.0cfxpnw7p).

### In silico targeting predictions

For TP analysis, all protein models available for each gene with similarity to the original Orthofinder hit (BLASTP e-value < 1e-6) were used, not just the longest. All transcripts for cytosolic aaRS, a “positive control” set of genes involved in mitochondrial oxidative phosphorylation/ plastid housekeeping and a “negative control” set of genes involved in cell-wall synthesis were run through targeting prediction programs LOCALIZER-1.0.5 (76) and TargetP-2.0 (75), with commands as follows:

LOCALIZER.py -p -i <input file>

targetp -fasta <input file> -mature -org pl

Protein models for some species were often incomplete due to the fragmented nature of some of the transcriptome assemblies. To ensure models missing the N-terminus were not used in retargeting analysis, predictions for models that did not align without gaps for 15 residues within 100 amino acids of the start codon for at least one *A. thaliana* ortholog were discarded. For transcript-by-transcript heatmaps (**Supplemental Dataset 6**), a custom R script was used to plot all transcripts for all species. For Fig. S3 (which shows predicted targeting for a multitude of different proteins) only the most retargeted ortholog for each species was used. Orthologs were then binned based on probability of targeting to the mitochondria or plastids. We chose a probability of 0.5 as a threshold for LOCALIZER to match the original publication describing the tool(76). TargetP-2.0 uses a “winner take all” strategy to declare one subcellular localization based on the location with the highest likelihood score, which is not compatible with assessing dual targeting (75, 108). Thus, we used the lowest likelihood score for known organelle-targeted *A. thaliana* proteins from our control dataset identified by TargetP-2.0 to be plastid or mitochondria targeted (∼0.5 for AT1G02410 cytochrome c oxidase 11). High confidence retargeting was defined as a > 0.5 probability/likelihood of targeting for both LOCALIZER and TargetP-2.0. Moderate confidence was defined as a > 0.5 probability/likelihood for one of the algorithms. Low confidence retargeting was defined as neither algorithm providing a probability/likelihood > 0.5. For subsequent analysis of ratios (**Fig. 2**, **Fig. S1-S2**), any ortholog with a “moderate” or better confidence was counted as retargeted. Note that *R. cantleyi* and *S. himalayana* proteins were excluded from predictions of targeting to plastids as these species likely lack plastomes. All analyses including plotting and statistics were carried out using custom Bash, Python, and R scripts available via Dryad (https://doi.org/10.5061/dryad.6hdr7sr5x).

### In vivo localization of S. himalayana aaRS transit peptides via tobacco infiltration and confocal microscopy

TPs were chosen based on having an at least moderate probability (see above) of targeting a protein to either the mitochondria or plastid and/or having an N-terminal extension preceding conserved sequence shared with *A. thaliana.* We selected either the sequence predicted to be a TP by TargetP-2.0 plus 10 residues or the entire N-terminal overhang plus the first 10 residues aligning with the functional portion of the *A. thaliana* protein. These amino acid sequences were then converted into DNA sequences using EMBOSS Backtranseq (109) with the *Nicotiana tabacum* codon usage table (see **Supplemental Dataset 3**). These sequences were then used to generate constructs for *Agrobacterium* transformation using a newly designed dual-expression vector containing both the mitochondrial control IVD-FP611 and GFP fused to the C-terminus of our TP of interest. Some TPs were also originally cloned into the single-gene expression vector described in (55). Transformation into *N. benthamiana* and confocal microscopy was constructed as described in (55). With our dual expression vectors, there was occasionally low GFP and/or FP611 signal relative to chlorophyll autofluorescence (see Fig. 3 GlyRS and MetRS). Thus, we scaled pixel intensity using look-up tables (LUTs) for all images to better see localization for low expression constructs. It is important to note that in cases where GFP fluorescence is low and LUTs are scaled, overlap of the chlorophyll florescence and GFP spectra can create the false impression of plastid GFP targeting (see negative control in Fig. S4 for an example). Thus, we only determined a TP to be plastid targeted if we either observed 1) GFP-fluorescing stromules in transformed plastids, an indication of true GFP signal as these structures occlude thylakoids or 2) Increased GFP florescence in plastids of transformed cells compared to untransformed cells in the same frame. Often guard cells would not be transformed, so we also used these as an in-frame negative control. Note that *in silico* predictions shown are for the TP attached to GFP, not the original protein model. Relative to the whole protein models, the GFP predictions yielded higher targeting probabilities and affected classification for MetRS (LOCALIZER likelihood score > 0.5 for mitochondria) and AsnRS (TargetP-2.0 likelihood score > 0.5 for plastid), but there were no substantive changes for any of the other transit peptides. For more information see supplemental dataset 3. Constructs are available from Addgene; see supplemental dataset 3 for ID numbers.

### Mitogenome assembly

Prior to tRNA analysis, the mitogenomes of *S. himalayana* (SRR629601) and *R. cantleyi* (SRR629613) were assembled *de novo* from raw reads available from the NCBI SRA database. To maximize the likelihood of finding all mitogenome tRNAs, genomes were assembled in several ways. Reads were first trimmed using FASTP (110), with either default parameters or “harsh” trimming where the --qualified_quality_phred and --unqualified_percent_limit parameters were set to 20 and 10, respectively. SPAdes v.3.15.4 (111) was then used to assemble trimmed reads *de novo,* with kmer sizes of 21, 33, 55, and 77. To avoid potential assembly fragmentation due to excessive levels of sequence coverage, a subset of reads (1,000,000 from S. *himalayana* and 500,000 from *R. cantleyi*) were sampled and assembled from each species as well.

### tRNA analysis

Excepting the Rafflesiaceae, for which we assembled mitogenomes *de novo,* the GenBank (.gbk) files for mitogenomes and plastomes from species of interest were downloaded from NCBI (**Supplemental Dataset 1**). A custom Perl script was used to extract annotated tRNAs from the .gbk files. tRNAscan-SE-2.0.9 (112, 113) was run on downloaded genomes (parameters -BQH), and results were cross-checked with preexisting GenBank annotations (**Supplemental Dataset 4**). For *S. himalayana* and *R. cantleyi*, the same tRNAscan-SE-2.0.9 pipeline was used on all generated assemblies, and the consensus set of tRNAs was checked for plant origin using BLASTN against all NCBI reference sequences and the *A. thaliana* mitogenome (NC_037304.1) and plastome (NC_000932.1). tRNA presence/absence patterns were plotted along with the corresponding aaRS presence/absence/retargeting data using a custom Python script. Scripts are available via Dryad (https://doi.org/10.5061/dryad.np5hqc009). The highly divergent and truncated trnE for *B. fungosa* was added based on prior publication(46).

### Rate Analysis

Primary protein sequences for cytosolic and organellar aaRS from heterotroph *M. hypopitys* and related autotroph *R. williamsianum* from the original Orthofinder search were used for comparison of rates, with *A. thaliana* orthologs forming the outgroup. CysRS, LeuRS, ThrRS, and TrpRS were excluded from the analysis because the organellar ortholog was missing or too short to create informative gene trees in one or both test species. Predicted TPs were trimmed from sequences using output of TargetP-2.0.

Sequences were then aligned using MAFFT v7.490 (104) and trimmed manually in MEGA 11 (106). Maximum likelihood trees were built using RAxML v 8.2.12 (107) as follows:

raxmlHPC-PTHREADS-SSE3 -f a -# 100 -p 12345 -x 12345 -s <orthogroup_aligned.fa> -n

<orthogroup> -m PROTGAMMALG -T 12

For orthogroups containing only three ortholog sequences, a “dummy” outgroup sequence represented by the *A. thaliana* ortholog with one branch-specific substitution was created to overcome RAxML’s input requirement of four or more taxa. The resulting trees were then evaluated for topology. In instances where species-specific paralogs were not sister to one another, separate multiple sequence alignments and trees were re-made for each paralog as described above. In instances where species-specific paralogs were sister, both sequences were retained in the same tree. Trees were rooted to the *A. thaliana* ortholog(s), then branch lengths to the last shared node from each *H. monotropa* and *R. williamsianum* ortholog were extracted from the trees. For instances where two sister paralogs from the same species were retained, the branch lengths were averaged. Then, the log10-transformed ratio of the *H. monotropa/R williamsianum* branches was plotted, and mean branch lengths for cytosolic and organellar enzymes were calculated. Statistical difference in mean branch ratio in cytosolic vs. organellar enzymes was tested using a *t*-test. Tajima’s relative rate tests were run in MEGA 11 using *A. thaliana* as the outgroup. Data is available in **Supplemental Dataset 5.** Scripts, alignments, and maximum likelihood trees are available via Dryad (https://doi.org/10.5061/dryad.tb2rbp05r).

## Acknowledgements

We thank Jessica Warren for helpful discussion and feedback on this project and Anna Pratt for helpful advice on the cloning approach. We thank undergraduate researchers Xiaorui Lou and Janna Novak for their assistance. This work was supported by grants MCB-2322154, MCB-2048407, and MCB-1933590 from the National Science Foundation (NSF) and utilized the RMACC Summit supercomputer (NSF ACI-1532235 and ACI-1532236). RAD was supported by an NSF Postdoctoral Research Fellowship in Biology (IOS-2208908). JC was supported by the University of Nebraska Foundation.

---

**Supplemental Dataset 1:** Species and datasets used for gene presence/absence, retargeting analysis, and tRNA content analysis.

**Supplemental Dataset 2.** Genes of interest and counts of gene copies for each species.

**Supplemental Dataset 3:** *S. himalayana* transit peptide sequences for transient expression and colocalization in *N. benthamiana*

**Supplemental Dataset 4:** tRNA content of organelle genomes.

**Supplemental Dataset 5:** Relative branch lengths and Tajima’s relative rate test for aaRS in Ericales spp.

**Supplemental Dataset 6:** Heatmaps for probability of subcellular targeting to plastids or mitochondria on a protein-by-protein basis.

**Supplemental Table 1:**
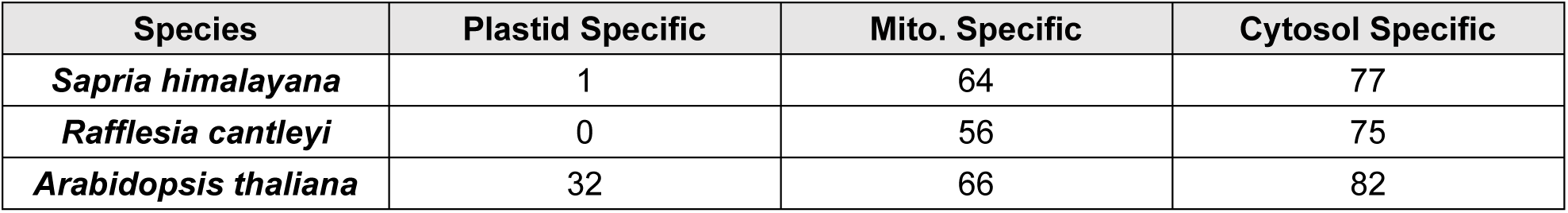
Count of Orthogroups Containing Riboprotein Ortholog(s)

| Species | Plastid Specific | Mito. Specific | Cytosol Specific |
| --- | --- | --- | --- |
| *Sapria himalayana* | 1 | 64 | 77 |
| *Rafflesia cantleyi* | 0 | 56 | 75 |
| *Arabidopsis thaliana* | 32 | 66 | 82 |

**Supplemental Table 2:** aaRS targeting and recovery of control groups.

| Protein Class | Mean % Mito-targeted |  | Mean % Plastid-targeted |  |
| --- | --- | --- | --- | --- |
|  | Autotrophs | Heterotrophs | Autotrophs | Heterotrophs |
| **Cytosolic aaRS** | 8.9% | 20.3% | 9%* | 15.2%* |
| **Mitochondrial OXPHOS** | 59.9% | 58.5% | 8.1% | 8.3% |
| **Plastid Housekeeping** | 20.5% | 19.9% | 92.6%* | 86.6%* |
| **Cell Wall Synth.** | 6.4% | 11.2% | 15.5%* | 9%* |

\*Rafflesiaceae not included in calculation

**Supplemental Figure 1:**
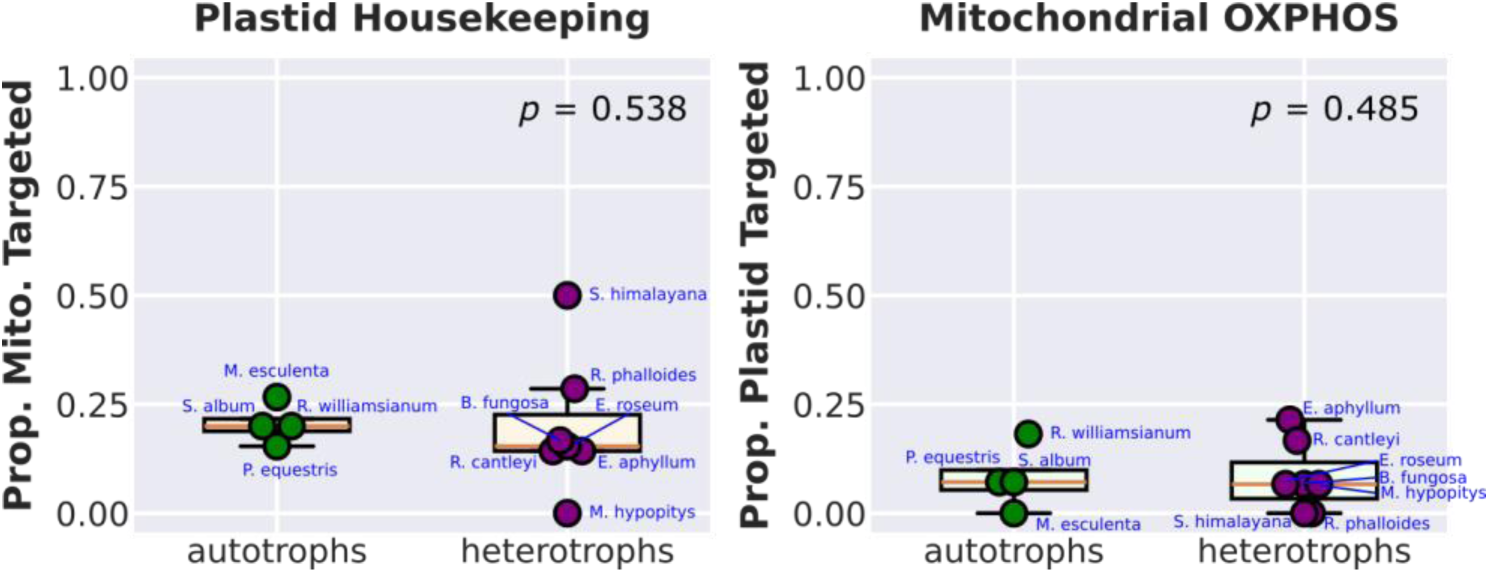
Proportion of Plastid Housekeeping Genes with Predicted Mitochondrial Retargeting (Left), and Proportion of Mitochondrial OXPHOS proteins with Predicted Plastid Retargeting (Right). Proportion of enzymes with predicted retargeting (probability ≥ 0.5) based on the outputs of LOCALIZER and/or TargetP. Autotrophs and heterotrophs are plotted in green and purple, respectively. Reported *p*-values from *t*-tests or Mann-Whitney U tests depending on distribution of the data. Note that *R. cantleyi* and *S. himalayana* are not included in plastid retargeting analysis due to suspected loss of the plastome. It should be noted that proportions of plastid housekeeping genes in the Rafflesiaceae represent predictions for only two proteins; all other proteins are missing from these lineages, see Fig. S3.

**Supplemental Figure 2:**
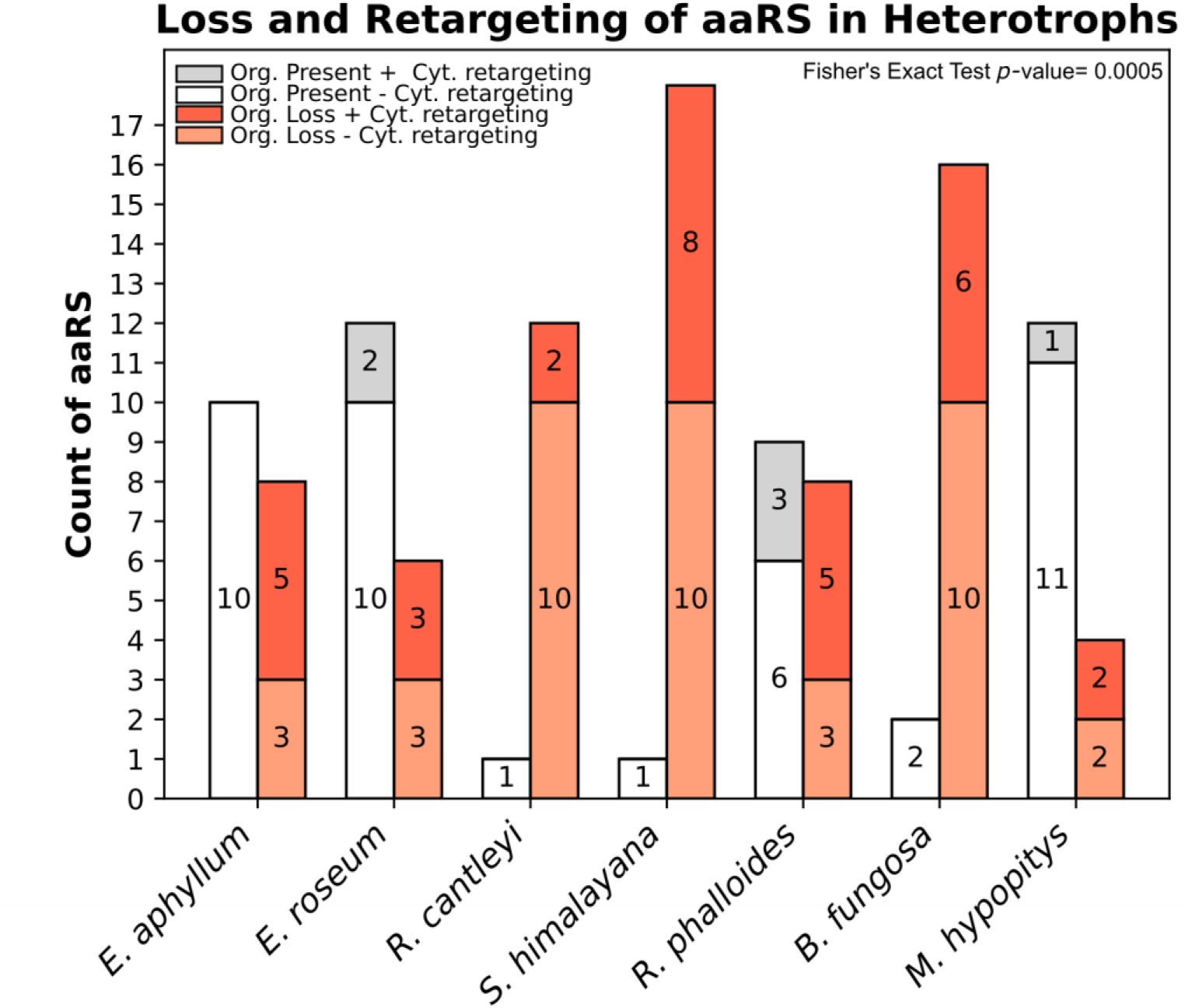
Significant overlap between loss of organellar aaRS and retargeting of cytosolic counterpart in heterotrophs. Reported counts refer to number of aaRS “families” defined by amino acids (e.g., GluRS, PheRS, TrpRS, etc.). Organellar gene presence/absence determined as for Fig. 1. Partial sequences for organellar enzymes are counted as present. Predicted retargeting (probability ≥ 0.5) to either mitochondria or plastids was based on the outputs of LOCALIZER and/or TargetP. Note that aaRS where cytosolic targeting could not be tested (i.e., enzyme was not present, or N-terminus was incomplete) were excluded. Organellar enzyme present = shades of gray. Organellar enzyme absent = shades of orange. Cytosolic enzyme retargeting = darker shading. No retargeting = lighter shading. Reported *p*-value is from a Fisher’s exact test for correlation between loss of organellar aaRS and retargeting of the corresponding cytosolic aaRS. Note that *R. cantleyi* and *S. himalayana* proteins are excluded from predictions of targeting to plastids as these species lack plastid genomes. Loss of GatCAB and associated cytosolic GlnRS retargeting are included in the counts.

**Supplemental Figure 3:**
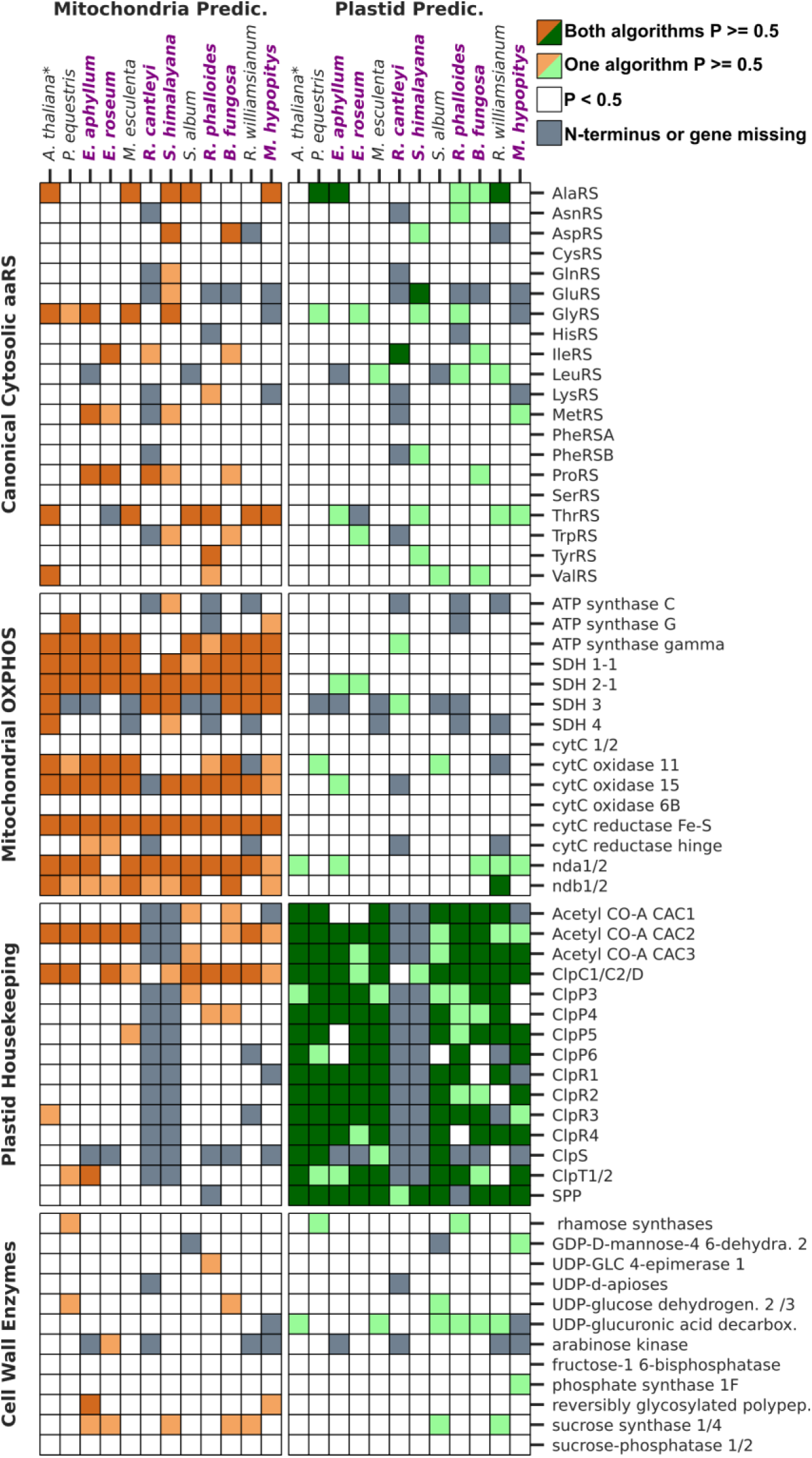
Targeting predictions for cytosolic-type aaRS. Deeper shades of green (plastid, right) and orange (mitochondrion, left) indicate probability of retargeting to that organelle is ≥ 0.5 based on both LOCALIZER and TargetP. Lighter shades of orange/green indicate one of the two algorithms yielded a probability ≥ 0.5. White indicates probability for both algorithms was < 0.5. Gray indicates that the gene/transcript was not present or that the N-terminus was incomplete, so no targeting prediction could be made. Heterotrophs are written in purple, closely related autotrophs in black. \**Arabidopsis thaliana* is included as a reference.

**Supplemental Figure 4:**
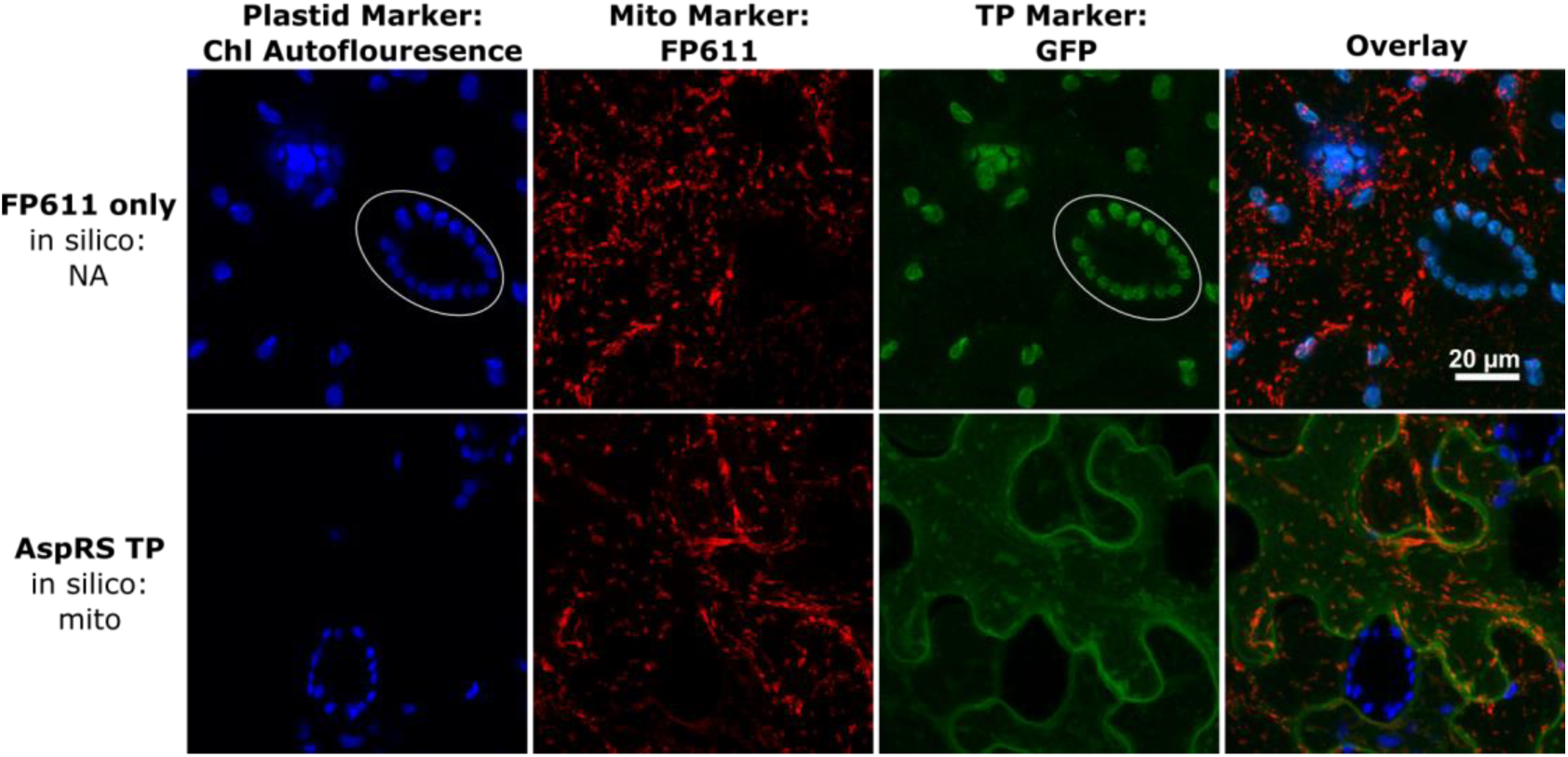
Cytosolic-type aaRS transit peptides from heterotroph *S. himalayana* transiently expressed in *Nicotiana benthamiana*. Transit peptides (TPs) from *S. himalayana* fused to GFP (middle right column) were transiently expressed in *N. benthamiana.* Chlorophyll autofluorescence (far left column) was used for colocalization with plastids. As a positive control for mitochondrial targeting, eqFP611 was tagged with the TP from mitochondrial isovaleryl-CoA dehydrogenase (IVD) (middle left). Overlay of all three channels on far right. Top row: IVD_FP611 infiltration only-negative control. White ovals indicate plastids in untransformed cells, allowing assessment of bleed-over chlorophyll autofluorescence in the GFP channel. See main text for how true plastid localization was distinguished from bleed-over signal. Bottom row: Expression of AspRS TP led to apparent aggregates of GFP in the cytosol, potentially resulting from an artefact due to overexpression. Scale indicated in top right panel is the same for all images.

**Supplemental Figure 5:**
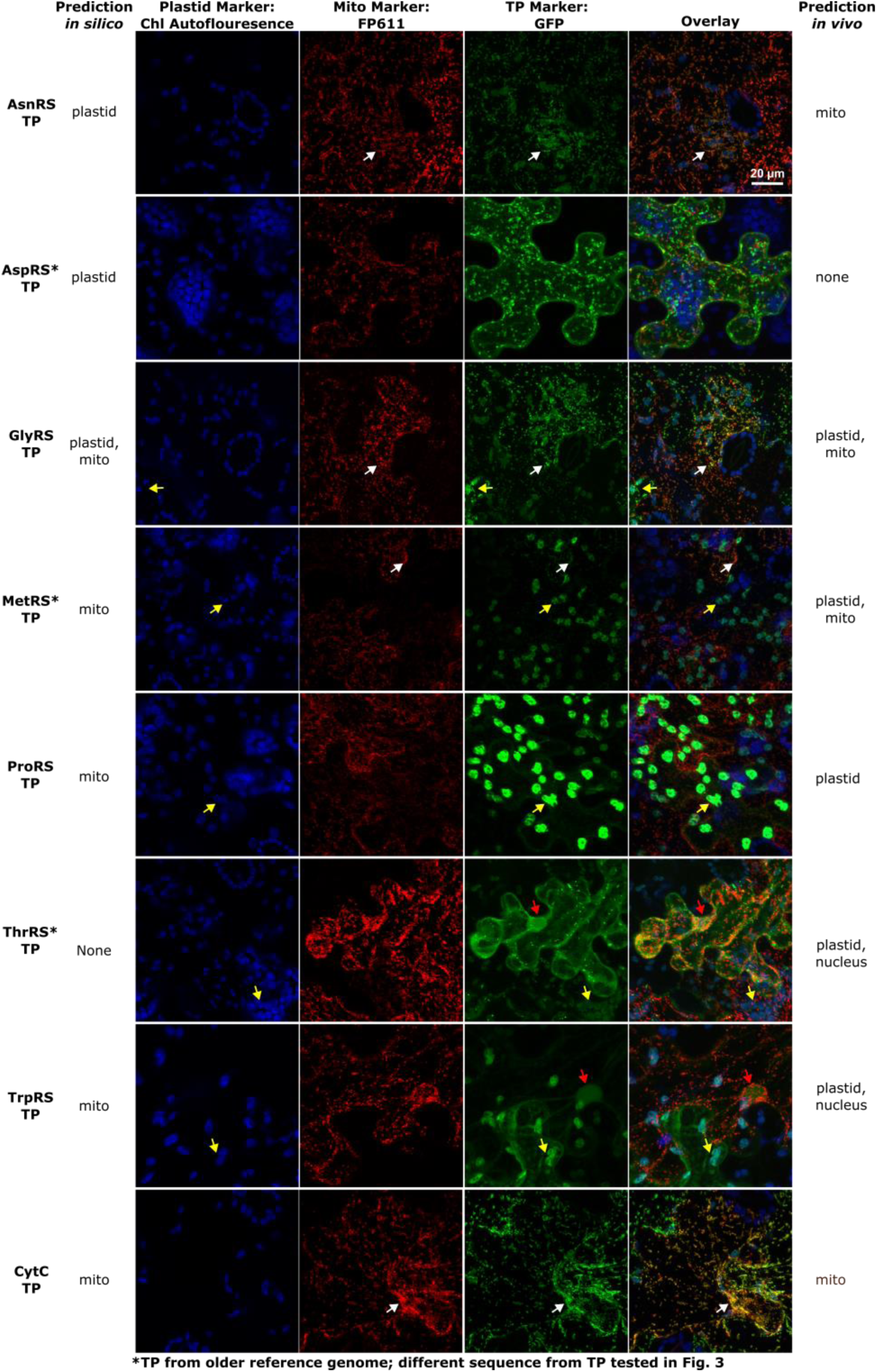
Initial round of microscopy data using single expression vector PK7FWG2. Transit peptides (TPs) from *S. himalayana* fused to GFP (middle right column) were transiently expressed in *N. benthamiana.* Chlorophyll autofluorescence (far left column) was used for colocalization with plastids. As a positive control for mitochondrial targeting, eqFP611 was tagged with the TP from mitochondrial isovaleryl-CoA dehydrogenase (middle left). The overlay of all three channels on far right. Arrows indicate examples of plastid (yellow), mitochondrial (white) and potential nuclear (red) localization. Scale indicated in top right panel is the same for all images.

**Supplemental Figure 6:**
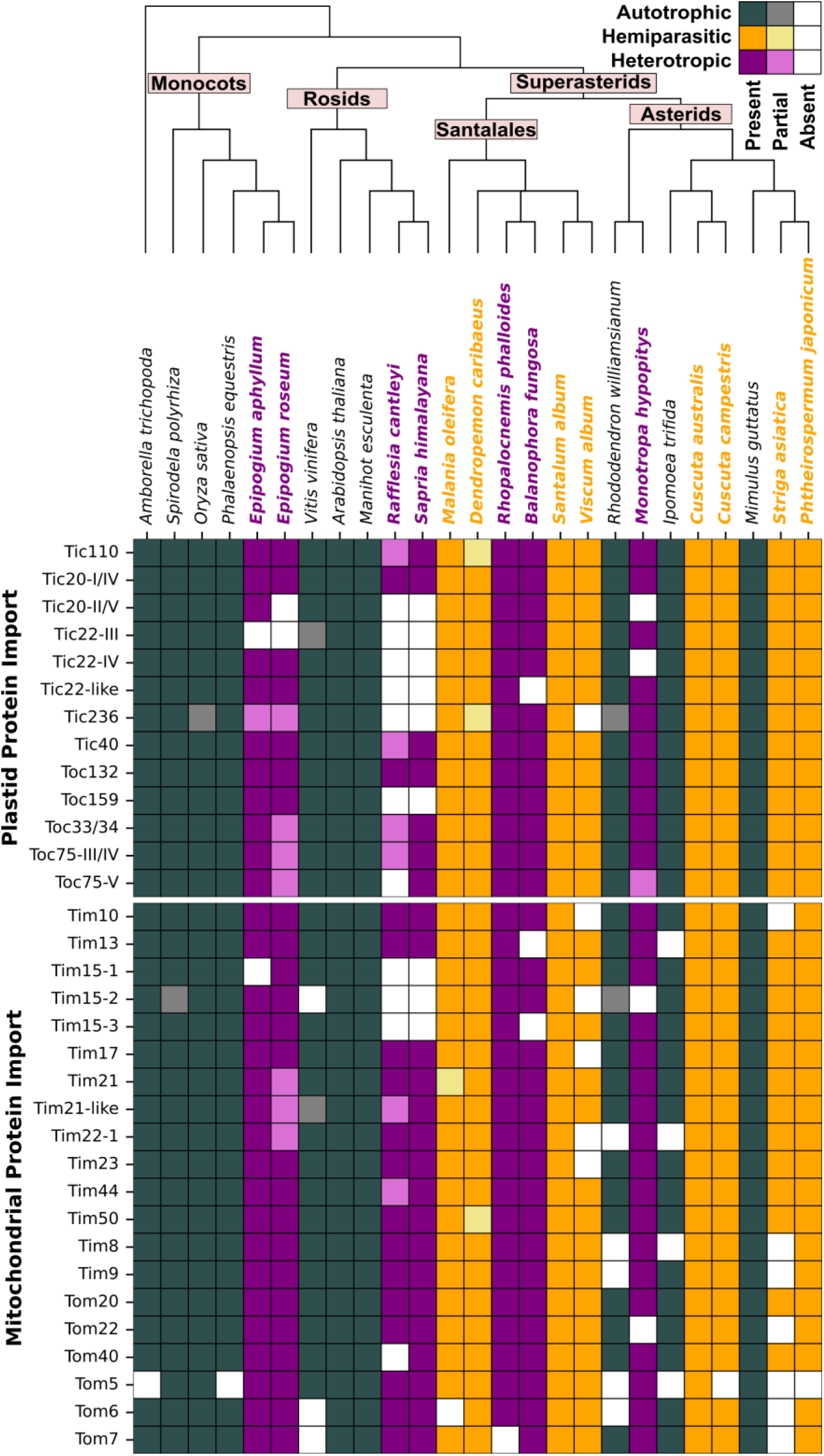
Loss of key subunits for plastid import apparatus in *Rafflesiaceae*. For complete names of proteins and corresponding *A. thaliana* gene ID, see supplemental dataset 2. Hue of boxes indicates metabolism (gray = autotroph, orange = hemiparasite, purple = heterotroph), shade indicates degree of presence/absence (white = no sequence found, light colors = partial sequence found, dark colors = complete or near complete sequence found).

**Supplemental Figure 7:**
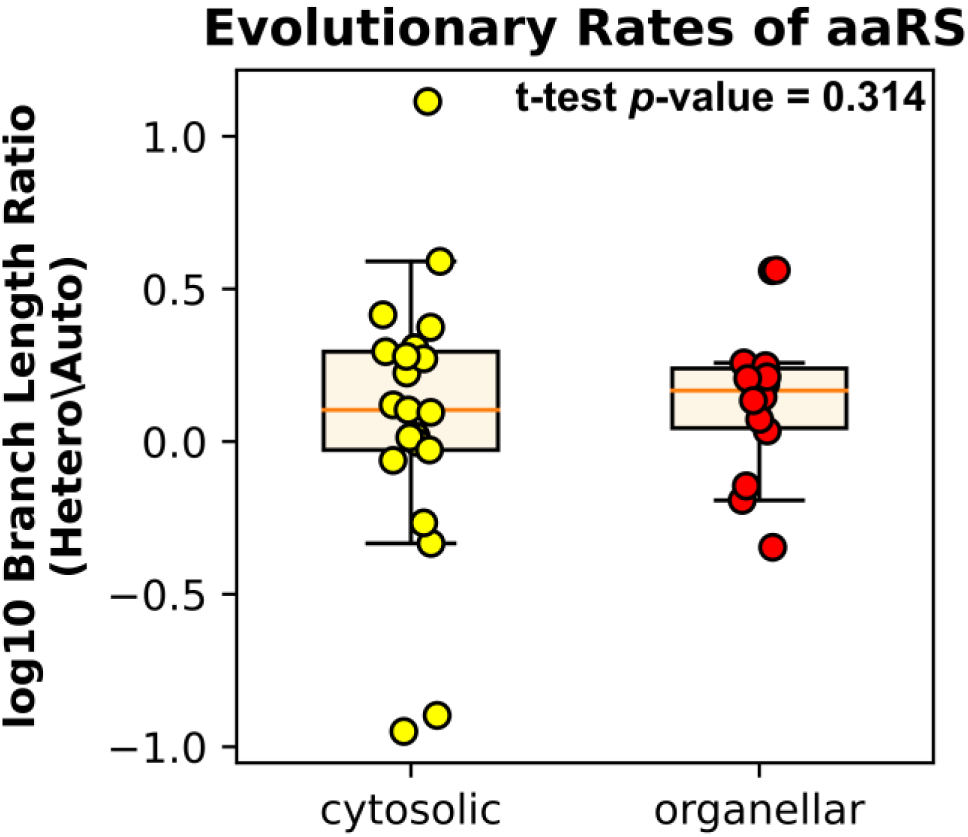
Relative rates of sequence evolution of cytosolic and organellar aaRS are similar in heterotroph compared to autotrophic relative. Each point represents an aaRS branch length of *M. hypopitys* (heterotrophic) normalized to branch length of *R. williamsianum* (autotrophic) for canonically cytosolic and organellar aaRSs (log10 scale).

**Supplemental Figure 8:**
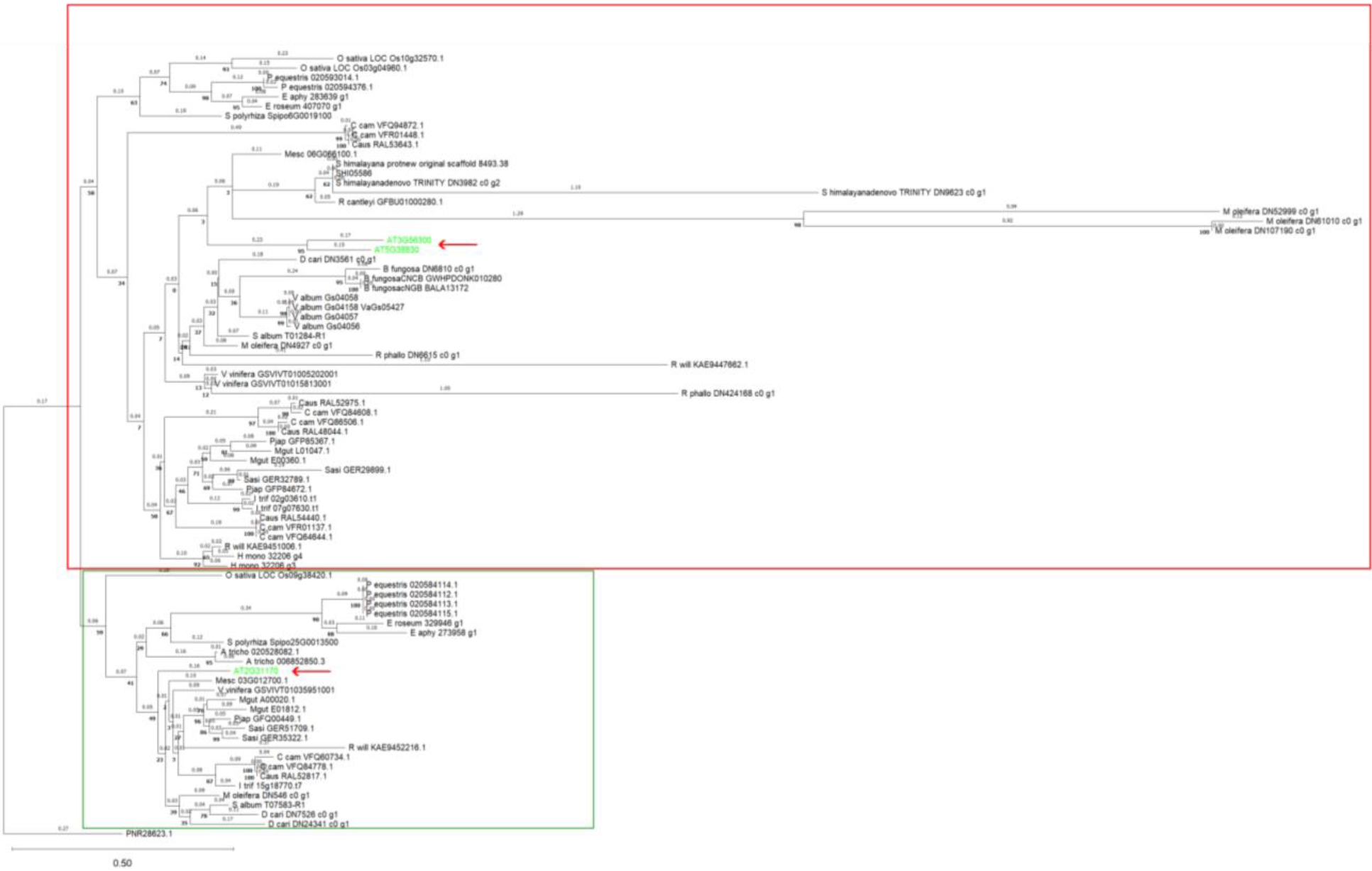
Maximum likelihood tree for CysRS. Bootstrap values are indicated at nodes. Branch length labels represent average substitutions per site. Red arrow denotes *A. thaliana* sequences for organellar and cytosolic CysRS. Orthologs for *A. thaliana* organellar CysRS (AT2G31170) outlined in green, orthologs for *A. thaliana* cytosolic CysRS (AT3G56300, AT5G38830) outlined in red.

**Supplemental Figure 9:**
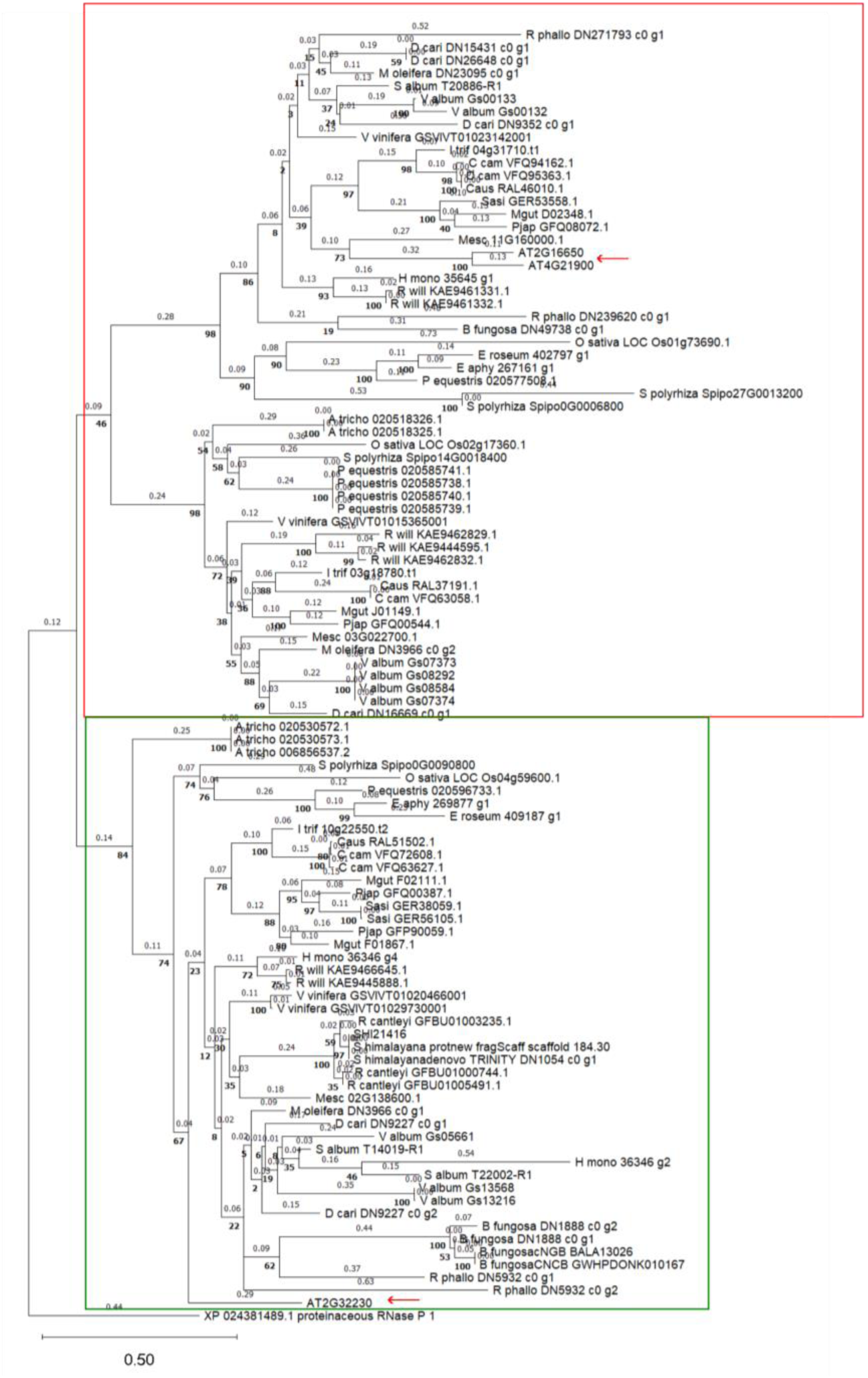
Maximum likelihood tree for Protein Only RNase Ps (PRORPs). Bootstrap values are indicated at nodes. Branch length labels represent average substitutions per site. Red arrow denotes *A. thaliana* sequences for nuclear and organellar PRORPs. Orthologs for *A. thaliana* organellar PRORP1 (AT2G32230) outlined in green, Orthologs for *A. thaliana* cytosolic PRORP2/3 (AT2G16650, AT4G21900) outlined in red.

**Supplemental Figure 10:**
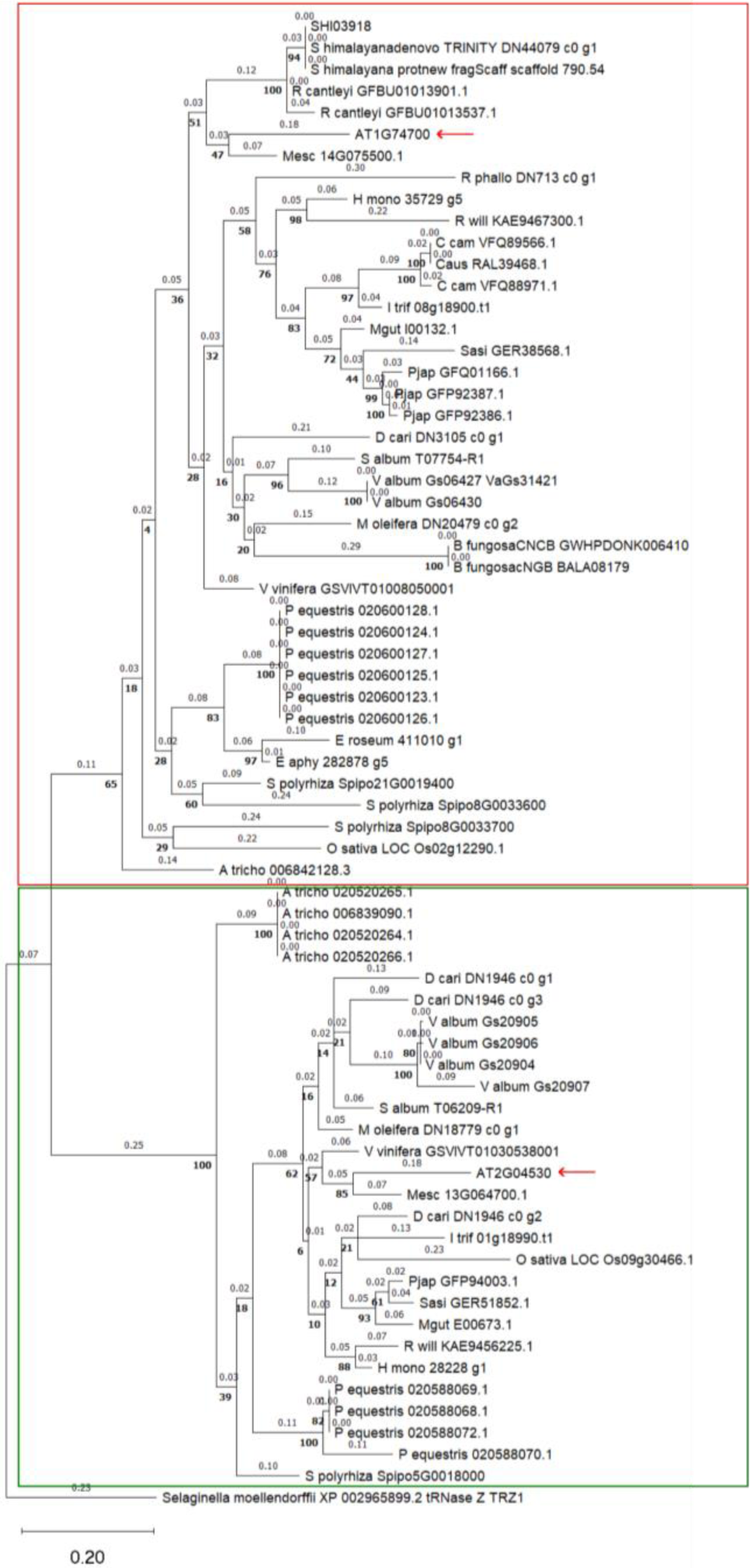
Maximum likelihood tree for tRNase Z 1/2. Bootstrap values are indicated at nodes. Branch length labels represent average substitutions per site. Red arrow denotes *A. thaliana* sequences for tRNAse Z type proteins. Orthologs for *A. thaliana* cytosolic tRNase Z1 (AT1G74700) are outlined in red, and orthologs for *A. thaliana* plastid tRNase Z2 (AT2G04530) are outlined in green.

