## Supplemental Dataset 6 for "Photosynthetic demands on translational machinery drive retention of redundant tRNA metabolism in plant organelles"

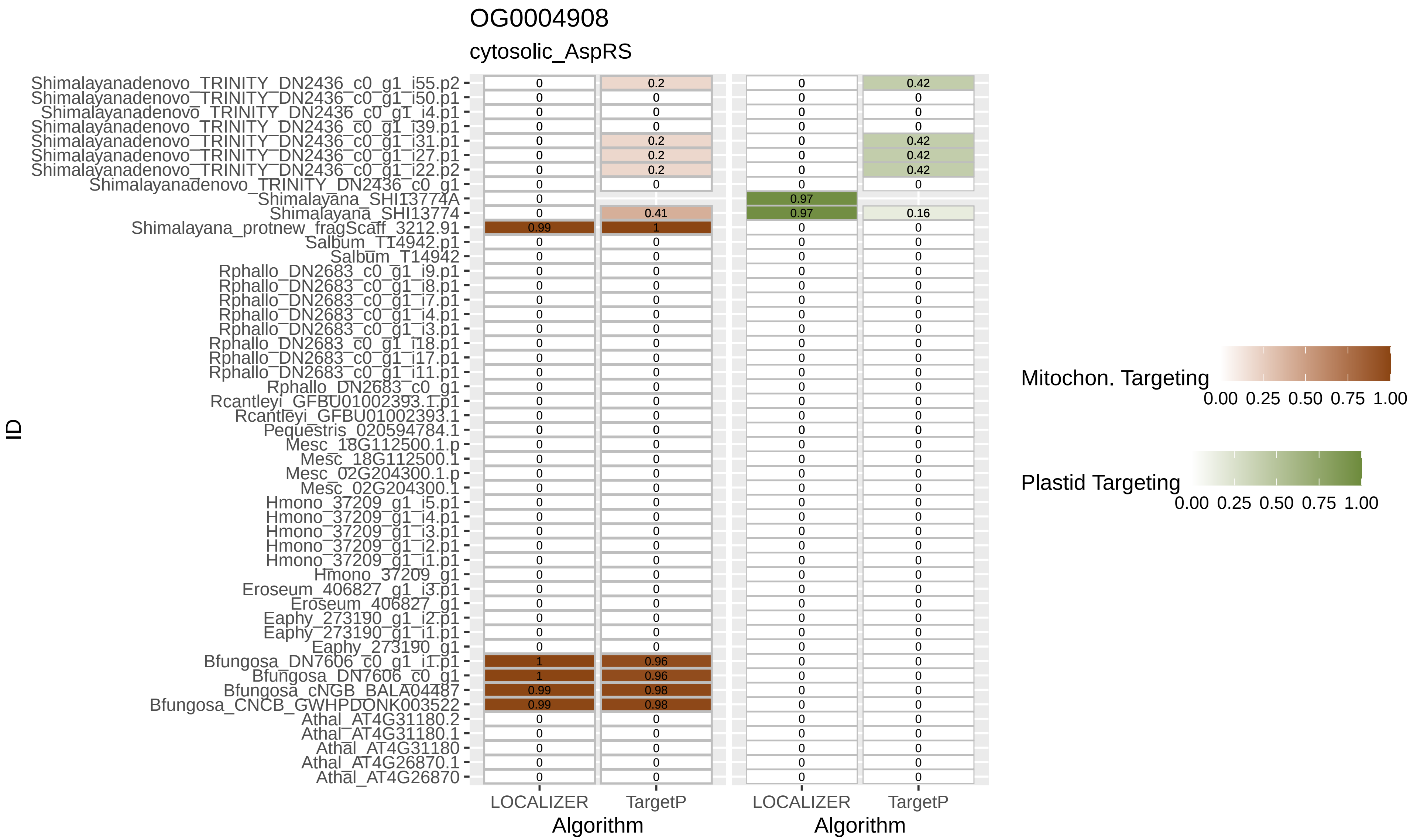
OG0004908
cytosolic_AspRS
Shimalayanadenovo_TRINITY_DN2436_c0_g1_i55.p2
Shimalayanadenovo_TRINITY_DN2436_c0_g1_i50.p1
Shimalayanadenovo_TRINITY_DN2436_c0_g1_i4.p1
Shimalayanadenovo_TRINITY_DN2436_c0_g1_i39.p1
Shimalayanadenovo_TRINITY_DN2436_c0_g1_i31.p1
Shimalayanadenovo_TRINITY_DN2436_c0_g1_i27.p1
Shimalayanadenovo_TRINITY_DN2436_c0_g1_i22.p2
Shimalayanadenovo_TRINITY_DN2436_c0_g1
Shimalayana_SHI13774A
Shimalayana_SHI13774
Shimalayana_protnew_fragScaff_3212.91
Salbum_T14942.p1
Salbum_T14942
Rphallo_DN2683_c0_g1_i9.p1
Rphallo_DN2683_c0_g1_i8.p1
Rphallo_DN2683_c0_g1_i7.p1
Rphallo_DN2683_c0_g1_i4.p1
Rphallo_DN2683_c0_g1_i3.p1
Rphallo_DN2683_c0_g1_i18.p1
Rphallo_DN2683_c0_g1_i17.p1
Rphallo_DN2683_c0_g1_i11.p1
Rphallo_DN2683_c0_g1
Rcantleyi_GFBU01002393.1.p1
Rcantleyi_GFBU01002393.1
Pequestris_020594784.1
Mesc_18G112500.1.p
Mesc_18G112500.1
Mesc_02G204300.1.p
Mesc_02G204300.1
Hmono_37209_g1_i5.p1
Hmono_37209_g1_i4.p1
Hmono_37209_g1_i3.p1
Hmono_37209_g1_i2.p1
Hmono_37209_g1_i1.p1
Hmono_37209_g1
Eroseum_406827_g1_i3.p1
Eroseum_406827_g1
Eaphy_273190_g1_i2.p1
Eaphy_273190_g1_i1.p1
Eaphy_273190_g1
Bfungosa_DN7606_c0_g1_i1.p1
Bfungosa_DN7606_c0_g1
Bfungosa_cNGB_BALA04487
Bfungosa_CNCB_GWHPDONK003522
Athal_AT4G31180.2
Athal_AT4G31180.1
Athal_AT4G31180
Athal_AT4G26870.1
Athal_AT4G26870
ID
LOCALIZER
TargetP
Algorithm
LOCALIZER
TargetP
Algorithm
Mitochon. Targeting
0.00 0.25 0.50 0.75 1.00
Plastid Targeting
0.00 0.25 0.50 0.75 1.00

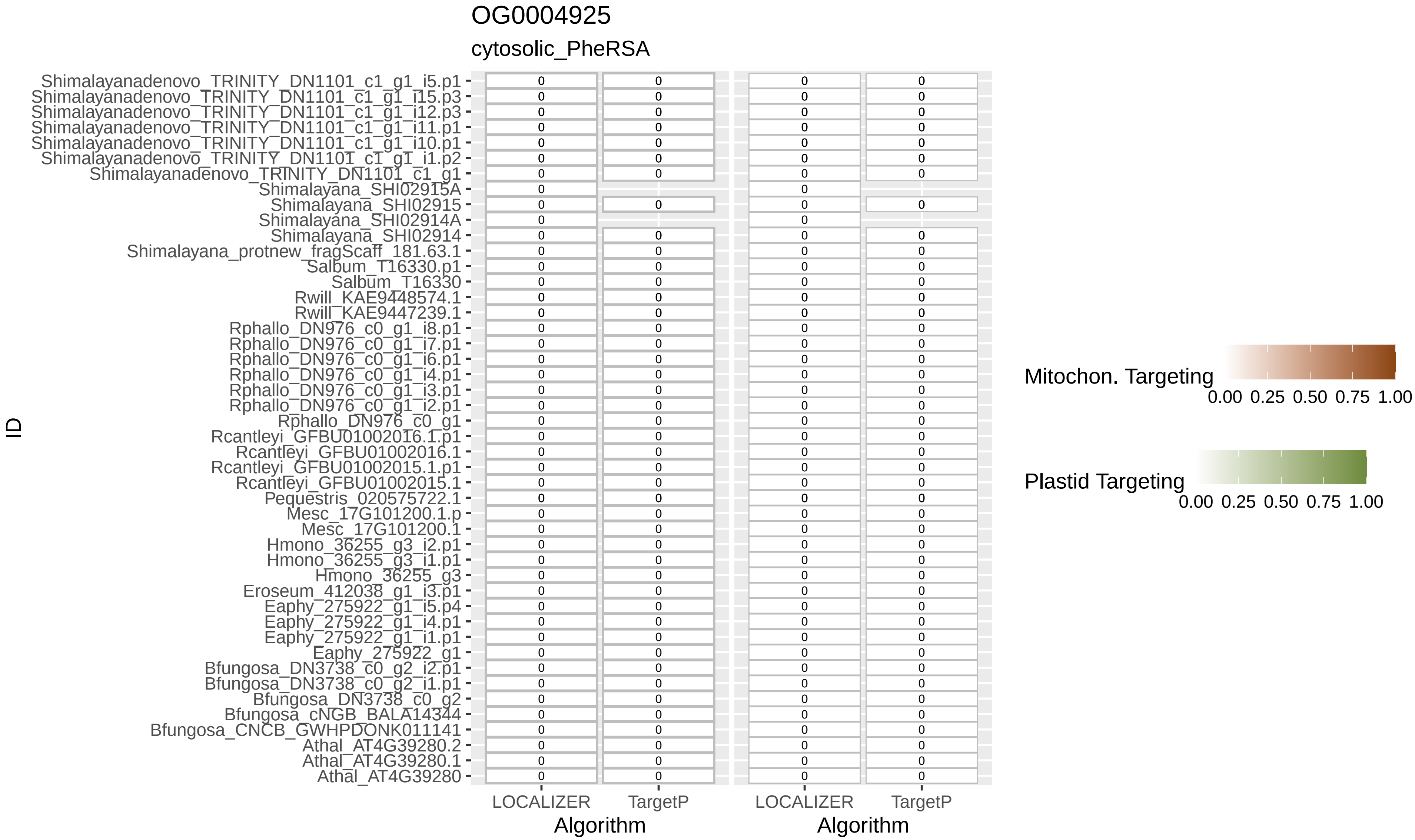
OG0004925
cytosolic_PheRSA
Shimalayanadenovo_TRINITY_DN1101_c1_g1_i5.p1
Shimalayanadenovo_TRINITY_DN1101_c1_g1_i15.p3
Shimalayanadenovo_TRINITY_DN1101_c1_g1_i12.p3
Shimalayanadenovo_TRINITY_DN1101_c1_g1_i11.p1
Shimalayanadenovo_TRINITY_DN1101_c1_g1_i10.p1
Shimalayanadenovo_TRINITY_DN1101_c1_g1_i1.p2
Shimalayanadenovo_TRINITY_DN1101_c1_g1
Shimalayana_SHI02915A
Shimalayana_SHI02915
Shimalayana_SHI02914A
Shimalayana_SHI02914
Shimalayana_protnew_fragScaff_181.63.1
Salbum_T16330.p1
Salbum_T16330
Rwill_KAE9448574.1
Rwill_KAE9447239.1
Rphallo_DN976_c0_g1_i8.p1
Rphallo_DN976_c0_g1_i7.p1
Rphallo_DN976_c0_g1_i6.p1
Rphallo_DN976_c0_g1_i4.p1
Rphallo_DN976_c0_g1_i3.p1
Rphallo_DN976_c0_g1_i2.p1
Rphallo_DN976_c0_g1
Rcantleyi_GFBU01002016.1.p1
Rcantleyi_GFBU01002016.1
Rcantleyi_GFBU01002015.1.p1
Rcantleyi_GFBU01002015.1
Pequestris_020575722.1
Mesc_17G101200.1.p
Mesc_17G101200.1
Hmono_36255_g3_i2.p1
Hmono_36255_g3_i1.p1
Hmono_36255_g3
Eroseum_412038_g1_i3.p1
Eaphy_275922_g1_i5.p4
Eaphy_275922_g1_i4.p1
Eaphy_275922_g1_i1.p1
Eaphy_275922_g1
Bfungosa_DN3738_c0_g2_i2.p1
Bfungosa_DN3738_c0_g2_i1.p1
Bfungosa_DN3738_c0_g2
Bfungosa_cNGB_BALA14344
Bfungosa_CNCB_GWHPDONK011141
Athal_AT4G39280.2
Athal_AT4G39280.1
Athal_AT4G39280
ID
LOCALIZER
TargetP
Algorithm
LOCALIZER
TargetP
Algorithm
Mitochon. Targeting
0.00 0.25 0.50 0.75 1.00
Plastid Targeting
0.00 0.25 0.50 0.75 1.00

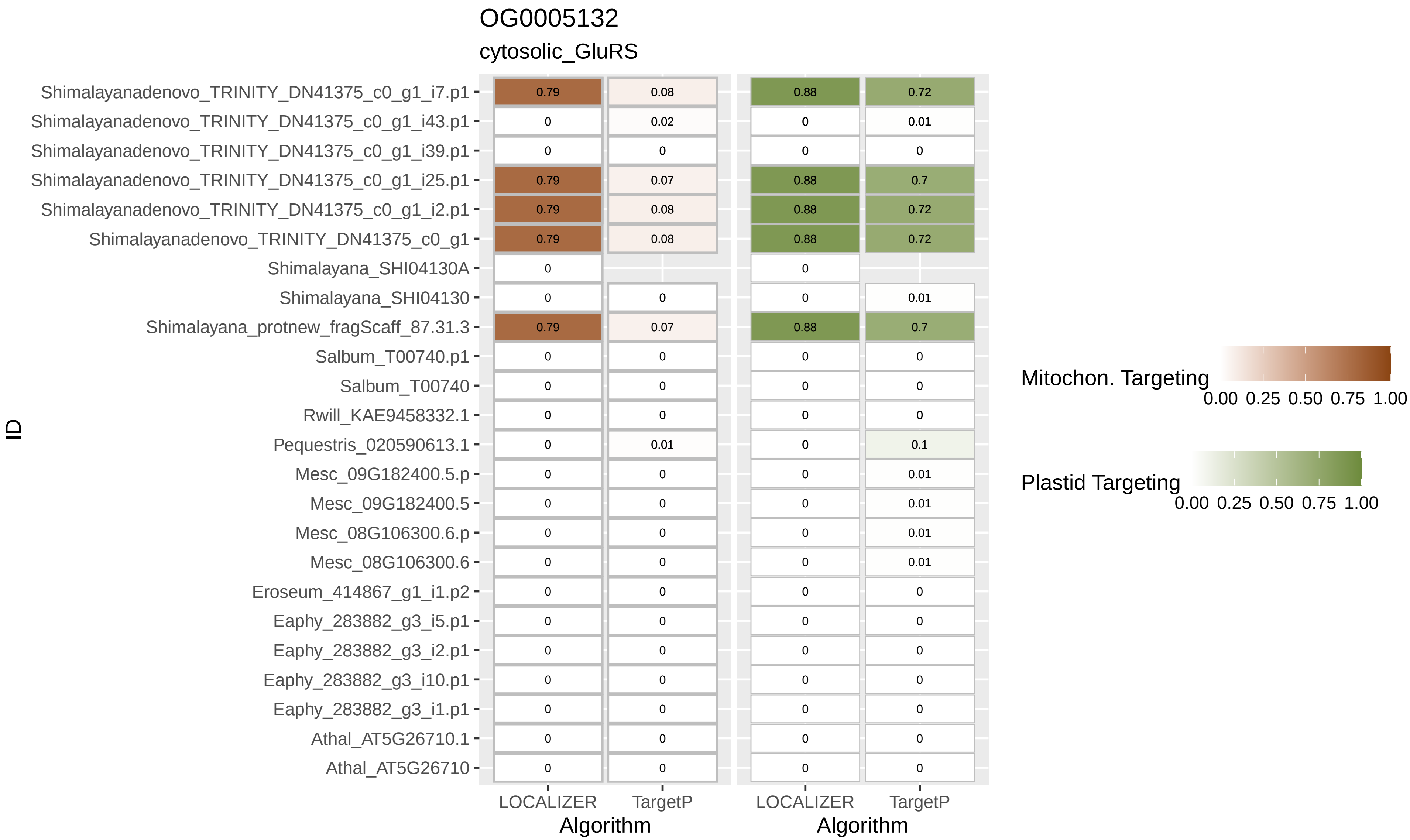
OG0005132
cytosolic_GluRS
Shimalayanadenovo_TRINITY_DN41375_c0_g1_i7.p1
Shimalayanadenovo_TRINITY_DN41375_c0_g1_i43.p1
Shimalayanadenovo_TRINITY_DN41375_c0_g1_i39.p1
Shimalayanadenovo_TRINITY_DN41375_c0_g1_i25.p1
Shimalayanadenovo_TRINITY_DN41375_c0_g1_i2.p1
Shimalayanadenovo_TRINITY_DN41375_c0_g1
Shimalayana_SHI04130A
Shimalayana_SHI04130
Shimalayana_protnew_fragScaff_87.31.3
Salbum_T00740.p1
Salbum_T00740
Rwill_KAE9458332.1
Pequestris_020590613.1
Mesc_09G182400.5.p
Mesc_09G182400.5
Mesc_08G106300.6.p
Mesc_08G106300.6
Eroseum_414867_g1_i1.p2
Eaphy_283882_g3_i5.p1
Eaphy_283882_g3_i2.p1
Eaphy_283882_g3_i10.p1
Eaphy_283882_g3_i1.p1
Athal_AT5G26710.1
Athal_AT5G26710
ID
LOCALIZER
TargetP
Algorithm
LOCALIZER
TargetP
Algorithm
Mitochon. Targeting
0.00 0.25 0.50 0.75 1.00
Plastid Targeting
0.00 0.25 0.50 0.75 1.00

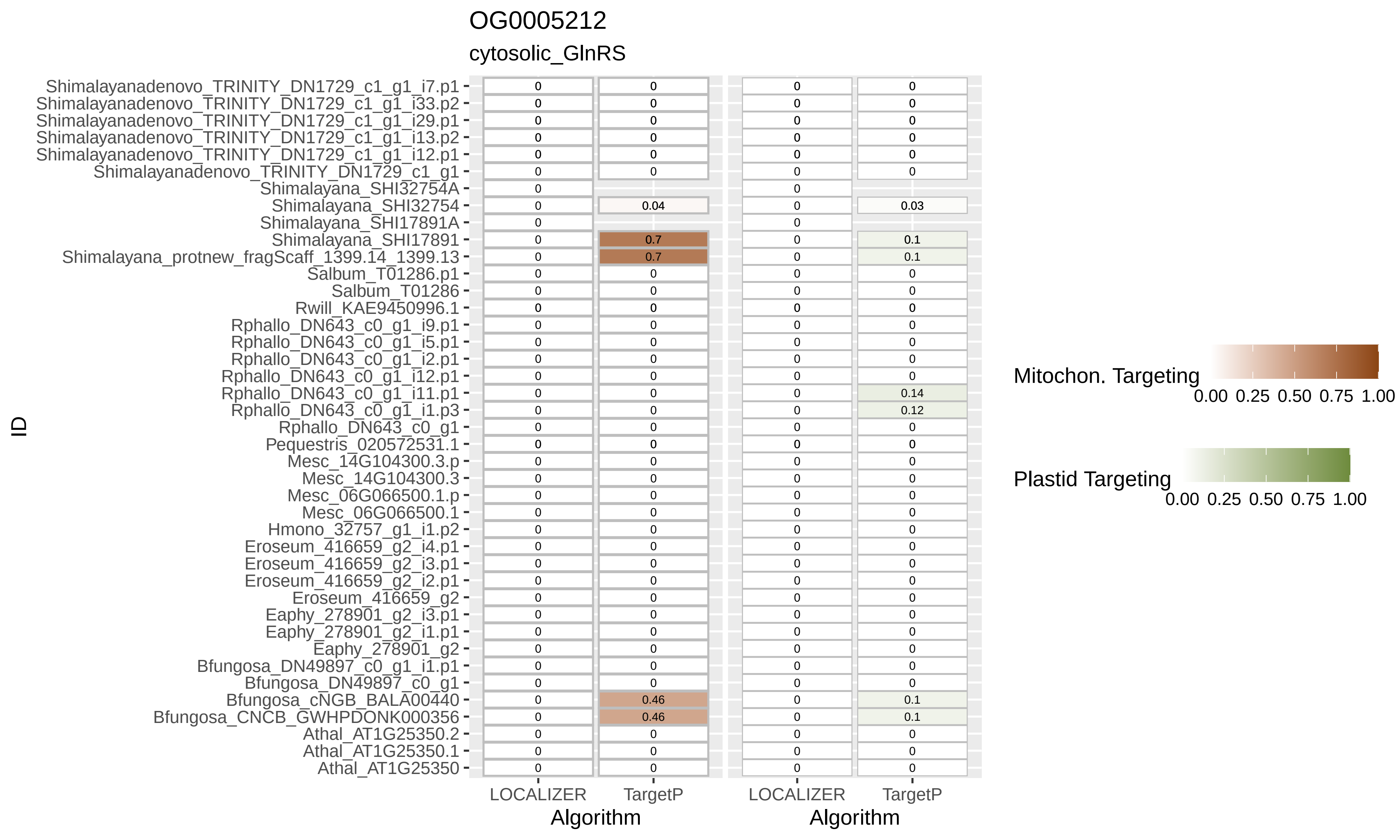

OG0005212
cytosolic_GlnRS
ID
Shimalayanadenovo_TRINITY_DN1729_c1_g1_i7.p1
Shimalayanadenovo_TRINITY_DN1729_c1_g1_i33.p2
Shimalayanadenovo_TRINITY_DN1729_c1_g1_i29.p1
Shimalayanadenovo_TRINITY_DN1729_c1_g1_i13.p2
Shimalayanadenovo_TRINITY_DN1729_c1_g1_i12.p1
Shimalayanadenovo_TRINITY_DN1729_c1_g1
Shimalayana_SHI32754A
Shimalayana_SHI32754
Shimalayana_SHI17891A
Shimalayana_SHI17891
Shimalayana_protnew_fragScaff_1399.14_1399.13
Salbum_T01286.p1
Salbum_T01286
Rwill_KAE9450996.1
Rphallo_DN643_c0_g1_i9.p1
Rphallo_DN643_c0_g1_i5.p1
Rphallo_DN643_c0_g1_i2.p1
Rphallo_DN643_c0_g1_i12.p1
Rphallo_DN643_c0_g1_i11.p1
Rphallo_DN643_c0_g1_i1.p3
Rphallo_DN643_c0_g1
Pequestris_020572531.1
Mesc_14G104300.3.p
Mesc_14G104300.3
Mesc_06G066500.1.p
Mesc_06G066500.1
Hmono_32757_g1_i1.p2
Eroseum_416659_g2_i4.p1
Eroseum_416659_g2_i3.p1
Eroseum_416659_g2_i2.p1
Eroseum_416659_g2
Eaphy_278901_g2_i3.p1
Eaphy_278901_g2_i1.p1
Eaphy_278901_g2
Bfungosa_DN49897_c0_g1_i1.p1
Bfungosa_DN49897_c0_g1
Bfungosa_cNGB_BALA00440
Bfungosa_CNCB_GWHPDONK000356
Athal_AT1G25350.2
Athal_AT1G25350.1
Athal_AT1G25350
LOCALIZER
TargetP
Algorithm
LOCALIZER
TargetP
Algorithm
Mitochon. Targeting
0.00 0.25 0.50 0.75 1.00
Plastid Targeting
0.00 0.25 0.50 0.75 1.00

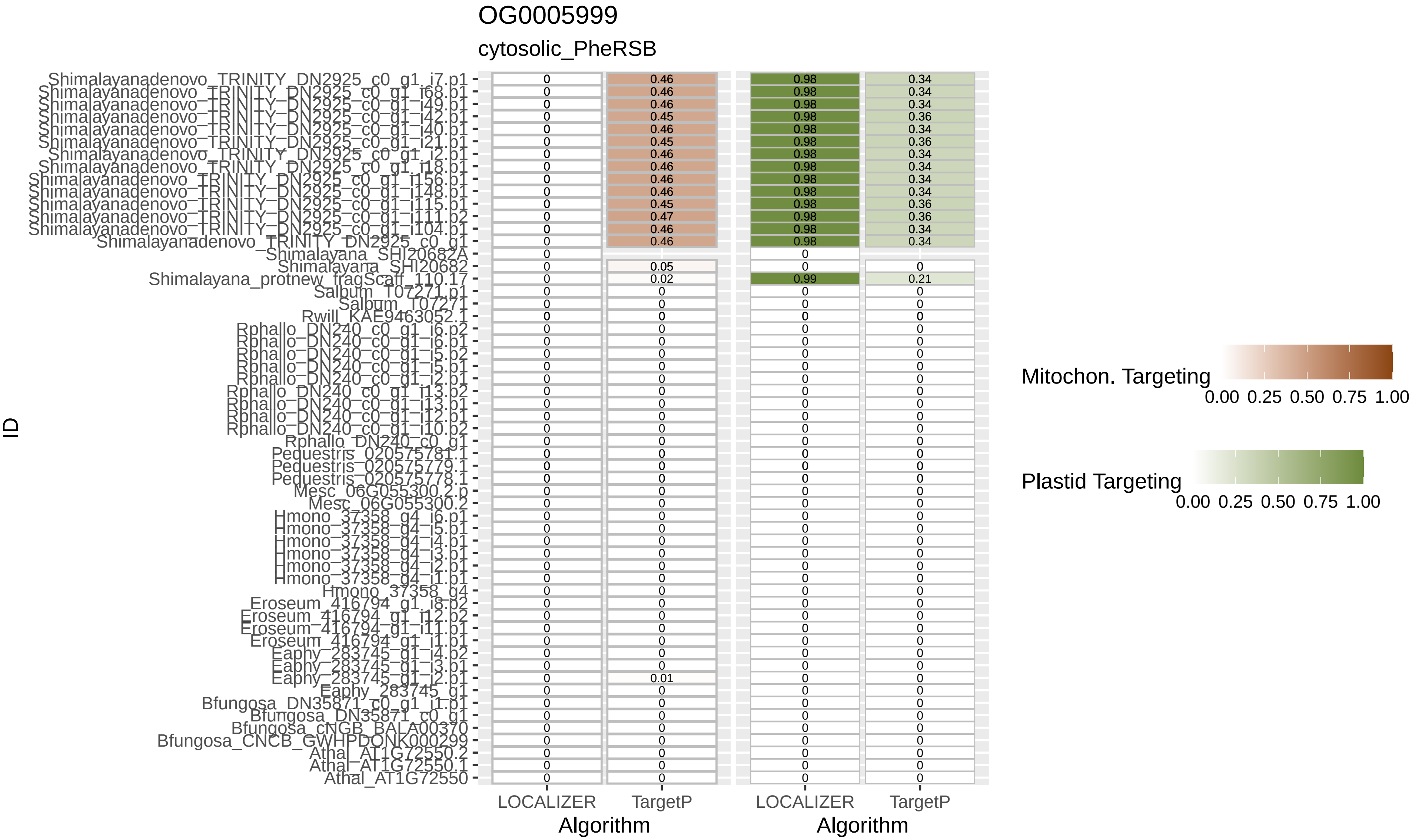
OG0005999
cytosolic_PheRSB
ID
Shimalayanadenovo_TRINITY_DN2925_c0_g1_i7.p1
Shimalayanadenovo_TRINITY_DN2925_c0_g1_i68.p1
Shimalayanadenovo_TRINITY_DN2925_c0_g1_i49.p1
Shimalayanadenovo_TRINITY_DN2925_c0_g1_i42.p1
Shimalayanadenovo_TRINITY_DN2925_c0_g1_i40.p1
Shimalayanadenovo_TRINITY_DN2925_c0_g1_i21.p1
Shimalayanadenovo_TRINITY_DN2925_c0_g1_i2.p1
Shimalayanadenovo_TRINITY_DN2925_c0_g1_i18.p1
Shimalayanadenovo_TRINITY_DN2925_c0_g1_i156.p1
Shimalayanadenovo_TRINITY_DN2925_c0_g1_i148.p1
Shimalayanadenovo_TRINITY_DN2925_c0_g1_i115.p1
Shimalayanadenovo_TRINITY_DN2925_c0_g1_i111.p2
Shimalayanadenovo_TRINITY_DN2925_c0_g1_i104.p1
Shimalayanadenovo_TRINITY_DN2925_c0_g1
Shimalayana_SHI20682A
Shimalayana_SHI20682
Shimalayana_protnew_fragScaff_110.17
Salbum_T07271.p1
Salbum_T07271
Rwill_KAE9463052.1
Rphallo_DN240_c0_g1_i6.p2
Rphallo_DN240_c0_g1_i6.p1
Rphallo_DN240_c0_g1_i5.p2
Rphallo_DN240_c0_g1_i5.p1
Rphallo_DN240_c0_g1_i2.p1
Rphallo_DN240_c0_g1_i13.p2
Rphallo_DN240_c0_g1_i13.p1
Rphallo_DN240_c0_g1_i12.p1
Rphallo_DN240_c0_g1_i10.p2
Rphallo_DN240_c0_g1
Pequestris_020575781.1
Pequestris_020575779.1
Pequestris_020575778.1
Mesc_06G055300.2.p
Mesc_06G055300.2
Hmono_37358_g4_i6.p1
Hmono_37358_g4_i5.p1
Hmono_37358_g4_i4.p1
Hmono_37358_g4_i3.p1
Hmono_37358_g4_i2.p1
Hmono_37358_g4_i1.p1
Hmono_37358_g4
Eroseum_416794_g1_i8.p2
Eroseum_416794_g1_i12.p2
Eroseum_416794_g1_i11.p1
Eroseum_416794_g1_i1.p1
Eaphy_283745_g1_i4.p2
Eaphy_283745_g1_i3.p1
Eaphy_283745_g1_i2.p1
Eaphy_283745_g1
Bfungosa_DN35871_c0_g1_i1.p1
Bfungosa_DN35871_c0_g1
Bfungosa_cNGB_BALA00370
Bfungosa_CNCB_GWHPDONK000299
Athal_AT1G72550.2
Athal_AT1G72550.1
Athal_AT1G72550
LOCALIZER
TargetP
Algorithm
LOCALIZER
TargetP
Algorithm
Mitochon. Targeting
0.00 0.25 0.50 0.75 1.00
Plastid Targeting
0.00 0.25 0.50 0.75 1.00

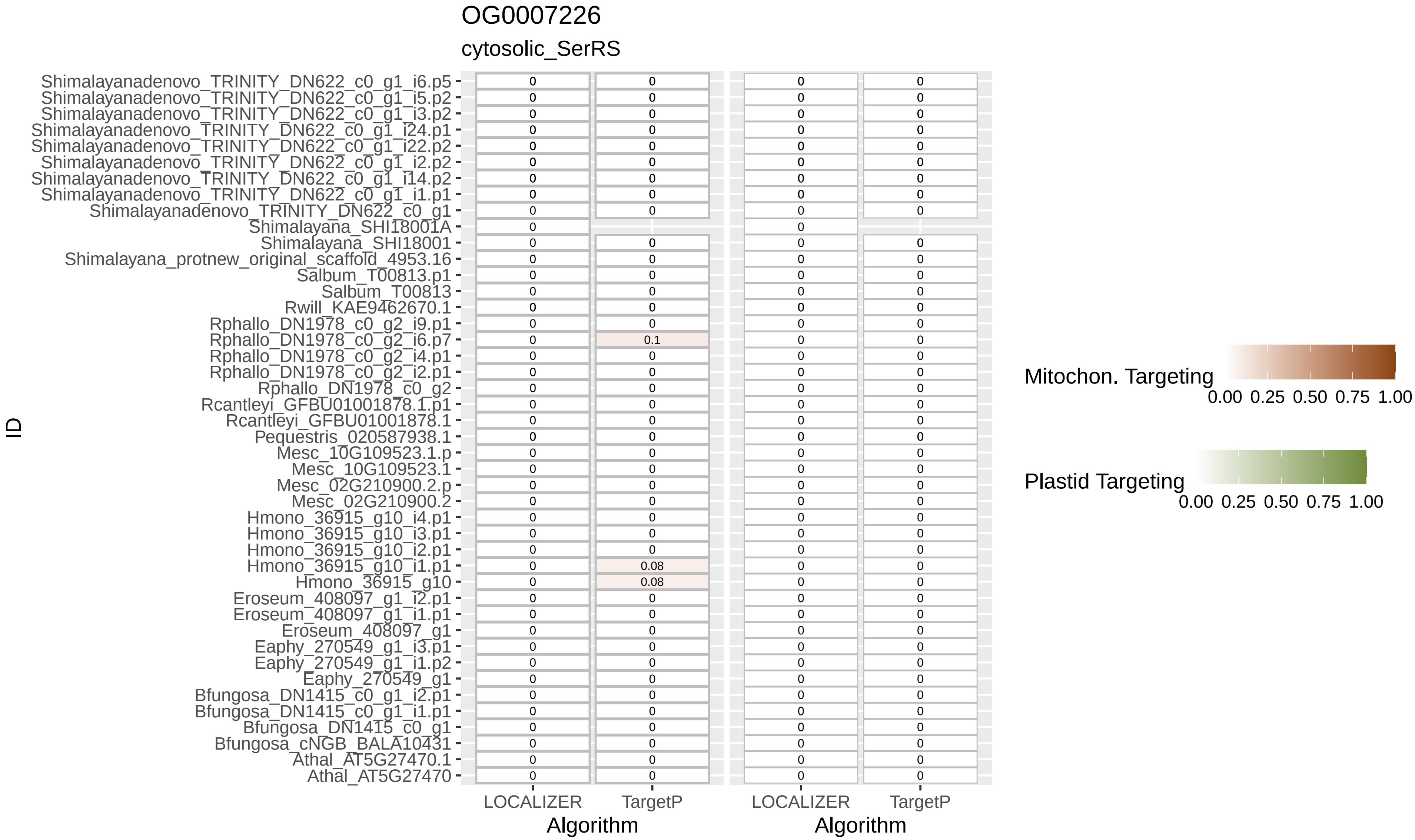

OG0007226
cytosolic_SerRS
ID
Shimalayanadenovo_TRINITY_DN622_c0_g1_i6.p5
Shimalayanadenovo_TRINITY_DN622_c0_g1_i5.p2
Shimalayanadenovo_TRINITY_DN622_c0_g1_i3.p2
Shimalayanadenovo_TRINITY_DN622_c0_g1_i24.p1
Shimalayanadenovo_TRINITY_DN622_c0_g1_i22.p2
Shimalayanadenovo_TRINITY_DN622_c0_g1_i2.p2
Shimalayanadenovo_TRINITY_DN622_c0_g1_i14.p2
Shimalayanadenovo_TRINITY_DN622_c0_g1_i1.p1
Shimalayanadenovo_TRINITY_DN622_c0_g1
Shimalayana_SHI18001A
Shimalayana_SHI18001
Shimalayana_protnew_original_scaffold_4953.16
Salbum_T00813.p1
Salbum_T00813
Rwill_KAE9462670.1
Rphallo_DN1978_c0_g2_i9.p1
Rphallo_DN1978_c0_g2_i6.p7
Rphallo_DN1978_c0_g2_i4.p1
Rphallo_DN1978_c0_g2_i2.p1
Rphallo_DN1978_c0_g2
Rcantleyi_GFBU01001878.1.p1
Rcantleyi_GFBU01001878.1
Pequestris_020587938.1
Mesc_10G109523.1.p
Mesc_10G109523.1
Mesc_02G210900.2.p
Mesc_02G210900.2
Hmono_36915_g10_i4.p1
Hmono_36915_g10_i3.p1
Hmono_36915_g10_i2.p1
Hmono_36915_g10_i1.p1
Hmono_36915_g10
Eroseum_408097_g1_i2.p1
Eroseum_408097_g1_i1.p1
Eroseum_408097_g1
Eaphy_270549_g1_i3.p1
Eaphy_270549_g1_i1.p2
Eaphy_270549_g1
Bfungosa_DN1415_c0_g1_i2.p1
Bfungosa_DN1415_c0_g1_i1.p1
Bfungosa_DN1415_c0_g1
Bfungosa_cNGB_BALA10431
Athal_AT5G27470.1
Athal_AT5G27470
0.1
0.08
0.08
LOCALIZER
TargetP
Algorithm
LOCALIZER
TargetP
Algorithm
Mitochon. Targeting
0.00 0.25 0.50 0.75 1.00
Plastid Targeting
0.00 0.25 0.50 0.75 1.00

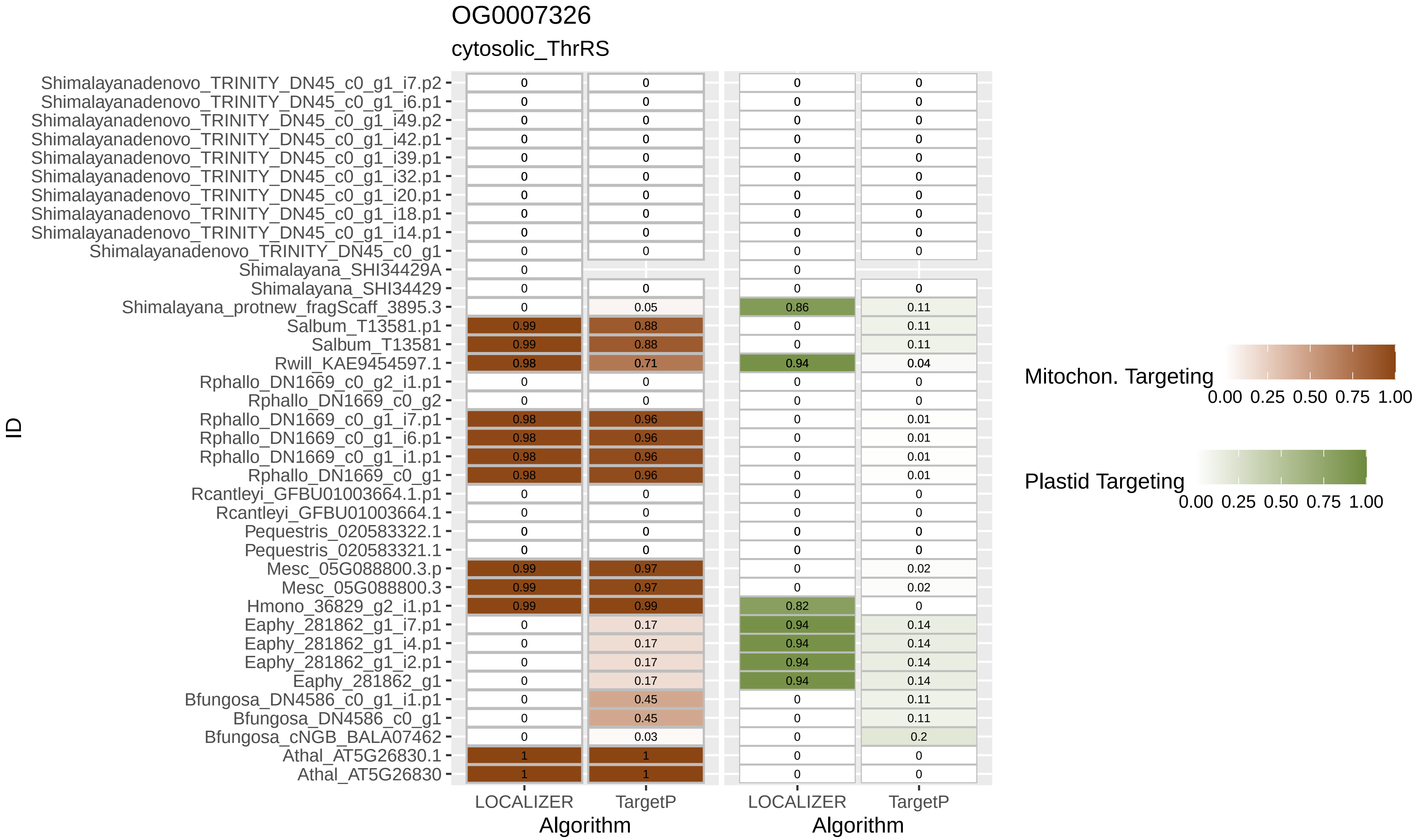
OG0007326
cytosolic_ThrRS
ID
Shimalayanadenovo_TRINITY_DN45_c0_g1_i7.p2
Shimalayanadenovo_TRINITY_DN45_c0_g1_i6.p1
Shimalayanadenovo_TRINITY_DN45_c0_g1_i49.p2
Shimalayanadenovo_TRINITY_DN45_c0_g1_i42.p1
Shimalayanadenovo_TRINITY_DN45_c0_g1_i39.p1
Shimalayanadenovo_TRINITY_DN45_c0_g1_i32.p1
Shimalayanadenovo_TRINITY_DN45_c0_g1_i20.p1
Shimalayanadenovo_TRINITY_DN45_c0_g1_i18.p1
Shimalayanadenovo_TRINITY_DN45_c0_g1_i14.p1
Shimalayanadenovo_TRINITY_DN45_c0_g1
Shimalayana_SHI34429A
Shimalayana_SHI34429
Shimalayana_protnew_fragScaff_3895.3
Salbum_T13581.p1
Salbum_T13581
Rwill_KAE9454597.1
Rphallo_DN1669_c0_g2_i1.p1
Rphallo_DN1669_c0_g2
Rphallo_DN1669_c0_g1_i7.p1
Rphallo_DN1669_c0_g1_i6.p1
Rphallo_DN1669_c0_g1_i1.p1
Rphallo_DN1669_c0_g1
Rcantleyi_GFBU01003664.1.p1
Rcantleyi_GFBU01003664.1
Pequestris_020583322.1
Pequestris_020583321.1
Mesc_05G088800.3.p
Mesc_05G088800.3
Hmono_36829_g2_i1.p1
Eaphy_281862_g1_i7.p1
Eaphy_281862_g1_i4.p1
Eaphy_281862_g1_i2.p1
Eaphy_281862_g1
Bfungosa_DN4586_c0_g1_i1.p1
Bfungosa_DN4586_c0_g1
Bfungosa_cNGB_BALA07462
Athal_AT5G26830.1
Athal_AT5G26830
LOCALIZER
TargetP
Algorithm
LOCALIZER
TargetP
Algorithm
Mitochon. Targeting
0.00 0.25 0.50 0.75 1.00
Plastid Targeting
0.00 0.25 0.50 0.75 1.00
0.05
0.99 0.88
0.99 0.88
0.98 0.71
0.98 0.96
0.98 0.96
0.98 0.96
0.98 0.96
0.99 0.97
0.99 0.97
0.99 0.99
0.17
0.17
0.17
0.17
0.45
0.45
0.03
1 1
1 1
0.86 0.11
0.11
0.11
0.94 0.04
0.01
0.01
0.01
0.01
0.02
0.02
0.82
0.94 0.14
0.94 0.14
0.94 0.14
0.94 0.14
0.11
0.11
0.2

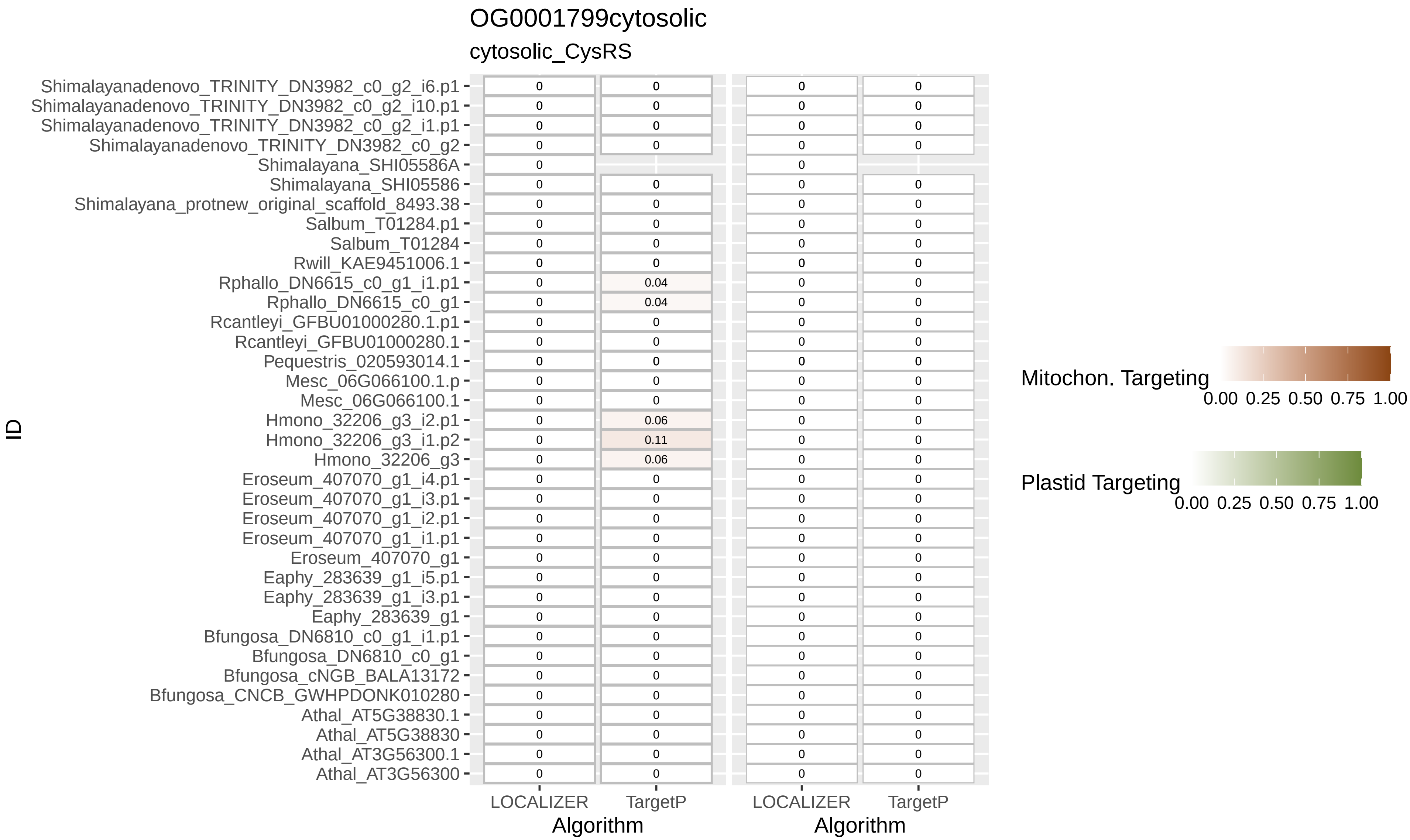

OG0001799cytosolic
cytosolic_CysRS
ID
Shimalayanadenovo_TRINITY_DN3982_c0_g2_i6.p1
Shimalayanadenovo_TRINITY_DN3982_c0_g2_i10.p1
Shimalayanadenovo_TRINITY_DN3982_c0_g2_i1.p1
Shimalayanadenovo_TRINITY_DN3982_c0_g2
Shimalayana_SHI05586A
Shimalayana_SHI05586
Shimalayana_protnew_original_scaffold_8493.38
Salbum_T01284.p1
Salbum_T01284
Rwill_KAE9451006.1
Rphallo_DN6615_c0_g1_i1.p1
Rphallo_DN6615_c0_g1
Rcantleyi_GFBU01000280.1.p1
Rcantleyi_GFBU01000280.1
Pequestris_020593014.1
Mesc_06G066100.1.p
Mesc_06G066100.1
Hmono_32206_g3_i2.p1
Hmono_32206_g3_i1.p2
Hmono_32206_g3
Eroseum_407070_g1_i4.p1
Eroseum_407070_g1_i3.p1
Eroseum_407070_g1_i2.p1
Eroseum_407070_g1_i1.p1
Eroseum_407070_g1
Eaphy_283639_g1_i5.p1
Eaphy_283639_g1_i3.p1
Eaphy_283639_g1
Bfungosa_DN6810_c0_g1_i1.p1
Bfungosa_DN6810_c0_g1
Bfungosa_cNGB_BALA13172
Bfungosa_CNCB_GWHPDONK010280
Athal_AT5G38830.1
Athal_AT5G38830
Athal_AT3G56300.1
Athal_AT3G56300
0.04
0.04
0.06
0.11
0.06
LOCALIZER
TargetP
Algorithm
LOCALIZER
TargetP
Algorithm
Mitochon. Targeting
0.00 0.25 0.50 0.75 1.00
Plastid Targeting
0.00 0.25 0.50 0.75 1.00

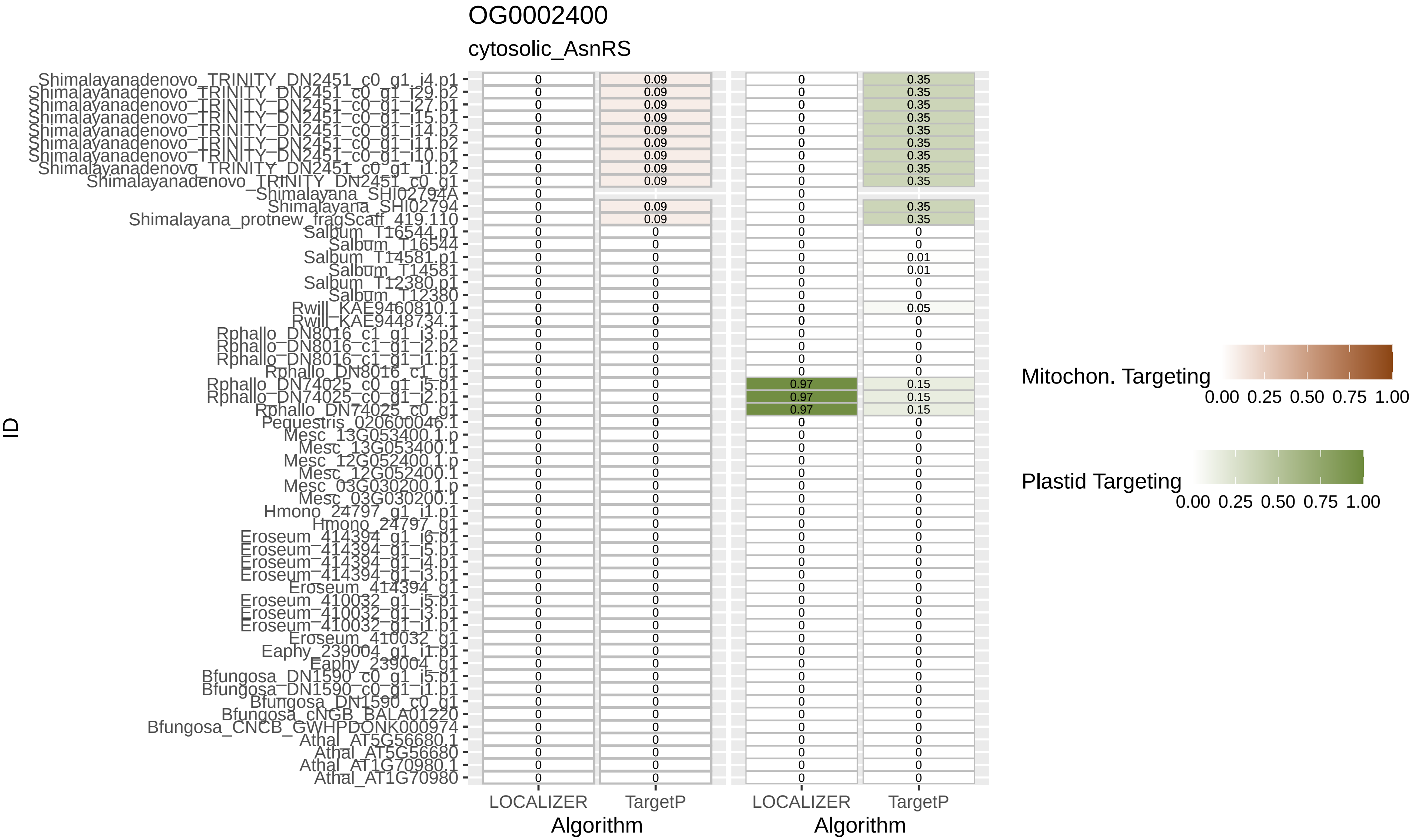
OG0002400
cytosolic_AsnRS
Shimalayanadenovo_TRINITY_DN2451_c0_g1_i4.p1
Shimalayanadenovo_TRINITY_DN2451_c0_g1_i29.p2
Shimalayanadenovo_TRINITY_DN2451_c0_g1_i27.p1
Shimalayanadenovo_TRINITY_DN2451_c0_g1_i15.p1
Shimalayanadenovo_TRINITY_DN2451_c0_g1_i14.p2
Shimalayanadenovo_TRINITY_DN2451_c0_g1_i11.p2
Shimalayanadenovo_TRINITY_DN2451_c0_g1_i10.p1
Shimalayanadenovo_TRINITY_DN2451_c0_g1_i1.p2
Shimalayanadenovo_TRINITY_DN2451_c0_g1
Shimalayana_SHI02794A
Shimalayana_SHI02794
Shimalayana_protnew_fragScaff_419.110
Salbum_T16544.p1
Salbum_T16544
Salbum_T14581.p1
Salbum_T14581
Salbum_T12380.p1
Salbum_T12380
Rwill_KAE9460810.1
Rwill_KAE9448734.1
Rphallo_DN8016_c1_g1_i3.p1
Rphallo_DN8016_c1_g1_i2.p2
Rphallo_DN8016_c1_g1_i1.p1
Rphallo_DN8016_c1_g1
Rphallo_DN74025_c0_g1_i5.p1
Rphallo_DN74025_c0_g1_i2.p1
Rphallo_DN74025_c0_g1
Pequestris_020600046.1
Mesc_13G053400.1.p
Mesc_13G053400.1
Mesc_12G052400.1.p
Mesc_12G052400.1
Mesc_03G030200.1.p
Mesc_03G030200.1
Hmono_24797_g1_i1.p1
Hmono_24797_g1
Eroseum_414394_g1_i6.p1
Eroseum_414394_g1_i5.p1
Eroseum_414394_g1_i4.p1
Eroseum_414394_g1_i3.p1
Eroseum_414394_g1
Eroseum_410032_g1_i5.p1
Eroseum_410032_g1_i3.p1
Eroseum_410032_g1_i1.p1
Eroseum_410032_g1
Eaphy_239004_g1_i1.p1
Eaphy_239004_g1
Bfungosa_DN1590_c0_g1_i5.p1
Bfungosa_DN1590_c0_g1_i1.p1
Bfungosa_DN1590_c0_g1
Bfungosa_cNGB_BALA01220
Bfungosa_CNCB_GWHPDONK000974
Athal_AT5G56680.1
Athal_AT5G56680
Athal_AT1G70980.1
Athal_AT1G70980
ID
LOCALIZER
TargetP
Algorithm
LOCALIZER
TargetP
Algorithm
Mitochon. Targeting
0.00 0.25 0.50 0.75 1.00
Plastid Targeting
0.00 0.25 0.50 0.75 1.00

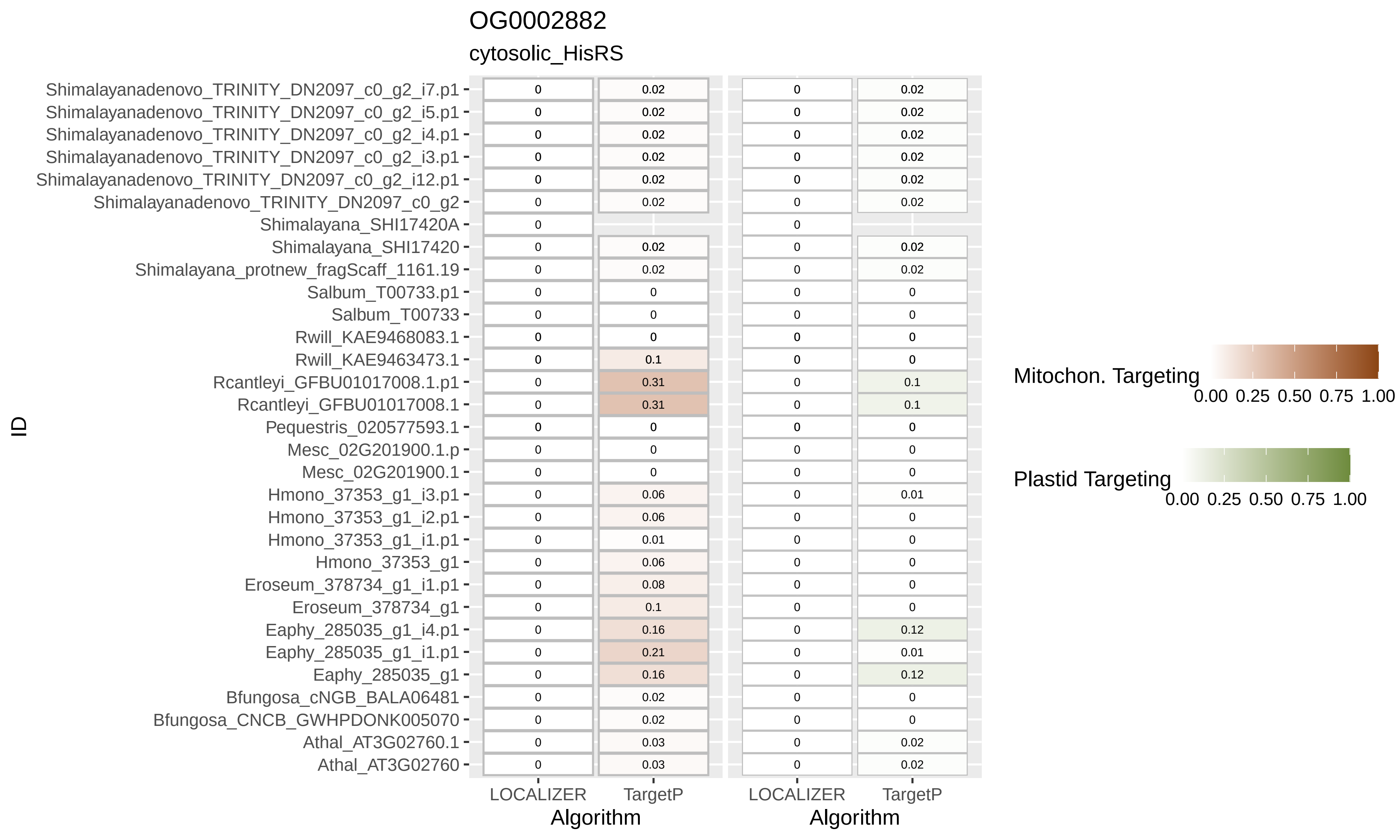
OG0002882
cytosolic_HisRS
ID
Shimalayanadenovo_TRINITY_DN2097_c0_g2_i7.p1 0 0.02 0 0.02
Shimalayanadenovo_TRINITY_DN2097_c0_g2_i5.p1 0 0.02 0 0.02
Shimalayanadenovo_TRINITY_DN2097_c0_g2_i4.p1 0 0.02 0 0.02
Shimalayanadenovo_TRINITY_DN2097_c0_g2_i3.p1 0 0.02 0 0.02
Shimalayanadenovo_TRINITY_DN2097_c0_g2_i12.p1 0 0.02 0 0.02
Shimalayanadenovo_TRINITY_DN2097_c0_g2 0 0.02 0 0.02
Shimalayana_SHI17420A 0 0
Shimalayana_SHI17420 0 0.02 0 0.02
Shimalayana_protnew_fragScaff_1161.19 0 0.02 0 0.02
Salbum_T00733.p1 0 0 0 0
Salbum_T00733 0 0 0 0
Rwill_KAE9468083.1 0 0 0 0
Rwill_KAE9463473.1 0 0.1 0 0
Rcantleyi_GFBU01017008.1.p1 0 0.31 0 0.1
Rcantleyi_GFBU01017008.1 0 0.31 0 0.1
Pequestris_020577593.1 0 0 0 0
Mesc_02G201900.1.p 0 0 0 0
Mesc_02G201900.1 0 0 0 0
Hmono_37353_g1_i3.p1 0 0.06 0 0.01
Hmono_37353_g1_i2.p1 0 0.06 0 0
Hmono_37353_g1_i1.p1 0 0.01 0 0
Hmono_37353_g1 0 0.06 0 0
Eroseum_378734_g1_i1.p1 0 0.08 0 0
Eroseum_378734_g1 0 0.1 0 0
Eaphy_285035_g1_i4.p1 0 0.16 0 0.12
Eaphy_285035_g1_i1.p1 0 0.21 0 0.01
Eaphy_285035_g1 0 0.16 0 0.12
Bfungosa_cNGB_BALA06481 0 0.02 0 0
Bfungosa_CNCB_GWHPDONK005070 0 0.02 0 0
Athal_AT3G02760.1 0 0.03 0 0.02
Athal_AT3G02760 0 0.03 0 0.02
LOCALIZER TargetP
Algorithm
LOCALIZER TargetP
Algorithm
Mitochon. Targeting
0.00 0.25 0.50 0.75 1.00
Plastid Targeting
0.00 0.25 0.50 0.75 1.00

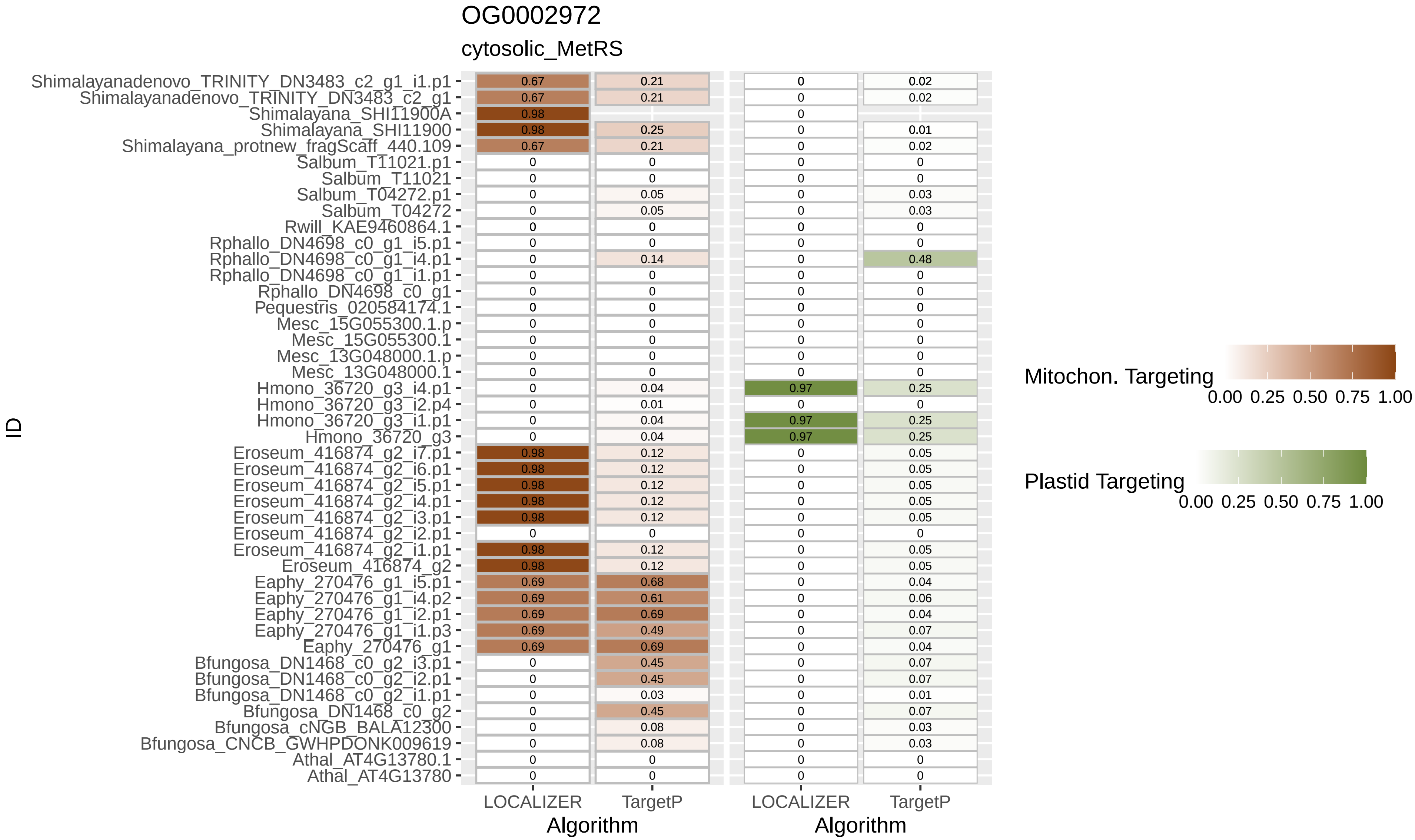

OG0002972
cytosolic_MetRS
ID
Algorithm
LOCALIZER
TargetP
Mitochon. Targeting
Plastid Targeting
0.00 0.25 0.50 0.75 1.00
Shimalayanadenovo_TRINITY_DN3483_c2_g1_i1.p1
Shimalayanadenovo_TRINITY_DN3483_c2_g1
Shimalayana_SHI11900A
Shimalayana_SHI11900
Shimalayana_protnew_fragScaff_440.109
Salbum_T11021.p1
Salbum_T11021
Salbum_T04272.p1
Salbum_T04272
Rwill_KAE9460864.1
Rphallo_DN4698_c0_g1_i5.p1
Rphallo_DN4698_c0_g1_i4.p1
Rphallo_DN4698_c0_g1_i1.p1
Rphallo_DN4698_c0_g1
Pequestris_020584174.1
Mesc_15G055300.1.p
Mesc_15G055300.1
Mesc_13G048000.1.p
Mesc_13G048000.1
Hmono_36720_g3_i4.p1
Hmono_36720_g3_i2.p4
Hmono_36720_g3_i1.p1
Hmono_36720_g3
Eroseum_416874_g2_i7.p1
Eroseum_416874_g2_i6.p1
Eroseum_416874_g2_i5.p1
Eroseum_416874_g2_i4.p1
Eroseum_416874_g2_i3.p1
Eroseum_416874_g2_i2.p1
Eroseum_416874_g2_i1.p1
Eroseum_416874_g2
Eaphy_270476_g1_i5.p1
Eaphy_270476_g1_i4.p2
Eaphy_270476_g1_i2.p1
Eaphy_270476_g1_i1.p3
Eaphy_270476_g1
Bfungosa_DN1468_c0_g2_i3.p1
Bfungosa_DN1468_c0_g2_i2.p1
Bfungosa_DN1468_c0_g2_i1.p1
Bfungosa_DN1468_c0_g2
Bfungosa_cNGB_BALA12300
Bfungosa_CNCB_GWHPDONK009619
Athal_AT4G13780.1
Athal_AT4G13780

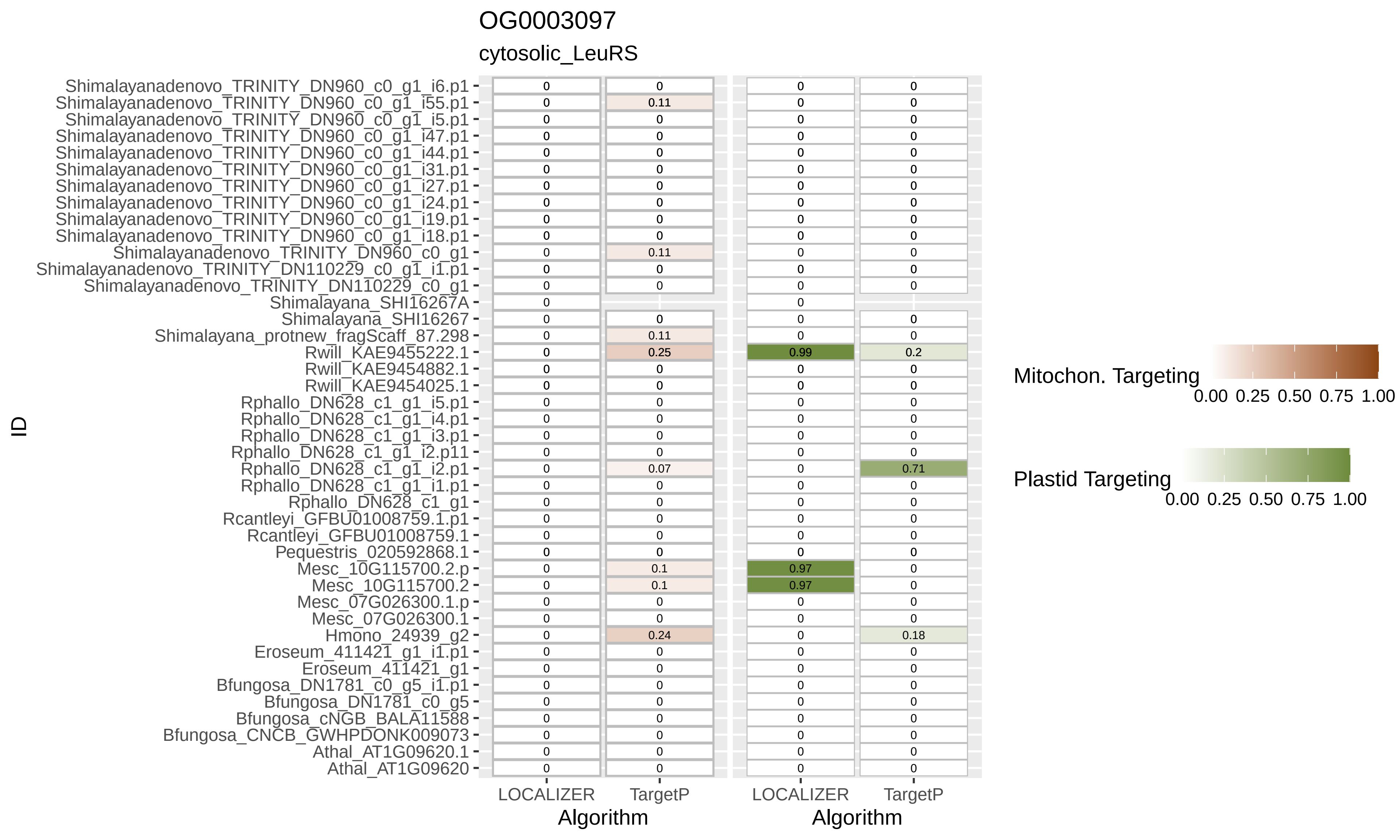
OG0003097
cytosolic_LeuRS
ID
Shimalayanadenovo_TRINITY_DN960_c0_g1_i6.p1 0 0 0 0
Shimalayanadenovo_TRINITY_DN960_c0_g1_i55.p1 0 0.11 0 0
Shimalayanadenovo_TRINITY_DN960_c0_g1_i5.p1 0 0 0 0
Shimalayanadenovo_TRINITY_DN960_c0_g1_i47.p1 0 0 0 0
Shimalayanadenovo_TRINITY_DN960_c0_g1_i44.p1 0 0 0 0
Shimalayanadenovo_TRINITY_DN960_c0_g1_i31.p1 0 0 0 0
Shimalayanadenovo_TRINITY_DN960_c0_g1_i27.p1 0 0 0 0
Shimalayanadenovo_TRINITY_DN960_c0_g1_i24.p1 0 0 0 0
Shimalayanadenovo_TRINITY_DN960_c0_g1_i19.p1 0 0 0 0
Shimalayanadenovo_TRINITY_DN960_c0_g1_i18.p1 0 0 0 0
Shimalayanadenovo_TRINITY_DN960_c0_g1 0 0.11 0 0
Shimalayanadenovo_TRINITY_DN110229_c0_g1_i1.p1 0 0 0 0
Shimalayanadenovo_TRINITY_DN110229_c0_g1 0 0 0 0
Shimalayana_SHI16267A 0 0
Shimalayana_SHI16267 0 0 0 0
Shimalayana_protnew_fragScaff_87.298 0 0.11 0 0
Rwill_KAE9455222.1 0 0.25 0.99 0.2
Rwill_KAE9454882.1 0 0 0 0
Rwill_KAE9454025.1 0 0 0 0
Rphallo_DN628_c1_g1_i5.p1 0 0 0 0
Rphallo_DN628_c1_g1_i4.p1 0 0 0 0
Rphallo_DN628_c1_g1_i3.p1 0 0 0 0
Rphallo_DN628_c1_g1_i2.p11 0 0 0 0
Rphallo_DN628_c1_g1_i2.p1 0 0.07 0 0.71
Rphallo_DN628_c1_g1_i1.p1 0 0 0 0
Rphallo_DN628_c1_g1 0 0 0 0
Rcantleyi_GFBU01008759.1.p1 0 0 0 0
Rcantleyi_GFBU01008759.1 0 0 0 0
Pequestris_020592868.1 0 0 0 0
Mesc_10G115700.2.p 0 0.1 0.97 0
Mesc_10G115700.2 0 0.1 0.97 0
Mesc_07G026300.1.p 0 0 0 0
Mesc_07G026300.1 0 0 0 0
Hmono_24939_g2 0 0.24 0 0.18
Eroseum_411421_g1_i1.p1 0 0 0 0
Eroseum_411421_g1 0 0 0 0
Bfungosa_DN1781_c0_g5_i1.p1 0 0 0 0
Bfungosa_DN1781_c0_g5 0 0 0 0
Bfungosa_cNGB_BALA11588 0 0 0 0
Bfungosa_CNCB_GWHPDONK009073 0 0 0 0
Athal_AT1G09620.1 0 0 0 0
Athal_AT1G09620 0 0 0 0
LOCALIZER TargetP
Algorithm
LOCALIZER TargetP
Algorithm
Mitochon. Targeting
0.00 0.25 0.50 0.75 1.00
Plastid Targeting
0.00 0.25 0.50 0.75 1.00

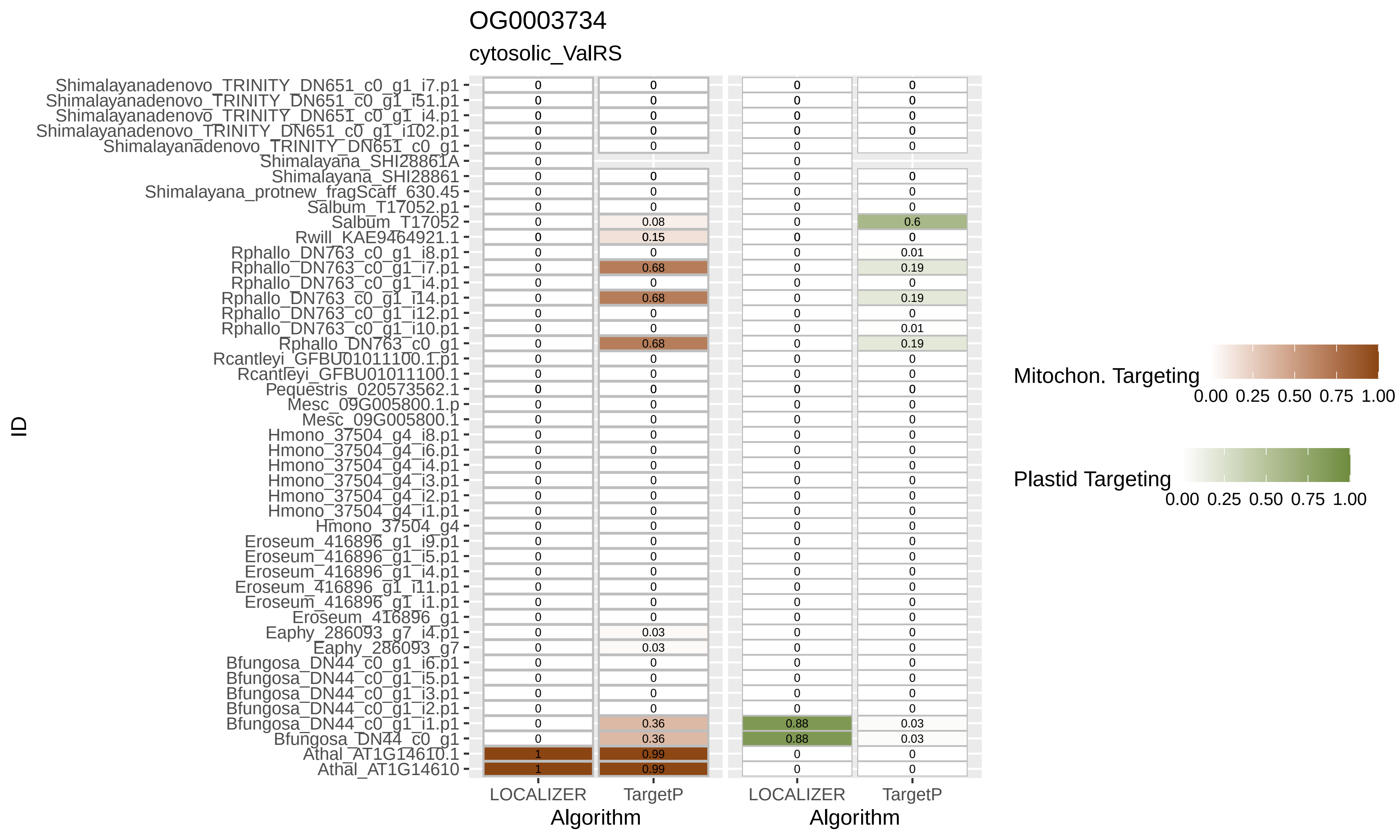
OG0003734
cytosolic_ValRS
ID
Algorithm
LOCALIZER
TargetP
Mitochon. Targeting
Plastid Targeting
0.00 0.25 0.50 0.75 1.00
Shimalayanadenovo_TRINITY_DN651_c0_g1_i7.p1
Shimalayanadenovo_TRINITY_DN651_c0_g1_i51.p1
Shimalayanadenovo_TRINITY_DN651_c0_g1_i4.p1
Shimalayanadenovo_TRINITY_DN651_c0_g1_i102.p1
Shimalayanadenovo_TRINITY_DN651_c0_g1
Shimalayana_SHI28861A
Shimalayana_SHI28861
Shimalayana_protnew_fragScaff_630.45
Salbum_T17052.p1
Salbum_T17052
Rwill_KAE9464921.1
Rphallo_DN763_c0_g1_i8.p1
Rphallo_DN763_c0_g1_i7.p1
Rphallo_DN763_c0_g1_i4.p1
Rphallo_DN763_c0_g1_i14.p1
Rphallo_DN763_c0_g1_i12.p1
Rphallo_DN763_c0_g1_i10.p1
Rphallo_DN763_c0_g1
Rcantleyi_GFBU01011100.1.p1
Rcantleyi_GFBU01011100.1
Pequestris_020573562.1
Mesc_09G005800.1.p
Mesc_09G005800.1
Hmono_37504_g4_i8.p1
Hmono_37504_g4_i6.p1
Hmono_37504_g4_i4.p1
Hmono_37504_g4_i3.p1
Hmono_37504_g4_i2.p1
Hmono_37504_g4_i1.p1
Hmono_37504_g4
Eroseum_416896_g1_i9.p1
Eroseum_416896_g1_i5.p1
Eroseum_416896_g1_i4.p1
Eroseum_416896_g1_i11.p1
Eroseum_416896_g1_i1.p1
Eroseum_416896_g1
Eaphy_286093_g7_i4.p1
Eaphy_286093_g7
Bfungosa_DN44_c0_g1_i6.p1
Bfungosa_DN44_c0_g1_i5.p1
Bfungosa_DN44_c0_g1_i3.p1
Bfungosa_DN44_c0_g1_i2.p1
Bfungosa_DN44_c0_g1_i1.p1
Bfungosa_DN44_c0_g1
Athal_AT1G14610.1
Athal_AT1G14610

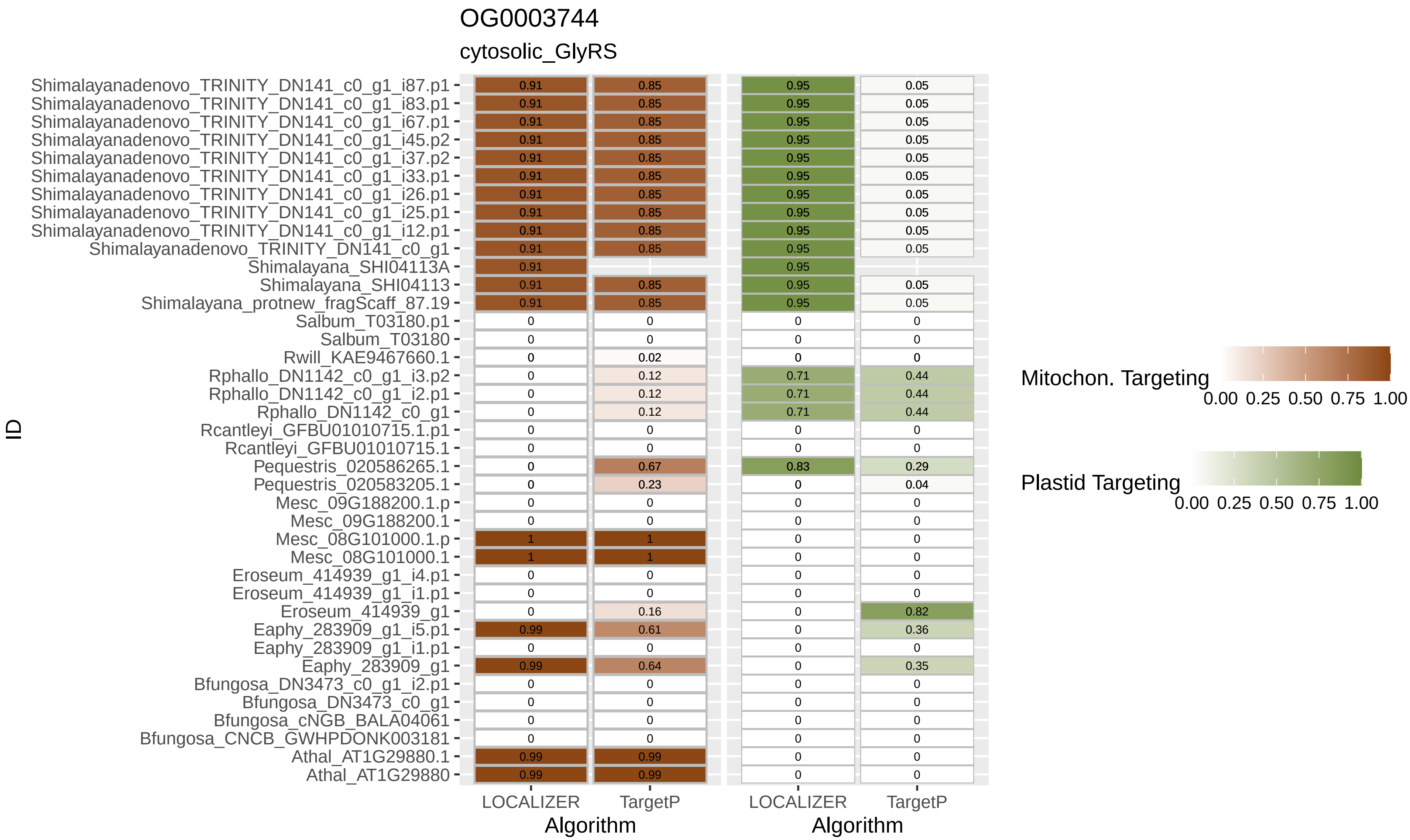

OG0003744
cytosolic_GlyRS
ID
Algorithm
LOCALIZER
TargetP
Mitochon. Targeting
Plastid Targeting
0.00 0.25 0.50 0.75 1.00
Shimalayanadenovo_TRINITY_DN141_c0_g1_i87.p1
Shimalayanadenovo_TRINITY_DN141_c0_g1_i83.p1
Shimalayanadenovo_TRINITY_DN141_c0_g1_i67.p1
Shimalayanadenovo_TRINITY_DN141_c0_g1_i45.p2
Shimalayanadenovo_TRINITY_DN141_c0_g1_i37.p2
Shimalayanadenovo_TRINITY_DN141_c0_g1_i33.p1
Shimalayanadenovo_TRINITY_DN141_c0_g1_i26.p1
Shimalayanadenovo_TRINITY_DN141_c0_g1_i25.p1
Shimalayanadenovo_TRINITY_DN141_c0_g1_i12.p1
Shimalayanadenovo_TRINITY_DN141_c0_g1
Shimalayana_SHI04113A
Shimalayana_SHI04113
Shimalayana_protnew_fragScaff_87.19
Salbum_T03180.p1
Salbum_T03180
Rwill_KAE9467660.1
Rphallo_DN1142_c0_g1_i3.p2
Rphallo_DN1142_c0_g1_i2.p1
Rphallo_DN1142_c0_g1
Rcantleyi_GFBU01010715.1.p1
Rcantleyi_GFBU01010715.1
Pequestris_020586265.1
Pequestris_020583205.1
Mesc_09G188200.1.p
Mesc_09G188200.1
Mesc_08G101000.1.p
Mesc_08G101000.1
Eroseum_414939_g1_i4.p1
Eroseum_414939_g1_i1.p1
Eroseum_414939_g1
Eaphy_283909_g1_i5.p1
Eaphy_283909_g1_i1.p1
Eaphy_283909_g1
Bfungosa_DN3473_c0_g1_i2.p1
Bfungosa_DN3473_c0_g1
Bfungosa_cNGB_BALA04061
Bfungosa_CNCB_GWHPDONK003181
Athal_AT1G29880.1
Athal_AT1G29880

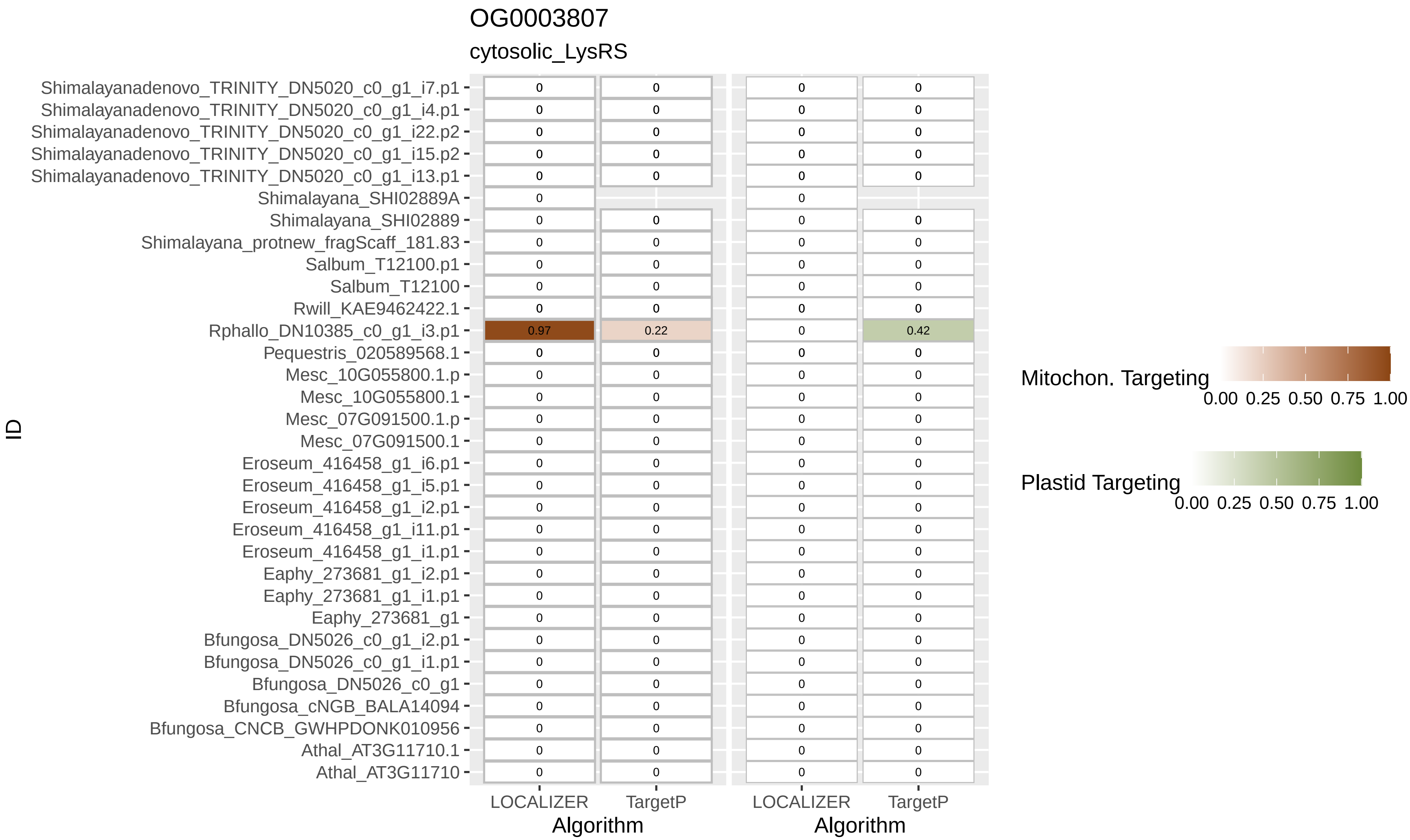
OG0003807
cytosolic_LysRS
Shimalayanadenovo_TRINITY_DN5020_c0_g1_i7.p1
Shimalayanadenovo_TRINITY_DN5020_c0_g1_i4.p1
Shimalayanadenovo_TRINITY_DN5020_c0_g1_i22.p2
Shimalayanadenovo_TRINITY_DN5020_c0_g1_i15.p2
Shimalayanadenovo_TRINITY_DN5020_c0_g1_i13.p1
Shimalayana_SHI02889A
Shimalayana_SHI02889
Shimalayana_protnew_fragScaff_181.83
Salbum_T12100.p1
Salbum_T12100
Rwill_KAE9462422.1
Rphallo_DN10385_c0_g1_i3.p1
Pequestris_020589568.1
Mesc_10G055800.1.p
Mesc_10G055800.1
Mesc_07G091500.1.p
Mesc_07G091500.1
Eroseum_416458_g1_i6.p1
Eroseum_416458_g1_i5.p1
Eroseum_416458_g1_i2.p1
Eroseum_416458_g1_i11.p1
Eroseum_416458_g1_i1.p1
Eaphy_273681_g1_i2.p1
Eaphy_273681_g1_i1.p1
Eaphy_273681_g1
Bfungosa_DN5026_c0_g1_i2.p1
Bfungosa_DN5026_c0_g1_i1.p1
Bfungosa_DN5026_c0_g1
Bfungosa_cNGB_BALA14094
Bfungosa_CNCB_GWHPDONK010956
Athal_AT3G11710.1
Athal_AT3G11710
ID
0.97
0.22
0.42
LOCALIZER
TargetP
Algorithm
LOCALIZER
TargetP
Algorithm
Mitochon. Targeting
0.00 0.25 0.50 0.75 1.00
Plastid Targeting
0.00 0.25 0.50 0.75 1.00

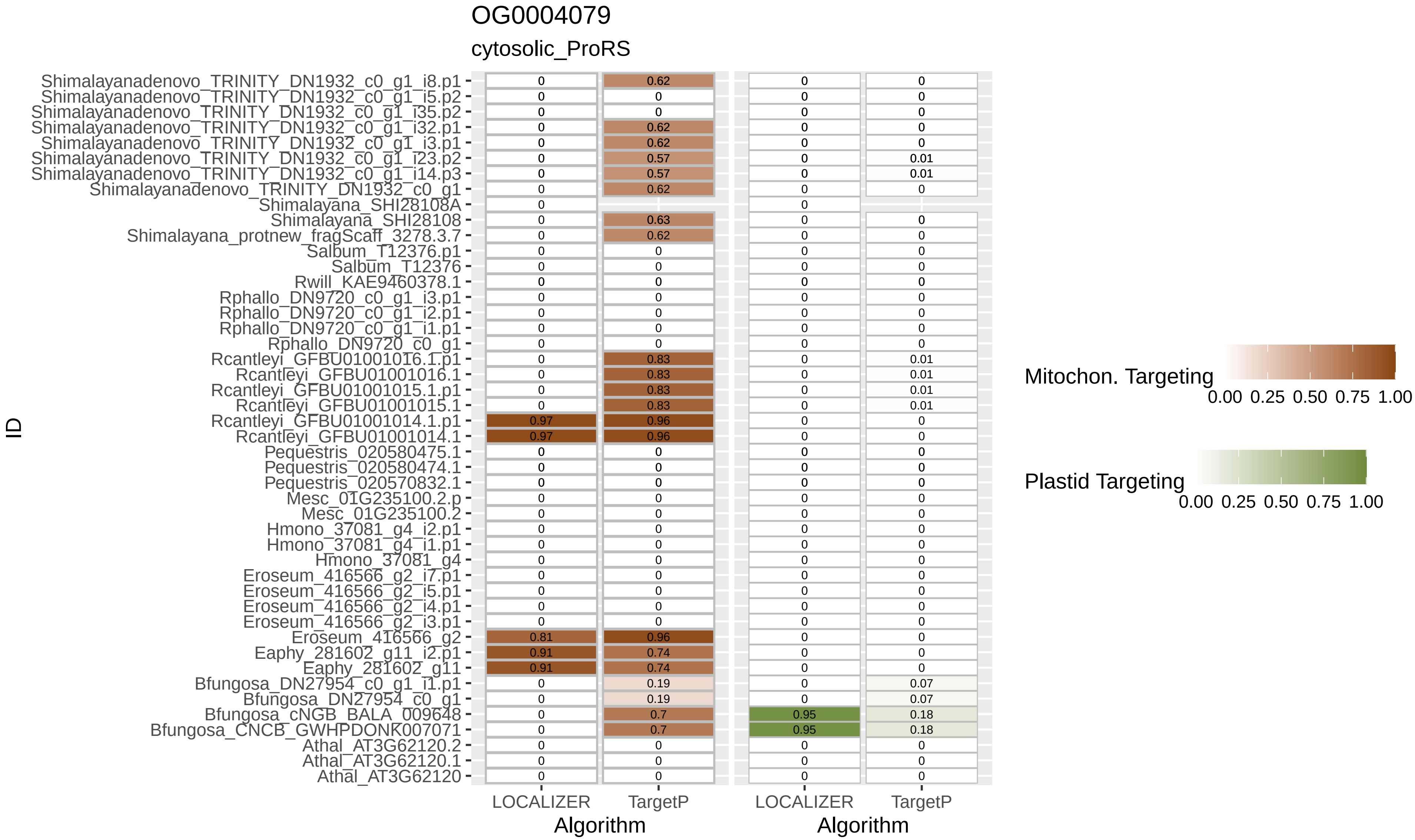
OG0004079
cytosolic_ProRS
ID
Shimalayanadenovo_TRINITY_DN1932_c0_g1_i8.p1 0 0.62 0 0
Shimalayanadenovo_TRINITY_DN1932_c0_g1_i5.p2 0 0 0 0
Shimalayanadenovo_TRINITY_DN1932_c0_g1_i35.p2 0 0 0 0
Shimalayanadenovo_TRINITY_DN1932_c0_g1_i32.p1 0 0.62 0 0
Shimalayanadenovo_TRINITY_DN1932_c0_g1_i3.p1 0 0.62 0 0
Shimalayanadenovo_TRINITY_DN1932_c0_g1_i23.p2 0 0.57 0 0.01
Shimalayanadenovo_TRINITY_DN1932_c0_g1_i14.p3 0 0.57 0 0.01
Shimalayanadenovo_TRINITY_DN1932_c0_g1 0 0.62 0 0
Shimalayana_SHI28108A 0 0
Shimalayana_SHI28108 0 0.63 0 0
Shimalayana_protnew_fragScaff_3278.3.7 0 0.62 0 0
Salbum_T12376.p1 0 0 0 0
Salbum_T12376 0 0 0 0
Rwill_KAE9460378.1 0 0 0 0
Rphallo_DN9720_c0_g1_i3.p1 0 0 0 0
Rphallo_DN9720_c0_g1_i2.p1 0 0 0 0
Rphallo_DN9720_c0_g1_i1.p1 0 0 0 0
Rphallo_DN9720_c0_g1 0 0 0 0
Rcantleyi_GFBU01001016.1.p1 0 0.83 0 0.01
Rcantleyi_GFBU01001016.1 0 0.83 0 0.01
Rcantleyi_GFBU01001015.1.p1 0 0.83 0 0.01
Rcantleyi_GFBU01001015.1 0 0.83 0 0.01
Rcantleyi_GFBU01001014.1.p1 0.97 0.96 0 0
Rcantleyi_GFBU01001014.1 0.97 0.96 0 0
Pequestris_020580475.1 0 0 0 0
Pequestris_020580474.1 0 0 0 0
Pequestris_020570832.1 0 0 0 0
Mesc_01G235100.2.p 0 0 0 0
Mesc_01G235100.2 0 0 0 0
Hmono_37081_g4_i2.p1 0 0 0 0
Hmono_37081_g4_i1.p1 0 0 0 0
Hmono_37081_g4 0 0 0 0
Eroseum_416566_g2_i7.p1 0 0 0 0
Eroseum_416566_g2_i5.p1 0 0 0 0
Eroseum_416566_g2_i4.p1 0 0 0 0
Eroseum_416566_g2_i3.p1 0 0 0 0
Eroseum_416566_g2 0.81 0.96 0 0
Eaphy_281602_g11_i2.p1 0.91 0.74 0 0
Eaphy_281602_g11 0.91 0.74 0 0
Bfungosa_DN27954_c0_g1_i1.p1 0 0.19 0 0.07
Bfungosa_DN27954_c0_g1 0 0.19 0 0.07
Bfungosa_cNGB_BALA_009648 0 0.7 0.95 0.18
Bfungosa_CNCB_GWHPDONK007071 0 0.7 0.95 0.18
Athal_AT3G62120.2 0 0 0 0
Athal_AT3G62120.1 0 0 0 0
Athal_AT3G62120 0 0 0 0
LOCALIZER
TargetP
Algorithm
LOCALIZER
TargetP
Algorithm
Mitochon. Targeting
0.00 0.25 0.50 0.75 1.00
Plastid Targeting
0.00 0.25 0.50 0.75 1.00

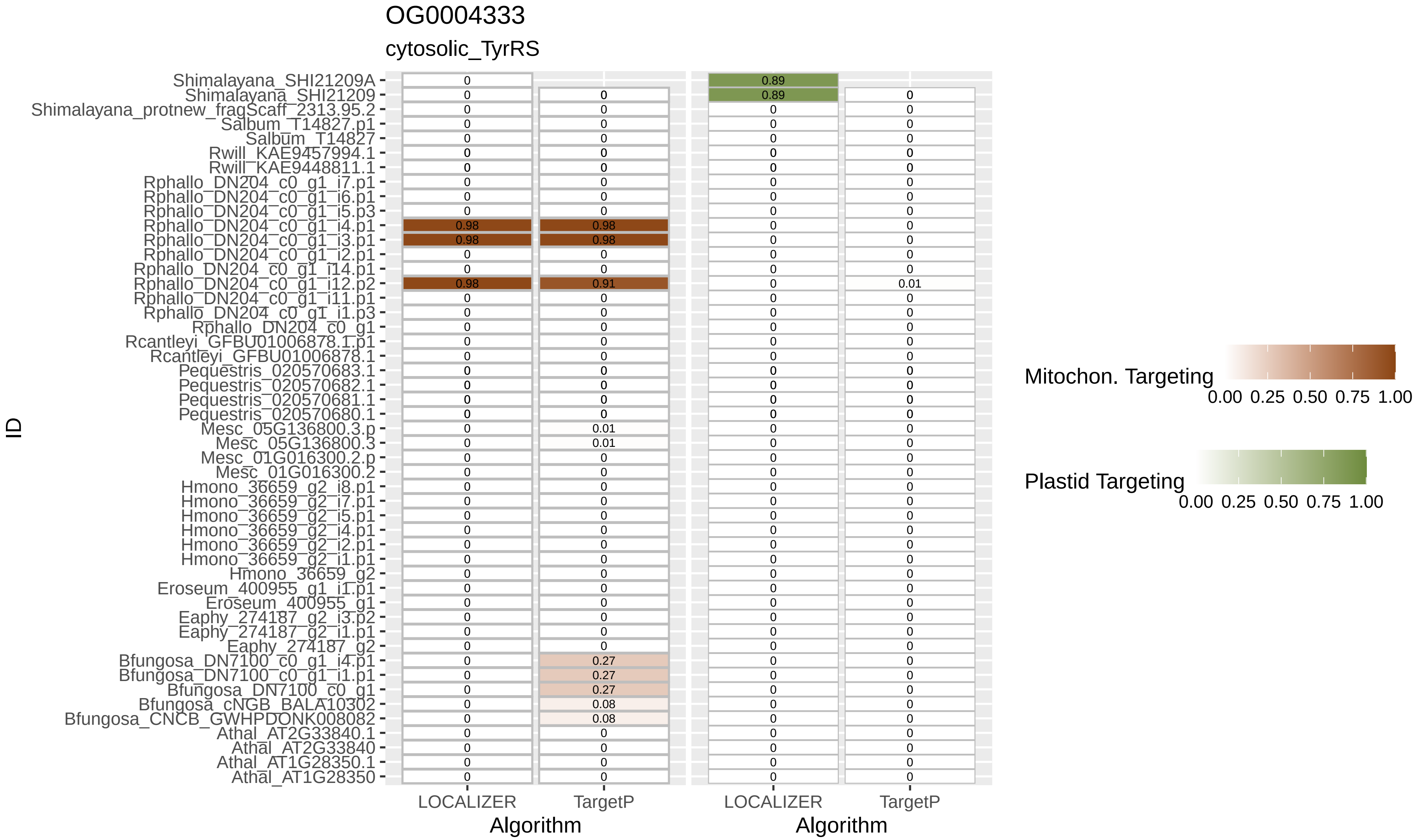
OG0004333
cytosolic_TyrRS
Shimalayana_SHI21209A
Shimalayana_SHI21209
Shimalayana_protnew_fragScaff_2313.95.2
Salbum_T14827.p1
Salbum_T14827
Rwill_KAE9457994.1
Rwill_KAE9448811.1
Rphallo_DN204_c0_g1_i7.p1
Rphallo_DN204_c0_g1_i6.p1
Rphallo_DN204_c0_g1_i5.p3
Rphallo_DN204_c0_g1_i4.p1
Rphallo_DN204_c0_g1_i3.p1
Rphallo_DN204_c0_g1_i2.p1
Rphallo_DN204_c0_g1_i14.p1
Rphallo_DN204_c0_g1_i12.p2
Rphallo_DN204_c0_g1_i11.p1
Rphallo_DN204_c0_g1_i1.p3
Rphallo_DN204_c0_g1
Rcantleyi_GFBU01006878.1.p1
Rcantleyi_GFBU01006878.1
Pequestris_020570683.1
Pequestris_020570682.1
Pequestris_020570681.1
Pequestris_020570680.1
Mesc_05G136800.3.p
Mesc_05G136800.3
Mesc_01G016300.2.p
Mesc_01G016300.2
Hmono_36659_g2_i8.p1
Hmono_36659_g2_i7.p1
Hmono_36659_g2_i5.p1
Hmono_36659_g2_i4.p1
Hmono_36659_g2_i2.p1
Hmono_36659_g2_i1.p1
Hmono_36659_g2
Eroseum_400955_g1_i1.p1
Eroseum_400955_g1
Eaphy_274187_g2_i3.p2
Eaphy_274187_g2_i1.p1
Eaphy_274187_g2
Bfungosa_DN7100_c0_g1_i4.p1
Bfungosa_DN7100_c0_g1_i1.p1
Bfungosa_DN7100_c0_g1
Bfungosa_cNGB_BALA10302
Bfungosa_CNCB_GWHPDONK008082
Athal_AT2G33840.1
Athal_AT2G33840
Athal_AT1G28350.1
Athal_AT1G28350
ID
LOCALIZER
TargetP
Algorithm
LOCALIZER
TargetP
Algorithm
Mitochon. Targeting
0.00 0.25 0.50 0.75 1.00
Plastid Targeting
0.00 0.25 0.50 0.75 1.00

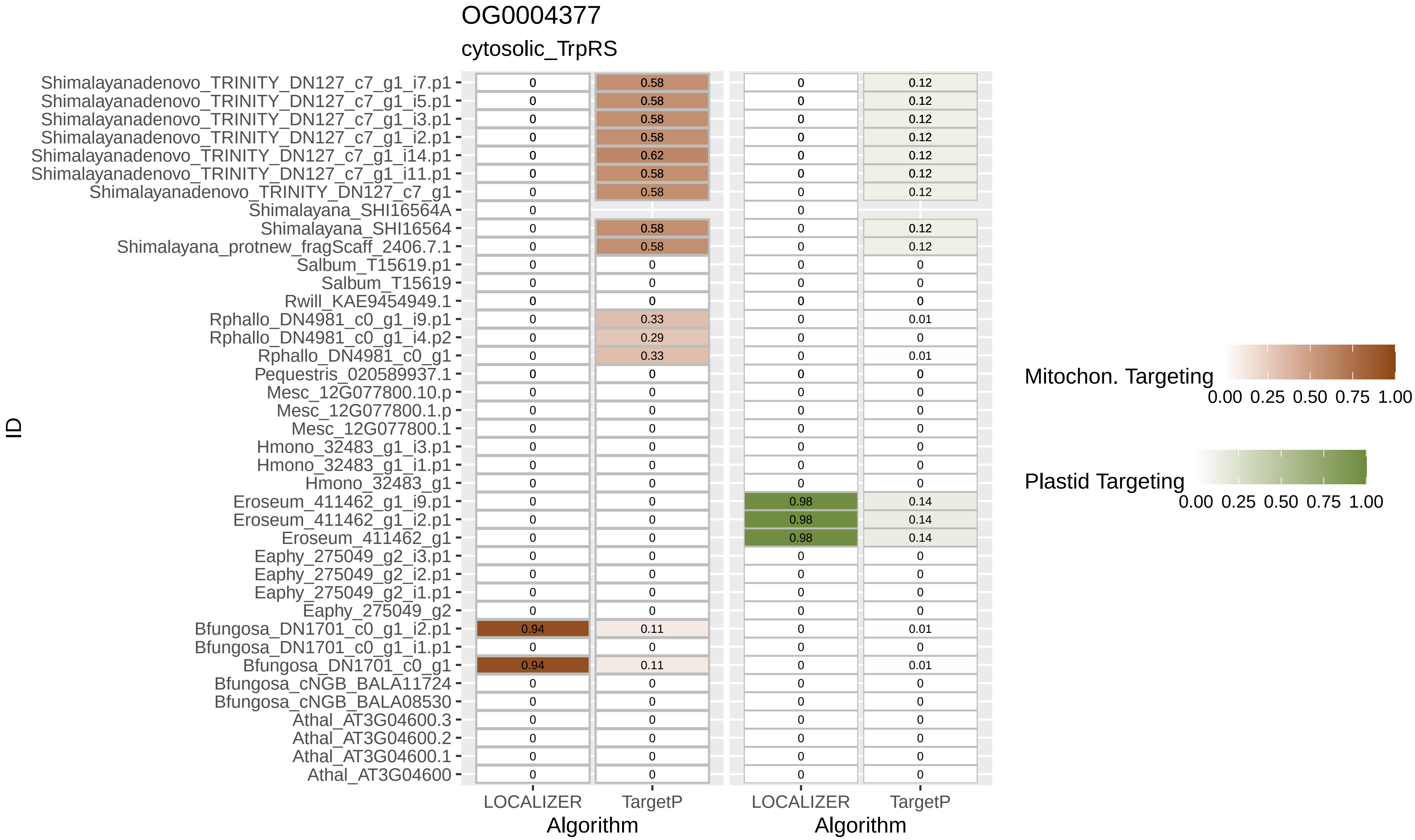

OG0004377
cytosolic_TrpRS
ID
Algorithm
LOCALIZER
TargetP
Mitochon. Targeting
Plastid Targeting
0.00 0.25 0.50 0.75 1.00
Shimalayanadenovo_TRINITY_DN127_c7_g1_i7.p1
Shimalayanadenovo_TRINITY_DN127_c7_g1_i5.p1
Shimalayanadenovo_TRINITY_DN127_c7_g1_i3.p1
Shimalayanadenovo_TRINITY_DN127_c7_g1_i2.p1
Shimalayanadenovo_TRINITY_DN127_c7_g1_i14.p1
Shimalayanadenovo_TRINITY_DN127_c7_g1_i11.p1
Shimalayanadenovo_TRINITY_DN127_c7_g1
Shimalayana_SHI16564A
Shimalayana_SHI16564
Shimalayana_protnew_fragScaff_2406.7.1
Salbum_T15619.p1
Salbum_T15619
Rwill_KAE9454949.1
Rphallo_DN4981_c0_g1_i9.p1
Rphallo_DN4981_c0_g1_i4.p2
Rphallo_DN4981_c0_g1
Pequestris_020589937.1
Mesc_12G077800.10.p
Mesc_12G077800.1.p
Mesc_12G077800.1
Hmono_32483_g1_i3.p1
Hmono_32483_g1_i1.p1
Hmono_32483_g1
Eroseum_411462_g1_i9.p1
Eroseum_411462_g1_i2.p1
Eroseum_411462_g1
Eaphy_275049_g2_i3.p1
Eaphy_275049_g2_i2.p1
Eaphy_275049_g2_i1.p1
Eaphy_275049_g2
Bfungosa_DN1701_c0_g1_i2.p1
Bfungosa_DN1701_c0_g1_i1.p1
Bfungosa_DN1701_c0_g1
Bfungosa_cNGB_BALA11724
Bfungosa_cNGB_BALA08530
Athal_AT3G04600.3
Athal_AT3G04600.2
Athal_AT3G04600.1
Athal_AT3G04600

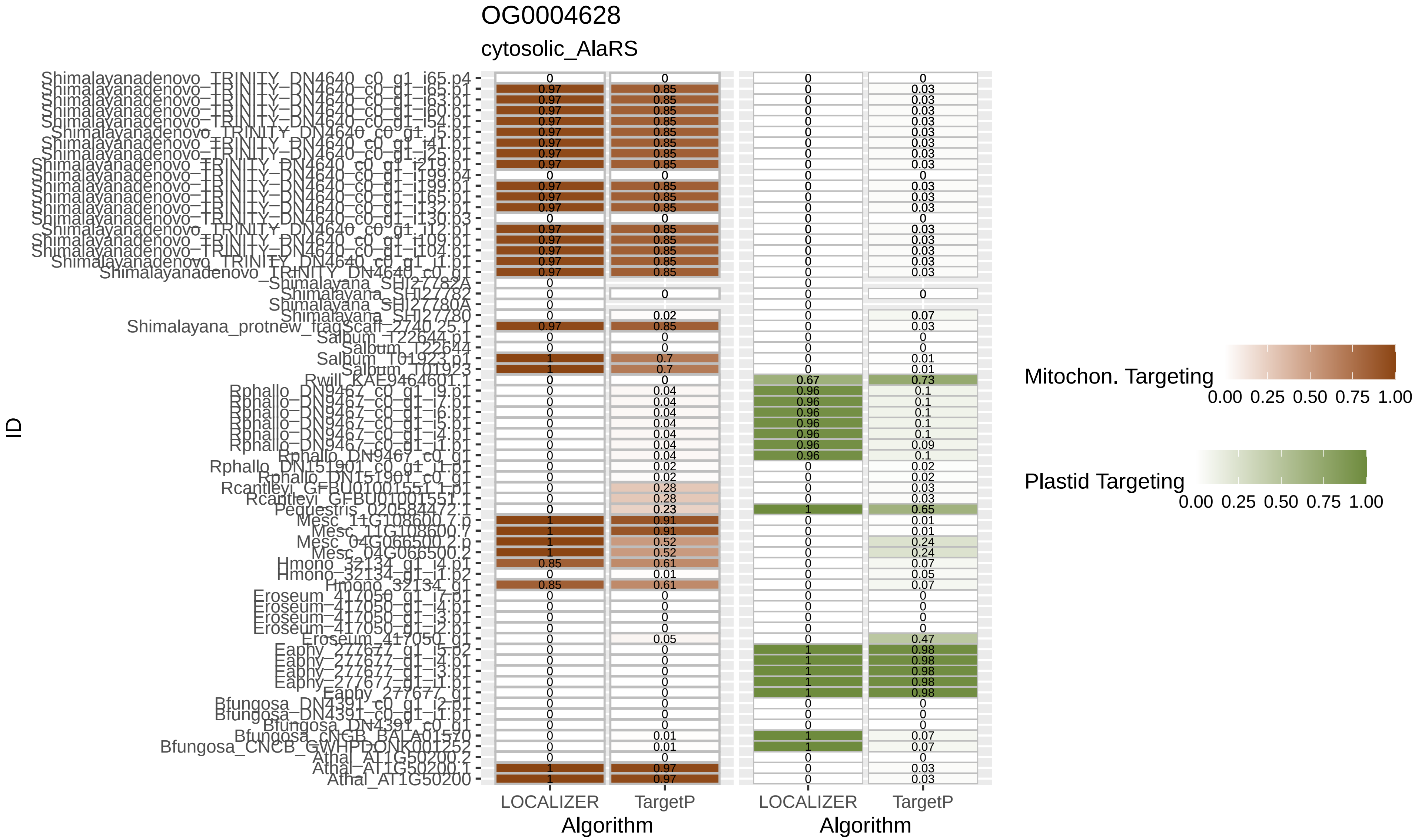

OG0004628
cytosolic_AlaRS
ID
LOCALIZER
TargetP
LOCALIZER
TargetP
Algorithm
Algorithm
Mitochon. Targeting
0.00 0.25 0.50 0.75 1.00
Plastid Targeting
0.00 0.25 0.50 0.75 1.00

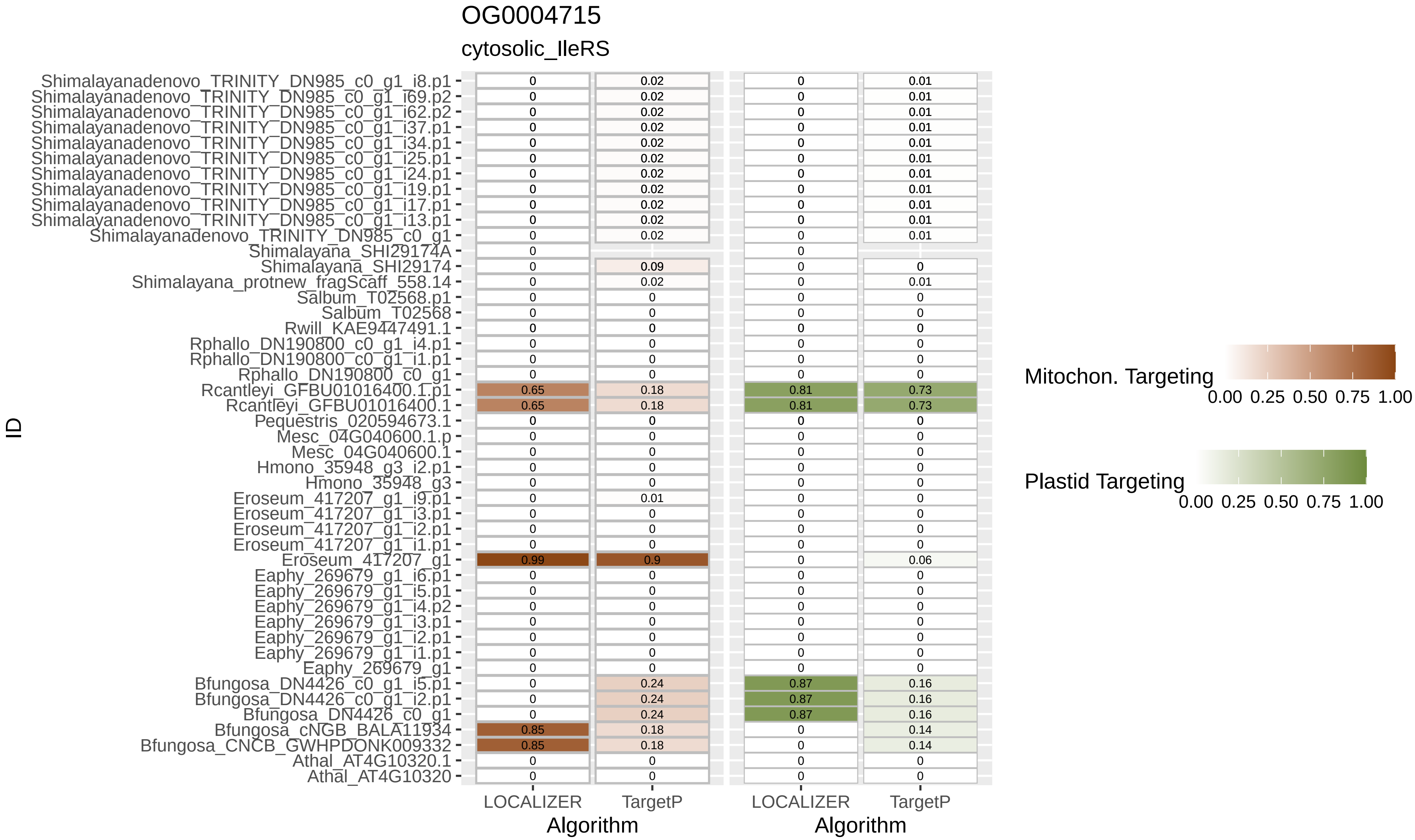
OG0004715
cytosolic_IleRS
ID
Shimalayanadenovo_TRINITY_DN985_c0_g1_i8.p1 0 0.02 0 0.01
Shimalayanadenovo_TRINITY_DN985_c0_g1_i69.p2 0 0.02 0 0.01
Shimalayanadenovo_TRINITY_DN985_c0_g1_i62.p2 0 0.02 0 0.01
Shimalayanadenovo_TRINITY_DN985_c0_g1_i37.p1 0 0.02 0 0.01
Shimalayanadenovo_TRINITY_DN985_c0_g1_i34.p1 0 0.02 0 0.01
Shimalayanadenovo_TRINITY_DN985_c0_g1_i25.p1 0 0.02 0 0.01
Shimalayanadenovo_TRINITY_DN985_c0_g1_i24.p1 0 0.02 0 0.01
Shimalayanadenovo_TRINITY_DN985_c0_g1_i19.p1 0 0.02 0 0.01
Shimalayanadenovo_TRINITY_DN985_c0_g1_i17.p1 0 0.02 0 0.01
Shimalayanadenovo_TRINITY_DN985_c0_g1_i13.p1 0 0.02 0 0.01
Shimalayanadenovo_TRINITY_DN985_c0_g1 0 0.02 0 0.01
Shimalayana_SHI29174A 0 0
Shimalayana_SHI29174 0 0.09 0 0
Shimalayana_protnew_fragScaff_558.14 0 0.02 0 0.01
Salbum_T02568.p1 0 0 0 0
Salbum_T02568 0 0 0 0
Rwill_KAE9447491.1 0 0 0 0
Rphallo_DN190800_c0_g1_i4.p1 0 0 0 0
Rphallo_DN190800_c0_g1_i1.p1 0 0 0 0
Rphallo_DN190800_c0_g1 0 0 0 0
Rcantleyi_GFBU01016400.1.p1 0.65 0.18 0.81 0.73
Rcantleyi_GFBU01016400.1 0.65 0.18 0.81 0.73
Pequestris_020594673.1 0 0 0 0
Mesc_04G040600.1.p 0 0 0 0
Mesc_04G040600.1 0 0 0 0
Hmono_35948_g3_i2.p1 0 0 0 0
Hmono_35948_g3 0 0 0 0
Eroseum_417207_g1_i9.p1 0 0.01 0 0
Eroseum_417207_g1_i3.p1 0 0 0 0
Eroseum_417207_g1_i2.p1 0 0 0 0
Eroseum_417207_g1_i1.p1 0 0 0 0
Eroseum_417207_g1 0.99 0.9 0 0.06
Eaphy_269679_g1_i6.p1 0 0 0 0
Eaphy_269679_g1_i5.p1 0 0 0 0
Eaphy_269679_g1_i4.p2 0 0 0 0
Eaphy_269679_g1_i3.p1 0 0 0 0
Eaphy_269679_g1_i2.p1 0 0 0 0
Eaphy_269679_g1_i1.p1 0 0 0 0
Eaphy_269679_g1 0 0 0 0
Bfungosa_DN4426_c0_g1_i5.p1 0 0.24 0.87 0.16
Bfungosa_DN4426_c0_g1_i2.p1 0 0.24 0.87 0.16
Bfungosa_DN4426_c0_g1 0 0.24 0.87 0.16
Bfungosa_cNGB_BALA11934 0.85 0.18 0 0.14
Bfungosa_CNCB_GWHPDONK009332 0.85 0.18 0 0.14
Athal_AT4G10320.1 0 0 0 0
Athal_AT4G10320 0 0 0 0
LOCALIZER
TargetP
Algorithm
LOCALIZER
TargetP
Algorithm
Mitochon. Targeting
0.00 0.25 0.50 0.75 1.00
Plastid Targeting
0.00 0.25 0.50 0.75 1.00
